# A novel reaction-diffusion architecture for engineering self-organized patterns in mammalian cells

**DOI:** 10.64898/2026.05.24.727552

**Authors:** Benjamin Swedlund, John J Danan, Ting-Xin Jiang, Soha Ben Tahar, Kyle Poon, Pranav S Bhamidipati, Zachary A Kreiger, Sandra Murillo, Minnal Kunnan, Daniel J G Pearce, Cheng-Ming Chuong, Ian M Ehrenreich, Leonardo Morsut

## Abstract

Reaction-diffusion circuits generate self-organized spatial patterns through local activation and long-range inhibition, but synthetic implementations in mammalian cells have been limited by the differential-diffusion requirement. Here, we introduce a novel architecture, juxtacrine activation with paracrine inhibition (JAPI), where the activator propagates through cell-cell contacts rather than diffusion. We demonstrate mathematically and numerically that JAPI accesses the same patterning regimes as classical diffusion-based circuits with one fewer free parameter. We then engineer compact synNotch-based JAPI circuits in mammalian fibroblasts and demonstrate their sufficiency for self-organized patterning through tunable, size-limited signal propagation. Functionalized to spatially control morphogen secretion, these circuits perturb feather bud formation on adjacent embryonic chicken epidermis. Finally, we develop a library-based approach to explore coupled, dual-JAPI circuits with tunable cross-inhibition, enabling programmable interactions between patterns and access to a broad morphospace of spatial states. Together, JAPI provides a compact, modular platform for programming self-organized multicellular patterning.

**Graphical Abstract:** 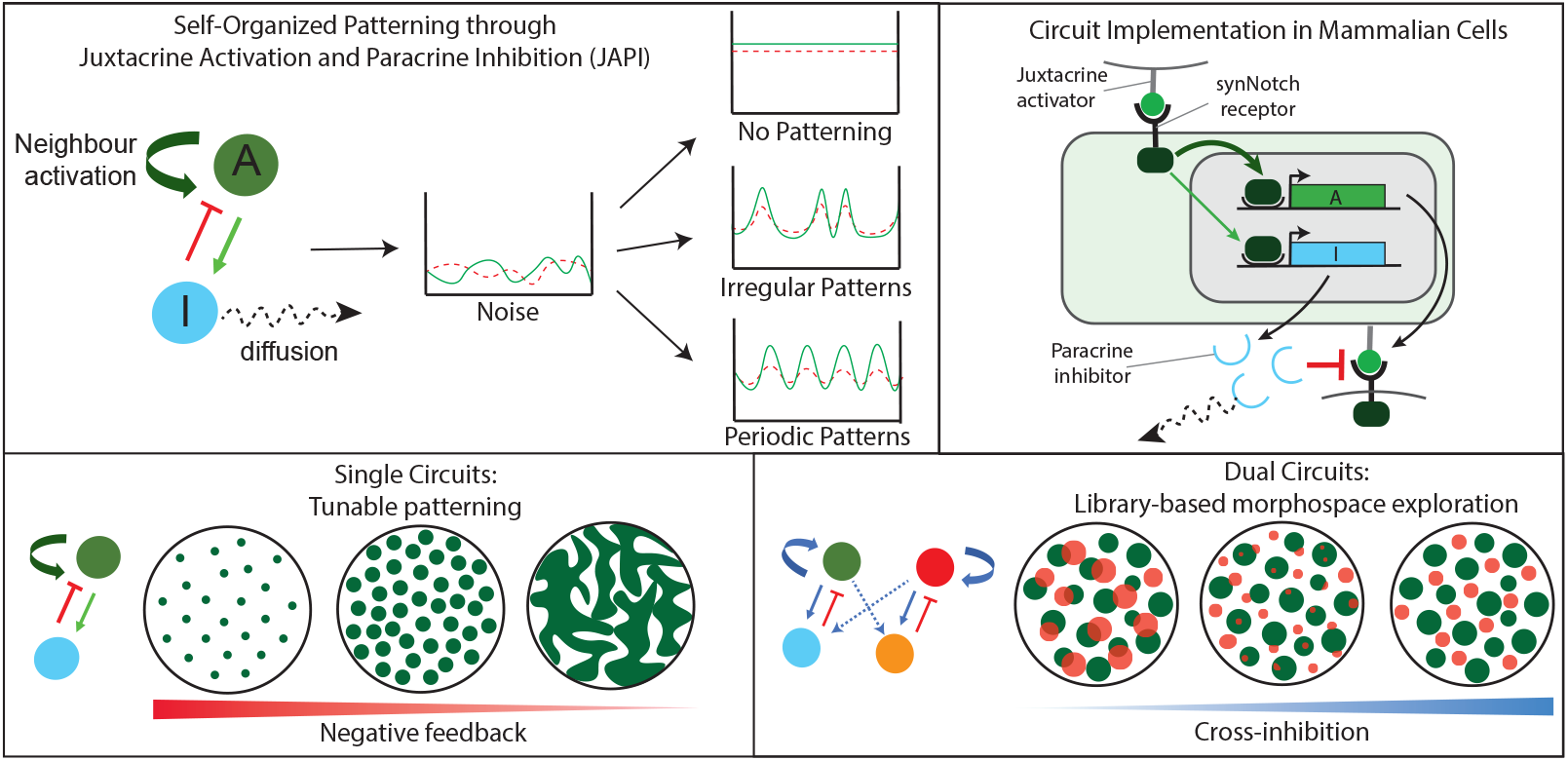

## Introduction

A central goal of synthetic biology is to program gene expression in space and time within multicellular systems, with applications to tissue engineering and the study of developmental processes (*1–3*). Of particular interest are genetic circuits capable of spontaneous symmetry breaking and formation of spatial patterns from a homogeneous field, without pre-patterned templates or externally imposed gradients (*4–7*). Reaction-diffusion circuits, composed of morphogens that interact through reaction and travel through diffusion, represent a paradigmatic class of such systems (*8–10*). In one of their most classical forms, a locally acting self-activating species and a long-range inhibitor interact to produce self-organized patterns from an initially uniform state (**Fig. 1A**, left). Natural variants of this logic have been shown to underlie the formation of feather buds, hair follicles, fingerprints, and left-right asymmetry, among other developmental patterns (*11–15*).

**Figure 1.**
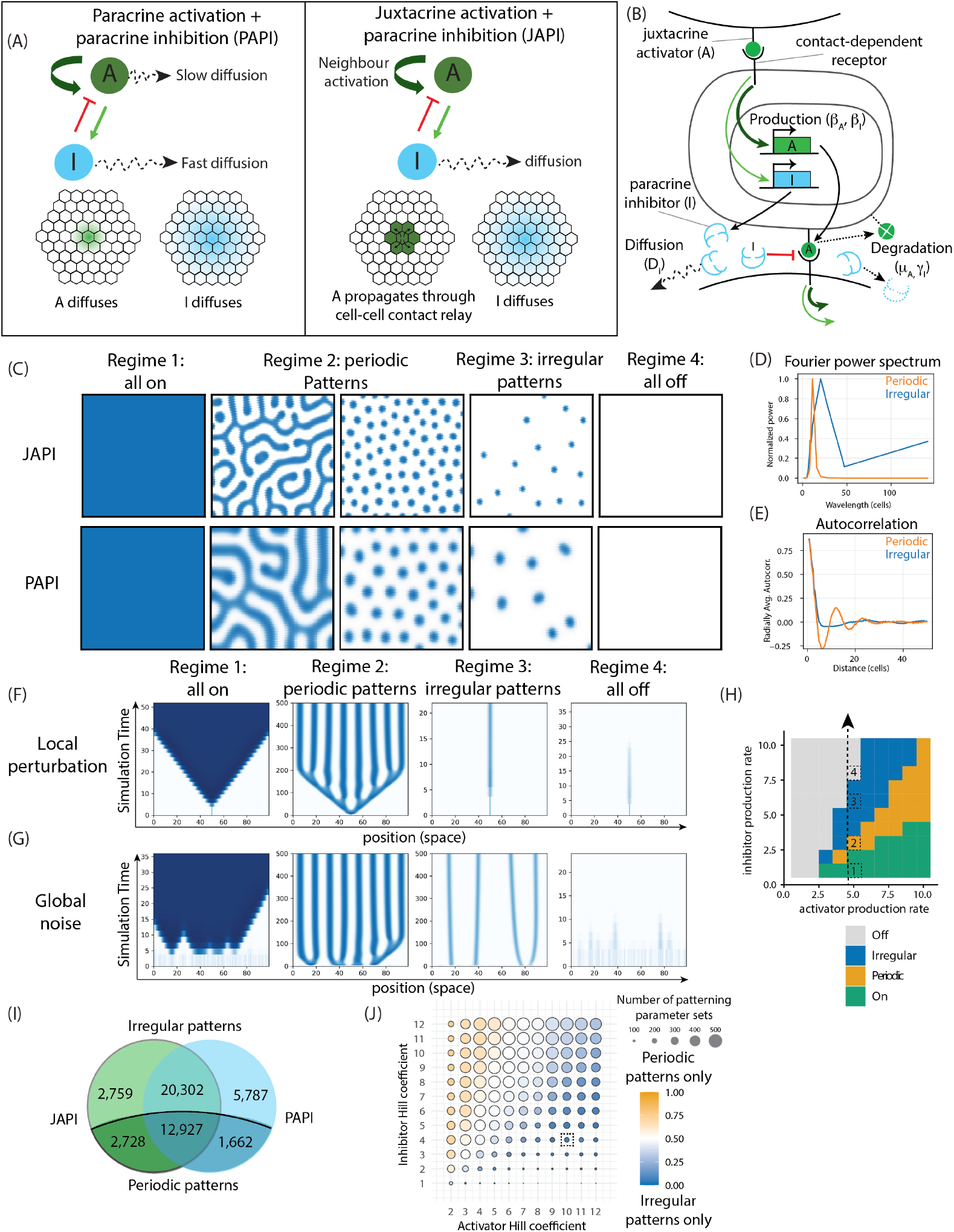
Juxtacrine activation with paracrine inhibition (JAPI) is a reaction-diffusion circuit architecture that accesses the same patterning regimes as dual-diffusion architectures without requiring differential diffusion. (**A**) Schematic design of two-species activator-inhibitor reaction-diffusion circuits comparing left, a dual-diffusion architecture (PAPI), and right, a mixed juxtacrine-paracrine alternative (JAPI). A stands for activator and I for inhibitor. The arrows in the circuit design represent the interactions between A and I: a dark green arrow for self-activation, a light green arrow for A activating I, and a red arrow for the inhibition of A by I. At the bottom is a cellular view of the movement of the species in a cellular lattice (black hexagons), with the activator represented in green, and the inhibitor in light blue, as above. For a JAPI circuit, the movement of A progresses through cell-cell contact-mediated juxtacrine signaling. (**B**) Close-up view of the JAPI architecture showing a possible molecular implementation of the regulatory steps in (A). The contact-dependent receptor detects the juxtacrine activator A (green) presented from the membrane of a neighboring cell. Inside the cell, the production of A and I downstream of receptor activation are shown with their respective production rates (b_a_, b_i_). The paracrine competitive inhibitor I (blue) is released into the extracellular space and shown diffusing away with a diffusion coefficient D_i_. Degradation of A and I (rates m_a_, g_i_) is shown close to the corresponding degraded species indicated as dotted shapes. (**C**) Endpoint snapshots of two-dimensional simulations for JAPI (top row) and PAPI (bottom row) circuits, for different values of the circuit parameters. Patterns of four qualitatively distinct outcomes are shown, labeled at the top. Blue intensity indicates dimensionless activator concentration (See supp Note 3 for description of dimensionless units). Simulation setup and parameter values per each simulation is found in Methods, Numerical Simulations. (**D-E**) Fourier power spectrum (D) and radially averaged autocorrelation (E) of endpoint patterns generated by two JAPI circuit simulations exhibiting qualitatively distinct spot regimes: irregular (blue) and periodic (orange), corresponding to the two ‘spots’ patterns shown in panel C (regime 2 periodic, right, and regime 3 irregular). (**F-G**) Kymographs of one-dimensional JAPI simulations from a centrally located activator burst (F) or from spatially uniform noise (G), for parameter sets representative of the four regimes labeled on top. The x axis is one-dimensional position; the y axis is simulation time progressing upward. Blue intensity indicates dimensionless activator concentration. Initial conditions and parameter values per each simulation is in Methods, Numerical Simulations. (**H**) Phase diagram of regime classification for a JAPI circuit as a function of activator production rate (x axis) and inhibitor production rate (y axis), with the remaining dimensionless parameters held constant (n_a_ = n_i_ = 3, γ = 0.5, D_i_ = 10). Tiles are colored by regime, as indicated in the legend. The four dashed boxes labeled 1 to 4 mark the parameter values used for the corresponding regimes in (F) and (G). Classification rules are found in Methods, Defining Patterning Regimes. (**I**) Venn diagram showing the distribution of the ∼46,000 parameter sets of the 5 dimensionless parameters that give rise to patterns in JAPI and PAPI parameter sweeps. Numbers in each region indicate parameter set counts and overlap producing irregular patterns (top) and periodic patterns (bottom), for JAPI (left) and PAPI (right). Sweep design in Methods, Defining Patterning Regimes. (**J**) Dot plot summarizing patterning outcomes from a parameter sweep of a JAPI circuit across ∼1,900 parameter combinations for each activator Hill coefficient (x axis) and inhibitor Hill coefficient (y axis) pair. Parameter combinations were generated by independently varying three non-dimensionalized parameters: activator and inhibitor production rates, and inhibitor degradation rate. For each (n_a_, n_i_) pair, the dot size is proportional to the number of parameter combinations producing patterns; dot color indicates the proportion of those producing periodic patterns (orange) versus irregular patterns (blue). The dashed box marks the (n_a_, n_i_) pair of a parametrized synNotch circuit in Fig. 2. Sweep ranges and pattern definitions in Methods, Defining Patterning Regimes.

Reaction-diffusion circuits are therefore an attractive target for synthetic reconstruction, and multiple efforts have sought to reconstitute their behavior using engineered signaling circuits (*16–18*). An ideal implementation would be compact, orthogonal, modular, and controllable, enabling exploration of theoretical design principles, integration with natural patterning systems, and expansion to multi-circuit networks capable of generating increasingly complex spatial patterns. Classical reaction-diffusion architectures, however, impose a stringent constraint: the inhibitor must diffuse substantially faster than the activator (*8, 9, 19*), a requirement that is difficult to satisfy with engineered components in mammalian cells. Existing reconstruction approaches have circumvented this either by harnessing stochasticity as a force in pattern formation (*16*), engineering larger, three-component circuits in Prokaryotes (*17*), or co-opting naturally differentially diffusing signaling pairs in mammalian cells (*18*) - but each relaxes one or more of the four ideal properties defined above. No existing approach combines compactness, orthogonality, modularity, and tunability in mammalian cells.

One signaling modality that has been largely overlooked in the context of reaction-diffusion is juxtacrine communication, in which cells signal exclusively through direct cell-cell contacts. Although juxtracrine signaling does not involve spatial travel through diffusion, it participates in multicellular patterning through mechanisms such as lateral inhibition, which enables spontaneous generation of single-cell-scale checkerboard patterns, and lateral induction, which propagates pre-existing patterns at larger spatial scales (*20–22*). More recently, a synthetic version of juxtacrine signaling has been introduced: Synthetic Notch (synNotch) receptors enable compact, orthogonal engineering of juxtacrine signaling in mammalian cells through custom ligand recognition and transcriptional responses (*23*). These receptors have been used to program lateral inhibition (*24*), lateral induction (*25*) and, more recently expanded to work in a paracrine manner to generate pre-patterned morphogen gradients (*26*). Together, these applications establish that synNotch can provide both juxtacrine and paracrine signaling as compact, orthogonal primitives. Whether a reaction-diffusion architecture can be built from synNotch parts, and whether it can support multicellular self-organized patterning in a compact, modular, and multiplexable form, has not yet been explored.

Here we develop and implement a novel reaction-diffusion architecture, which we name juxtacrine activation with paracrine inhibition (JAPI). In JAPI, the activator cannot diffuse freely but can propagate through direct cell-cell contact relay, replacing the paracrine auto-activator of classical systems. We first demonstrate analytically and numerically that this substitution removes the differential-diffusion requirement while accessing the same patterning regimes as dual-diffusion architectures. We then engineer compact synNotch-based JAPI circuits in mammalian fibroblasts and demonstrate their sufficiency for producing self-organized non-periodic patterns *in vitro*. These patterns emerge from stochastic nucleation, juxtacrine relay, and inhibitor-mediated arrest, with domain size tunable through circuit parameters. We further functionalize these circuits to drive patterned expression of endogenous signaling factors, allowing them to interface with natural patterning processes such as feather bud formation in embryonic chicken skin. Finally, we show that the modular architecture of JAPI supports multiplexing: two orthogonal circuits operating in parallel produce spatially independent patterns. The cross-correlation between the two patterning circuits can be tuned positively through initial conditions or negatively through engineered cross-inhibition, generating a broad, tunable dual-patterning morphospace. Together, these results establish JAPI as a compact, orthogonal platform for synthetic self-organized patterning in mammalian cells, define a new entry in a broader architecture-level design space, and open a path toward compositional patterning networks and synthetic interfaces with developmental processes.

## Results

### Juxtacrine activation with paracrine inhibition (JAPI) is a functional reaction-diffusion circuit architecture that does not require differential diffusion

We propose a novel two-component reaction-diffusion circuit design in which a diffusible inhibitor is combined with a membrane-tethered activator that propagates through cell-cell contacts rather than diffusion (**Fig. 1A**). In these juxtacrine activator-paracrine inhibitor (JAPI) circuits, the spatial mobility of the activator arises through juxtacrine activation of neighboring cells, an active process, rather than through paracrine diffusion, the passive equilibration of concentrations. Assuming a common reaction function for activator and inhibitor production, these JAPI circuits can be described by the discrete pair of equations [**Eq. 1-2**] and compared with the classical dual-diffusion paracrine activator paracrine inhibitor (PAPI) system [**Eq. 3-4**], where *K a* denotes the kernel-weighted activator input from neighboring cells (see **Supp. Note 1**).

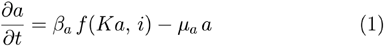

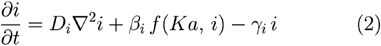

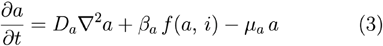

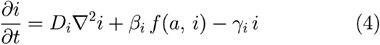

The network topology of an activator activating its own production and the production of the inhibitor is shared between the two systems, as well as the parameters for production rates, degradation rates, and the diffusion coefficient of the inhibitor. The key architectural substitution is that the discrete cell-to-cell coupling described by K replaces the activator diffusion coefficient D_a_, which results in one less parameter, and therefore one less degree of freedom in the JAPI architecture (**Fig. 1B**). This mixed discrete-continuous architecture preserves the network topology but is mechanistically and mathematically distinct from classical PAPI systems.

We first assessed whether this substitution in the activator mode of transport affects the potential for Turing instabilities to arise. To answer this, we formulated both architectures on a 1D cellular lattice and performed linear stability analysis (LSA) around the non-trivial (on) steady state (**Supp. Note 1, Eq. 6-12**). Decomposing perturbations into Fourier modes at each wavenumber q reduces the linearized dynamics to a 2×2 eigenvalue problem at each q: the linearized system is described by a 2×2 matrix A(q) (a wavenumber-dependent Jacobian) whose eigenvalues give the growth rate of perturbations at that wavenumber. The matrices A(q) for JAPI and PAPI differ in their entries [**Eq. 5-6**] (**Supp. Note 1, Eq. 13-24**).

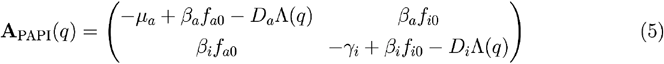

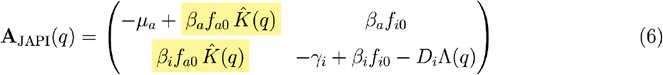

The architectural differences translate directly into the linear operators. In PAPI, spatial dependence (q) enters only through diffusive penalties on the diagonal entries. In JAPI, two structural differences arise: first, the activator-to-inhibitor cross-coupling acquires a kernel factor 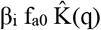, because inhibitor production is driven by the same juxtacrine input. Second, and more importantly, the activator diagonal carries 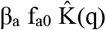 instead of −D_a_ Λ(q), entangling the spatial term with the local self-amplification: this shapes both the instability mechanism and the parameter space available for patterning, as we discuss below.

The diagonal entries of the operator are particularly significant for stability: we call them P(q) (activator self-coupling) and Q(q) (inhibitor self-coupling). Turing instability requires P(q) > 0 at intermediate wavenumbers combined with Q(q) < 0. In PAPI, it is well established that this condition can be met when D_i_ ≫ D_a_ (*8, 9*). The inhibitor self-coupling Q(q) is identical in both architectures and always negative, but since the activator self-coupling P(q) differs between PAPI and JAPI, whether the Turing instability condition can be met in JAPI, where D_a_ is absent, is not obvious. As we prove in **Supplementary Note 1 Eq. 30-39**, the JAPI operator admits Turing-type finite-wavenumber instabilities under the same local activator–inhibitor prerequisites familiar from PAPI: activator self-amplification combined with stability of the activated steady state to uniform perturbations. The proof rests on a distinctive JAPI mechanism: mode selection is provided by the kernel geometry through 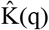, rather than by a ratio of diffusion coefficients. Long-wavelength modes inherit strong juxtacrine coupling 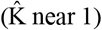 and remain self-amplifying; combined with the q-dependent inhibitor diffusion penalty, this opens a finite wavenumber instability without any activator diffusion. The proof is constructive: all modes are stable at D_i_ = 0 and a destabilizing mode is exhibited explicitly. JAPI therefore realizes the same class of predicted finite-wavenumber regimes as PAPI in a parameter-compact form, without requiring an independently diffusing activator species.

What allows the activator to have a spatial reach without diffusion? In JAPI, the activator’s spatial reach is characterized by an effective propagation velocity v_a_, set by the time required for a newly stimulated cell to produce sufficient activator ligand to drive its next neighbor (a cycle rate-limited by transcription, translation, and surface ligand presentation, see **Supp. Note 2**). Two biological consequences follow from this substitution of relay for diffusion. First, the activator front advances discretely, one cell at a time, rather than as a continuous spatial profile; therefore, JAPI’s spatial coupling is intrinsically cellular, not molecular. Second, v_a_ is tunable through interventions on promoter strength, Hill cooperativity, and protein half-life, rather than through molecular engineering of diffusion properties, which makes JAPI accessible to implementation with mammalian gene circuits, removing a key protein-engineering constraint (**Supp. Note 2, Fig 1B**). The mathematical analysis above and the physical interpretation here together establish that JAPI is, in principle, a viable reaction-diffusion architecture for self-organized patterning.

To assess whether it is viable in practice, we turn to numerical simulation to explore how much of parameter space supports patterning, what the resulting patterns look like, and how they compare to dual-diffusion implementations. To numerically simulate both architectures on 2D hexagonal cellular lattices, we non-dimensionalized the system (**Supp. Note 3**) and we specified the production function as a two-input Hill function representing competitive inhibition, shared between activator and inhibitor (**Eq.7**), a standard choice for activator-inhibitor reaction-diffusion circuits with saturating transcriptional responses (*27, 28*).

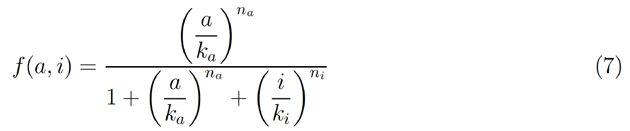

Starting from stochastic initial conditions, simulations recovered the three LSA-predicted regimes (uniform activation, uniform quiescence, periodic Turing patterns) in both architectures (**Fig. 1C**). Both circuits additionally produced a fourth regime, irregular non-periodic patterns, in parameter regions where LSA predicts homogeneous stability; these patterns, previously described in reaction-diffusion systems, arise from finite-amplitude perturbations in the linearly stable regime (*29, 30*). The two non-homogeneous regimes are distinguished by their spatial signatures: Turing patterns show a sharp Fourier peak and oscillatory autocorrelation; irregular patterns show a broad Fourier spectrum and monotonically decaying autocorrelation (**Fig. 1D-E**). In all the non-homogeneous regimes, one qualitative difference between architectures is directly visible: JAPI patterns show sharper, near-single-cell-width activator domain boundaries, while PAPI patterns show graded transitions on a length scale set by D_a_ (**Fig. 1C**). This difference is specific to the activator; in contrast, the inhibitor, diffusible in both architectures, exhibits a similar spatial profile in the two architectures (**Fig. S1A**).

Having identified the four regimes for both the architectures in targeted 2D simulations, we next compared their distribution across parameter space. We performed high-throughput 1D simulations for both architectures over a uniformly-spaced grid of more than 250,000 parameter combinations, classifying the outcome of each according to the propagation behavior of a centrally seeded activator perturbation: in parameter sets allowing for Turing instabilities, the perturbation propagates across the field and resolves into a periodic pattern, while in irregular parameter sets it expands into a single size-limited activated domain (**Fig. 1F, S1B**; see Methods, ‘Defining Patterning Regimes’). Simulations from initial stochastic noise confirmed the validity of this classification method (**Fig. 1G, S1C**). Across a larger parameter set, we observed that, for both architectures JAPI and PAPI, when the four regimes are present in a two-dimensional parameter projection, they are traversable in a defined order: increasing the inhibitor production rate β_i_ drives the system through uniform activation, periodic patterns, irregular patterns, and uniform quiescence sequentially (**Fig. 1H, S1D**). Globally, the number of parameter combinations producing each non-homogeneous regime was comparable between architectures (**Fig. 1I**), establishing global functional equivalence between JAPI and PAPI at the level of regime distribution.

We performed a focused screen on two of the dimensionless groups governing JAPI patterning, the Hill coefficients n_a_ and n_i_, which set the steepness and threshold of the reaction function (**Fig. S1E**). These are particularly interesting: in PAPI circuits with equal Hill coefficients (n_a_ = n_i_), increased cooperativity has been shown to expand the Turing-unstable parameter space and relax the differential-diffusion requirement (*28*). In JAPI, n_a_ additionally affects the activator propagation velocity v_a_ without the D_a_-mediated decoupling available in PAPI (see also Supp. Note 2). We therefore asked whether sweeping n_a_ and n_i_ independently reveals architecture-specific Hill-coefficient signatures. The independent sweep uncovered a clear sorting of patterning regimes by the relative cooperativity of activation and inhibition, which is of interest in itself: increasing n_a_ relative to n_i_ shifts the dominant regime from periodic toward irregular patterns, while the opposite ordering (n_i_ > n_a_) favors periodic patterns (**Fig. 1J**). Notably, this sorting was conserved in both JAPI and PAPI architectures (**Fig. S1F, S1G-H**). The qualitative regime sorting is therefore dominated by the shared local-kinetics route, and the architectural difference in how n_a_ enters v_a_ does not produce architecture-specific patterning differences through this route.

Together, these results establish JAPI as a compact reaction-diffusion architecture that accesses the same class of patterning regimes with a similar efficiency as the dual-diffusion PAPI implementation, without requiring differential diffusion.

### A synNotch-based JAPI circuit produces self-organized non-periodic patterns in mammalian cells

Given the potential of this novel JAPI architecture to give rise to emergent patterning phenotypes, we designed an experimental implementation for mammalian cells using synthetic, orthogonal cell-cell communication circuits (**Fig. 2A**). The architecture requires two components: a contact-dependent activator that self-amplifies through juxtacrine signaling, and a soluble inhibitor whose production is driven by the same activation event. We reasoned that synNotch receptor-based circuits (*23*) could provide both functions: a synNotch receptor could be configured to activate transcription of its own membrane-tethered ligand, encoding the contact-dependent auto-activation logic; if the same activation event also drove the production of a secreted competitive inhibitor, the full JAPI logic would be reconstructed. We first validated that each branch operates as required in isolation in L929 mouse fibroblast cells. The self-activating juxtacrine relay implemented with the anti-GFP synNotch driving its own GFP ligand has been characterized previously in isolation (*25*). We confirmed that, in clonal cell lines containing the juxtacrine relay circuit, stochastic activation events nucleate locally and propagate as expanding waves of GFP positive cells without any external trigger, providing the auto-activator branch of JAPI (**Fig. S2A-B**). For the inhibitory branch, we demonstrated that a secreted anti-GFP nanobody dimer that blocks ligand binding to the receptor (*26*) inhibited the juxtacrine relay (**Fig. S2C-D**). Additionally, when provided locally from a cluster of inhibitor-secreting cells, inhibition decayed with distance from the source, indicating local action with a spatial reach consistent with diffusion (**Fig. S2E-G**). Both components therefore behave as required for the JAPI architecture: local juxtacrine auto-activation and diffusible paracrine inhibition.

**Figure 2.**
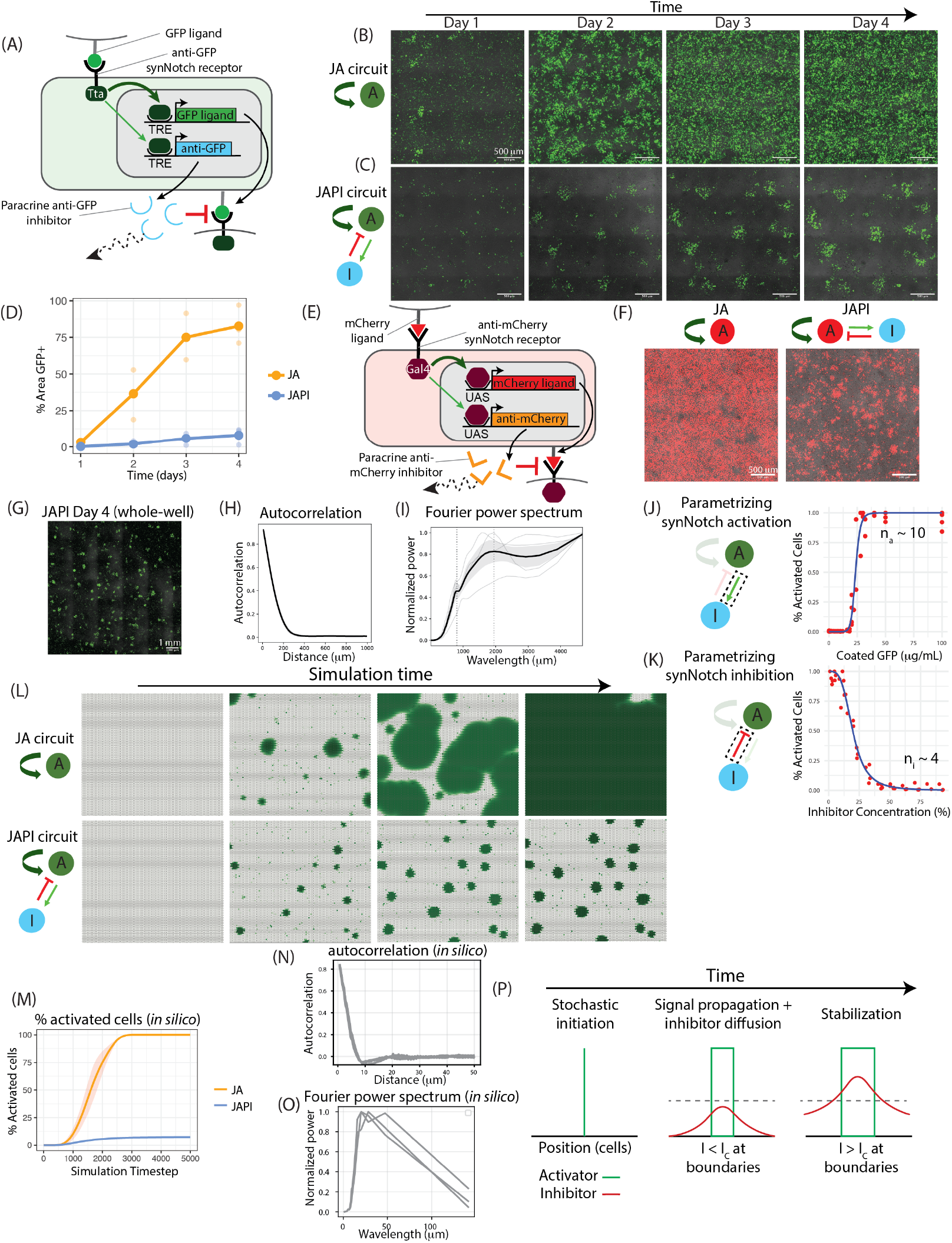
A synNotch-based JAPI circuit produces self-organized non-periodic patterns in mammalian cells through inhibitor-mediated arrest. (**A**) Schematic of a synNotch-based JAPI circuit implementation in mammalian cells. A neighboring cell presents a juxtacrine activator, GFP ligand, on its membrane; the anti-GFP synNotch receptor with an intracellular Tta transactivation domain (dark green) detects this ligand and activates transcription from TRE promoters of two downstream effector genes: one encoding the membrane-tethered GFP ligand (juxtacrine activator, A, green), another one encoding a diffusible anti-GFP nanobody dimer (paracrine inhibitor, I, light blue) that competitively blocks ligand-receptor binding. (**B-C**) Fluorescence microscope images at the indicated timepoints of a timelapse experiment, showing the same field of view across time. Green indicates activated cells (GFP signal); brightfield in grey. (B), L929 cells containing a juxtacrine self-activation (JA) circuit without inhibition; (C), L929 cells containing a complete JAPI circuit with activation and inhibition as indicated on the left schematic. In both cases, the experiment is initiated from homogeneous inactivated cells. Scale bar = 500 um. (**D**) Dot-plot graph of area fraction occupied by GFP signal over time, for cells containing a JA circuit (orange) or a JAPI circuit (blue). Each dot is an independent technical replicate, from a clonal population of JA and JAPI cells (n = 3); the lines connect replicate means. For the quantification pipeline, see Methods, Image Analysis. (**E**) Schematic of a synNotch-based JAPI circuit with the same architecture as in (A) but built from an orthogonal molecular trio: an anti-mCherry synNotch receptor with an intracellular Gal4 transactivation domain (dark red) detects juxtacrine ligand mCherry on a neighboring cell and drives transcription from UAS promoters of two downstream cassettes: one encoding the membrane-tethered mCherry ligand (juxtacrine activator, red), another encoding a diffusible anti-mCherry nanobody dimer (paracrine inhibitor, orange) that competitively blocks ligand-receptor binding. (**F**) Fluorescence microscope endpoint images of mCherry-based JA (left) and JAPI (right) circuits on day 4. Red indicates activated cells (mCherry signal); brightfield in grey. Initial condition: homogeneous inactivated cell lawn. Scale bar = 500 µm. (**G**) Whole-well fluorescence microscope endpoint image at day 4 of a clonal cell line containing the GFP-based JAPI circuit from (A). Green indicates activated cells (GFP signal); brightfield in grey. Initial condition: homogeneous inactivated cell lawn. Scale bar = 500 µm. (**H**) Line plot of the radially averaged autocorrelation of processed GFP signal, computed from images such as (G). Light grey curves are individual experiments, black curve is the average, grey shading is standard deviation (n = 4). Image processing in Methods, Image Analysis. (**I**) Line plot of the Fourier power spectrum of processed GFP signal, computed from images such as (G). Light grey curves are individual experiments, black curve is the average, grey shading is standard deviation (n = 4). Dotted vertical lines mark the identified local maxima. Image processing in Methods, Image Analysis. (**J**) Left, schematic of the JAPI circuit with the activator production branch highlighted by a dashed box. Right, dose-response curve of the proportion of synNotch activated cells at 48h (red dots, individual experimental data points) as a function of the concentration of plate-bound GFP ligand presented to the cells (x axis, in µg/mL), fitted to an activating Hill function (blue curve), from n = 5 independent replicates. The fitted Hill coefficient n_a_ is indicated in the figure. Experimental setup and fitting procedure in Methods, synNotch parametrization experiments. (**K**) Left, schematic of the JAPI circuit with the inhibition branch highlighted by a dashed box. Right, dose-response curve of the proportion of synNotch activated cells at 48h (red dots, individual experimental data points) as a function of the concentration of soluble anti-GFP inhibitor provided in the presence of a constant concentration of plate-bound GFP ligand (x axis, in % relative dilution of inhibitor-containing conditioned media), fitted to an inhibitory Hill function (blue curve), from n = 5 independent replicates. The fitted Hill coefficient n_i_ is indicated in the figure. (**L**) Rendering of spatial activator distribution on a 2D cell lattice at progressive timepoints of a simulation, showing the entire simulation field across time. Green intensity indicates dimensionless activator concentration. JA circuit (top row) and JAPI circuit (bottom row), parametrized with the Hill coefficients measured in (J) and (K) (n_a_ = 10, n_i_ = 4), initiated from an inactivated state with continuous stochastic activation bursts. Simulation setup and complete parameter values per each simulation is in Methods, Numerical Simulations. (**M**) Line plot of percentage of activated cells over simulation time, for the JA circuit (orange) and JAPI circuit (blue) simulations like the ones in (L). Mean curves shown with shading for standard deviation across three simulated replicates. (**N**) Line plot of the radially averaged autocorrelation of the simulated activator field, computed from JAPI simulations as in (L) at the simulation endpoint of three replicated simulations. Simulation result processing in Methods, Spatial analysis of 2D simulations. Compare to (H), which shows the same metric computed from experimental images. (**O**) Line plot of the Fourier power spectrum of the simulated activator field, computed from JAPI simulations as in (L) at the simulation endpoint. The results from three replicated simulations are shown. Compare to (I), which shows the same metric computed from experimental images. Data processing in Methods, Spatial analysis of 2D simulations. (**P**) Schematic representation of the inhibitor-mediated arrest mechanism, shown as three successive stages labeled at the top. Left, stochastic initiation: a single cell crosses the activation threshold and begins producing the activator. Middle, signal propagation and inhibitor diffusion: the activated domain expands through juxtacrine relay while accumulating inhibitor through paracrine diffusion, with inhibitor concentration at the boundary of the green domain falling below the critical threshold I_C_. Right, stabilization: as the domain expands, inhibitor concentration at the boundaries exceeds I_C_, stalling further activation and arresting the domain at a finite size. Activator concentration is shown as on/off active/inactive; inhibitor concentration is shown as a graded red curve; the threshold I_C_ is indicated as a dashed horizontal line. Mechanism derivation in Supp. Note 4.

We then assembled the full JAPI architecture by combining the validated parts in a single cell line: an anti-GFP synNotch receptor driving expression of both GFP ligand (juxtacrine activation) and a secreted anti-GFP nanobody dimer (paracrine inhibition) (**Fig. 2A**). We compared the full JAPI circuit to a control circuit lacking the inhibitor branch (juxtacrine activator only, JA). Cells containing the JA circuit showed spontaneous local activation events that propagated outward as expanding waves until the entire plate was activated (**Fig. 2B**,**D**). In contrast, clonal cell lines containing the full JAPI circuit showed local activation that resolved into discrete, finite-sized stable domains (**Fig. 2C-D, S3A**, Supplemental Movie 1). Well-defined activation domains emerged from genetically homogeneous populations grown directly into 96-well plates from single FACS-isolated cells, showing that patterning and symmetry breaking do not require pre-existing cellular nor genetic heterogeneity, but instead arise as intrinsic properties of the circuit (**Fig. S3B-D**). Pattern formation was not exclusive to the anti-GFP synNotch system: an analogous JAPI circuit built around anti-mCherry synNotch, mCherry ligand, and a paracrine anti-mCherry nanobody dimer (*26*) also produced stable patterns (**Fig. 2E-F**), indicating that patterning reflects the JAPI architecture rather than properties of any specific molecular pair.

Several features of the observed patterning mechanism matched the conceptual setup of the JAPI architecture as described in Figure 1. We first confirmed that the resulting patterns of ligand GFP expression are a direct reflection of synNotch receptor activity (**Fig. S4A-B**), and that all cells maintain the ability to activate from plate-bound ligand presentation, ruling out transgene silencing as being implicated in patterning (**Fig. S4C**). Secondly, the activated and inhibited states of JAPI cells are both reversible and dynamically maintained, as shown by the spontaneous re-emergence of patterns from FACS-isolated homogeneous populations of activated and inhibited cells separately (**Fig. S4D-F**). Finally, when juxtaposing locally seeded growing domains of JAPI cells next to JA cells lacking the inhibitor branch, JAPI cells generated stable spatial patterns, while neighboring JA cells propagated unrestricted, supporting the notion that inhibitor action is mediated locally through gradients of concentration, instead of acting at the well-scale (**Fig. S4G-I**). Together, these results support the notion that our experimental system represents a faithful implementation of the JAPI logic.

We next sought to determine which patterning regime this implementation occupies. As described below, three lines of evidence place it within the irregular regime: (1) the spatial features of the patterns, (2) a perturbative test of initial condition sensitivity, and (3) a parametric mapping based on experimentally measured Hill coefficients.

First, we characterized the spatial structure of the resulting JAPI patterns at the culture well-scale (**Fig. 2G**). The radially averaged autocorrelation of the activated signal decayed monotonically with distance, without the oscillatory structure that would indicate periodicity (**Fig. 2H**). The Fourier power spectrum was broad and lacked a sharp peak characteristic of wavelength selection in Turing patterning (**Fig. 2I**). We noted the Fourier spectrum did show non-monotonic structure with characteristic length scales near 800 and 2000 µm absent in the JA control during signal propagation (**Fig. 2I, S5A-B**), indicating that in this system the inhibitor introduces specific spatial scales into the activated distribution even in the absence of wavelength selection, though not the dominant wavelength characteristic of periodic patterning. Moreover, activated domains showed a broad size distribution centered around a mean equivalent diameter of approximately 150 µm (**Fig. S5C**), in contrast to the uniform feature size Turing patterning produces through wavelength selection. The same non-Turing signatures (monotonic autocorrelation and broad Fourier spectrum without a sharp peak) were observed in the anti-mCherry-based circuit (**Fig. S5D-E**).

Second, the irregular regime makes a distinct dynamical prediction: because irregular patterns require finite-amplitude perturbations against a stable homogeneous background, they should be sensitive to initial conditions in a way Turing patterns are not. In 1D simulations, irregular patterns formed only over a bounded range of initial noise amplitudes, transitioning to uniform activation or inhibition at high noise depending on the existence of an activated steady-state, while Turing patterns formed robustly across the full noise range (**Fig. S5F-H**). This sensitivity held across the irregular regime, which spans both monostable and bistable parameter sets (**Fig. S5H**). To experimentally test sensitivity to initial conditions, we mixed pre-activated and non-activated cells at different ratios and plated at confluence (**Fig. S5I**). Pattern formation occurred at low pre-activated fractions (0-25%), but at high pre-activated fractions (50-100%) the entire culture activated uniformly without forming patterns (**Fig. S5J-K**). This initial-condition sensitivity is inconsistent with Turing patterning and consistent with the irregular regime interpretation.

Third, we experimentally measured the activator and inhibitor Hill coefficients (n_a_ and n_i_) and asked where the resulting pair places the implementation in the parameter space of Fig. 1J. To estimate the activator coefficient n_a_, we titrated the response of an anti-GFP synNotch receptor against varying concentrations of activator ligand in the absence of inhibitory branch (**Fig. S6A**). Increasing concentrations of GFP ligand produced a switch-like response that fit an activating Hill function with n_a_ ≈ 10 (**Fig. 2J, S6B**). This ultrasensitivity was directly visible in timelapse microscopy, where small changes in ligand concentration around the dose-response inflection point translated into all-or-none activation over time (**Fig. S6D-F**). The same approach applied to an anti-mCherry receptor with a different ligand-binding domain and intracellular effector yielded n_a_ ≈ 4 (**Fig. S6G-I**), indicating that the Hill coefficient depends on receptor architecture rather than being shared across synNotch implementations. To estimate the inhibitor coefficient n_i_, we provided increasing concentrations of a soluble anti-GFP inhibitor in presence of constant ligand concentration: this produced a gradual response with n_i_ ≈ 4 (**Fig. 2K, S6C**). The anti-GFP activation function is therefore substantially more cooperative than its inhibition function (n_a_/n_i_ ≈ 2.5). Mapping these values onto the phase space of Figure 1 places the anti-GFP implementation in a region dominated by irregular patterns (**Fig. 1J**, dashed box): for this Hill coefficient pair ∼99% of parameter sets that produce non-homogeneous patterns do so in the irregular regime (**Fig. S7A**). Together, these three lines of evidence place this synNotch implementation of JAPI in the irregular regime.

Based on this parametrization (n_a_ ≈ 10, n_i_ ≈ 4), we constructed a 2D computational model driven by continuous stochastic activation bursts, matching the nucleation pattern observed experimentally (Supplemental Movie 1, see Methods for model construction). Starting from initially homogeneous inactive cells, the parametrized model produced stable irregular patterns, whereas the JA control (no inhibition) instead showed runaway activation until uniform coverage (**Fig. 2L-M**, Supplemental Movie 2). The simulated patterns reproduced the experimental spatial signatures: a radially averaged autocorrelation that decayed monotonically without oscillation (**Fig. 2N**), a broad Fourier power spectrum lacking a sharp peak (**Fig. 2O**), and a wide distribution of arrested domain sizes comparable to that observed experimentally (**Fig. S7B**).

What patterning mechanism operates in the irregular regime, given that it is mechanistically distinct from Turing instability? Linear stability analysis classifies the homogeneous steady states in this regime as stable to small spatial perturbations: infinitesimal noise cannot grow into structured patterns. We turned to numerical simulations to explore the mechanism. We performed 1D simulations from a single localized activation burst of increasing magnitude and found that although at low magnitude the perturbation does dissipate, when it crosses a threshold, it stabilizes into an activated domain of finite width (**Fig. S7C-D**). Patterns could thus initiate from finite-amplitude events: stochastic transcriptional fluctuations occasionally drive a single cell to cross the activation threshold and begin producing the ligand, triggering activation of its neighbors and initiating a relay that propagates outward.

Once a positive relay has nucleated, what determines its size? Intuitively, as the activated domain expands, more cells produce the inhibitor; it then diffuses outward, accumulating in a halo around the domain. The expanding activation front stalls when the inhibitor concentration at the boundary reaches a critical threshold I_C_ sufficient to prevent further activation of neighboring cells (**Fig. 2P**). The arrested domain half-width R is therefore set by the balance between three quantities: the inhibitor’s spatial reach (set by the inhibitor diffusion length λ_i_), how strongly the domain produces inhibitor (the inhibitor production rate β_i_), and activation threshold I_C_. I_C_ is itself a derived quantity, set by the receiver cell’s activator-side parameters, including the activator production rate β_a_: a stronger activator drive raises the inhibitor level required to block activation, and therefore raises I_C_. The closed-form derivation in **Supplementary Note 4** yields R as a function of these three quantities: larger λ_i_ gives larger domains (inhibitor reaches further before decaying), larger β_i_ produces smaller domains (the boundary inhibitor concentration reaches I_C_ sooner), and higher I_C_ generates larger domains (the front can advance further before the inhibitor halo becomes sufficient to stall it). Numerical simulations confirm these dependencies (**Fig. S7E)**: domain size varies systematically with β_i_ (varying inhibitor production) and with β_a_ (which contributes to the receiver cell’s effective activation threshold I_C_. Numerical simulations also show that domain size responds to these parameters specifically when the activation function is more cooperative than the inhibitory function (**Fig. S7F-G)**, the regime our implementation occupies (n_a_ / n_i_ ≈ 10/4 ≈ 2.5).

The circuit architecture, parts measurements, regime classification, mechanism, and observed patterning are therefore consistent with a model in which random single-cell activation events nucleate locally, propagate to neighbors via juxtacrine relay, and arrest at finite domain sizes when accumulated inhibitor at the boundary reaches the critical threshold I_C_ (**Fig. 2P**). Given that irregular-regime features were observed across a broad range of parameter values and in both JAPI and PAPI implementations (**Fig. 1J, S1H**), and that the derivation in **Supplementary Note 4** is architecture-agnostic, the arrest mechanism seems to be a property of this class of reaction-diffusion systems rather than a JAPI-specific phenomenon.

Together, these results establish synNotch-mediated JAPI as functional, compact reaction-diffusion implementation in mammalian cells, which is sufficient for producing self-organized multicellular non-periodic patterns through nucleation, juxtacrine relay, and inhibitor-mediated arrest.

### Predictive tuning of JAPI domain size through modulation of inhibition and tissue geometry

Having generated a synNotch realization of the JAPI architecture and characterized its irregular patterning mechanism *in vitro*, we next tested predictions of the arrest mechanism (**Supp. Note 4**): that domain size depends on the inhibitor production rate β_i_ and the inhibitor diffusion length λ_i_, with stronger or slow-diffusing inhibition producing smaller domains.

To test the dependency of domain size on the inhibitor production rate, we started by observing that distinct clonal cell lines, initiated from a homogeneously inactivated state, produced widely different patterns ranging from small isolated domains to large labyrinth-like connected fields (**Fig. 3A**). We hypothesized that these differences reflect stochastic variation in transgene integration site and copy number across clones, which would translate into different production rates of the inhibitor and activator. Consistent with this, small changes in production rates of both species lead to similar pattern variations in 2D simulations (**Fig. 3B**). To test whether inhibitor production rate β_i_ specifically controls domain size, we used a fluorescent marker linked to the inhibitor expression cassette as a proxy for this parameter. Within a population of ∼20 clonal lines, the proportion of activated cells correlated inversely with reporter expression, indicating that clones with higher β_i_ produce smaller activation domains (**Fig. S8A**). To test this more directly, we bulk-sorted cells by reporter level into low, medium, and high β_i_ categories and quantified the resulting patterns: mean equivalent domain diameter decreased systematically as β_i_ increased, from approximately 150 µm in low producers to 50 µm in high producers (**Fig. 3C**, quantification; **Fig. S8B-C** for corresponding FACS plots and microscope images). Parametrized 2D simulations with individually tuned β_i_ recapitulated this dependence: stronger inhibitor production generated smaller domains *in silico* (**Fig. 3D** for quantifications; **Fig. S8D**, for simulated patterning outcomes), consistent with the arrest mechanism’s prediction that stronger inhibitor production stalls the activation front at a smaller domain size.

**Figure 3.**
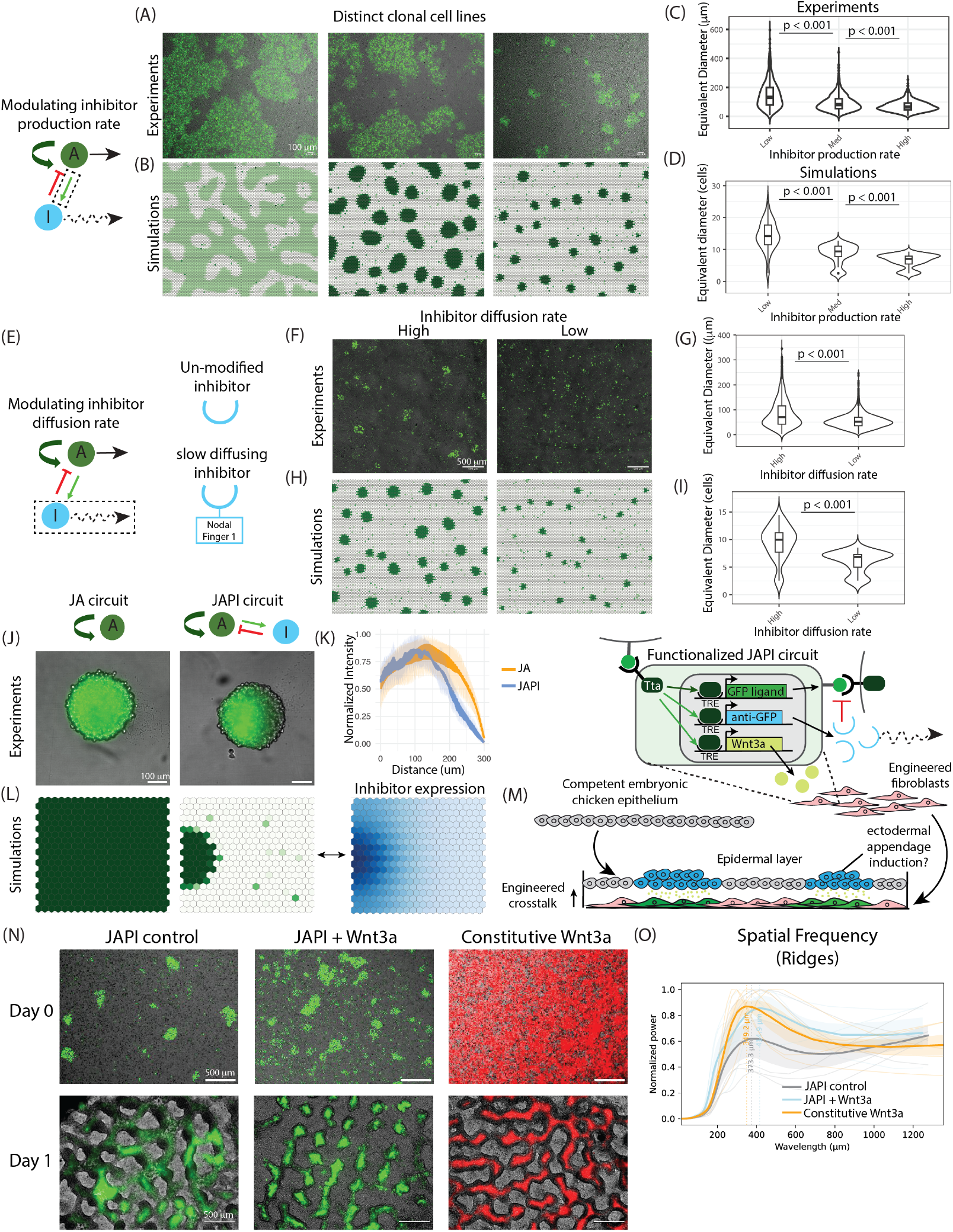
Parameter control of JAPI patterning and interfacing with embryonic chicken skin patterning. (**A-D**) Effect of the inhibitor production rate on JAPI patterning. Left schematic, the inhibitor production step in the JAPI circuit is highlighted by a dashed box. (**A**) Fluorescence microscope endpoint images at day 4 of three distinct clonal JAPI cell lines. Green indicates activated cells (GFP signal); brightfield in grey. Initial condition: homogeneous inactivated cell lawn. Scale bar = 100 µm. (**B**) Endpoint snapshots of two-dimensional JAPI simulations parametrized with the Hill coefficients from Fig. 2J-K (n_a_ = 10, n_i_ = 4) for three distinct parameter sets of activator and inhibitor production rates. Green intensity indicates dimensionless activator concentration (see Supp. Note 3 for description of dimensionless units). Simulation setup and parameter values per each simulation is in Methods, Numerical Simulations. (**C**) Violin plot of the distribution of equivalent diameters of activated domains, for three polyclonal cell populations sorted by reporter expression of the inhibitor-encoding transgene as a proxy for inhibitor production rate β_i_ (x axis: low, medium, high), over three independent replicates. y axis: equivalent diameter in µm. Box plot inside the violin shows median and interquartile range. p-values calculated from Welch’s Two Sample t-test. (**D**) Violin plot of the distribution of equivalent diameters of activated domains from parametrized JAPI simulations (n_a_ = 10, n_i_ = 4) with three values of inhibitor production rate β_i_ (x axis: low, medium, high), from three simulated replicates. y axis: equivalent diameter in cells. Box plot inside the violin shows median and interquartile range. p-values calculated from Welch’s Two Sample t-test. Simulation setup and parameter values in Methods, Numerical Simulations. (**E-I**) Effect of the inhibitor diffusion on JAPI patterning. (**E**) Left, schematic JAPI circuit diagram with the inhibitor diffusion step highlighted by a dashed box. Right, two variants of the inhibitor compared: top, the unmodified soluble anti-GFP nanobody dimer; bottom, the same nanobody dimer fused to the Nodal Finger 1 domain (drawn as a small box), reducing the inhibitor’s effective diffusion rate. (**F**) Fluorescence microscope endpoint images at day 4 of clonal JAPI cell lines containing either the original JAPI circuit (“High”, high inhibitor diffusion rate) or a JAPI circuit with the modified slow-diffusing inhibitor (“Low”, low inhibitor diffusion rate). Green indicates activated cells (GFP signal); brightfield in grey. Initial condition: homogeneous inactivated cell lawn. Scale bar = 500 µm. (**G**) Violin plot of the distribution of equivalent diameters of activated domains in the cell lines shown in (F), separated by inhibitor diffusion rate, from n = 3 independent replicates. Box plot inside the violin shows median and interquartile range. p-values calculated from Welch’s Two Sample t-test. (**H**) Endpoint snapshots of two-dimensional JAPI simulations parametrized with the Hill coefficients from Fig. 2J-K (n_a_ = 10, n_i_ = 4), comparing two values of inhibitor diffusion coefficient D_i_ (left, high; right, low) with all other parameters held constant. Green intensity indicates dimensionless activator concentration (see Supp. Note 3 for description of dimensionless units). Simulation setup and parameter values per each simulation is in Methods, Numerical Simulations. (**I**) Violin plot of the distribution of equivalent diameters of activated domains from the simulations in (H), separated by inhibitor diffusion rate (x axis: high, low), from three simulation replicates per condition. Box plot inside the violin shows median and interquartile range. p-values calculated from Welch’s Two Sample t-test. (**J**) Fluorescence microscope endpoint images at day 4 of spheroids aggregated from 500 cells in a U-bottom 96-well plate, for cells containing a juxtacrine self-activation (JA) circuit (left) or a JAPI circuit (right). Green indicates activated cells (GFP signal); brightfield in grey. Scale bar = 100 µm. (**K**) Line plot of normalized GFP intensity as a function of distance across the diameter of the spheroids in (J), for the JA circuit (red) and JAPI circuit (blue). Curves are means of n ∼50 spheroids from n = 3 independent replicates with shading for standard deviation. Image processing in Methods, Image Analysis. (**L**) Endpoint snapshots of two-dimensional JAPI simulations on a small domain (20 × 20 cells) from continuous stochastic activation bursts. Left, JA circuit. Middle, JAPI circuit, activator channel (green intensity indicates dimensionless activator concentration). Right, JAPI circuit, inhibitor channel (blue intensity indicates dimensionless inhibitor concentration). See Supp. Note 3 for description of dimensionless units. Simulation setup and parameter values per each simulation is in Methods, Numerical Simulations. (**M**) Schematic of the recombination experiment between embryonic chicken epidermis and engineered fibroblasts. Top, genetic circuit schematic for the functionalized JAPI circuit: GFP ligand, anti-GFP inhibitor, and Wnt3a are all downstream of synNotch activation. Middle, a fragment of competent embryonic dorsal chicken epidermis is placed on top of a lawn of engineered fibroblasts. Bottom, tested hypothesis: patterned Wnt3a secretion from the engineered fibroblasts influences ectodermal appendages in the overlying epithelium. (**N**) Fluorescence microscope images of engineered fibroblast cell lines alone (day 0, top row) and one day after recombination with embryonic dorsal chicken epidermis (day 1, bottom row). Columns label the circuits within the fibroblasts: left, JAPI control (no Wnt3a); middle, JAPI + Wnt3a (functionalized circuit from (M)); right, constitutive Wnt3a and mCherry expression. Green indicates activated cells in JAPI-containing lines (GFP signal); red indicates constitutive Wnt3a-mCherry expression; brightfield in grey. Scale bar = 500 µm. (**O**) Line plot of the Fourier power spectrum of the spatial frequency of epidermal ridges 24 hours after recombination with the three cell lines shown in (N): JAPI control (dark grey), JAPI + Wnt3a (light blue), constitutive Wnt3a (orange). Spatial frequency was computed from images such as the bottom row of (N), from n = 4 technical replicates. Light, thin curves are individual experiments, thick curves represent the average for each condition, and shading is standard deviation. Dotted vertical lines mark identified local maxima. Image processing in Methods, Image Analysis.

To test the dependency of domain size on the inhibitor’s spatial reach, set by the diffusion length 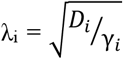, we chose to modulate D_i_ experimentally, which requires engineering the inhibitor molecule itself. The Finger 1 domain from the endogenous morphogen Nodal is known to substantially reduce its diffusion speed (*18*); inspired by this, we asked whether fusing the Finger 1 domain to our secreted anti-GFP nanobody dimer would reduce its diffusive reach (**Fig 3E**). In a localized inhibitor secretion assay, the fusion inhibitor showed reduced spatial reach in suppressing signal propagation compared to the unmodified inhibitor (**Fig. S8E-G**). JAPI cell lines incorporating the shorter-range inhibitor produced systematically smaller activated domains, with mean equivalent domain diameter decreasing from approximately 80 µm to 50 µm (**Fig. 3F-G**). Parametrized 2D simulations with reduced D_i_ reproduced this dependence (**Fig. 3H-I**), confirming the second prediction of the arrest mechanism: shorter inhibitor reach stalls the front at a smaller domain size.

Having confirmed the predictions of the arrest mechanism through parameter perturbations, we next asked how the architecture behaves in three-dimensional geometries. Theoretical analyses of reaction-diffusion systems have shown that when the domain is small relative to the inhibitor diffusion length, the natural outcome is a single activated and inhibited domain, as observed in left-right patterning mediated by the endogenous Nodal-Lefty PAPI circuit (*31, 32*). To test whether this holds in the irregular patterning regime of JAPI, we cultured JAPI cells in spheroids grown from approximately 500 aggregated cells. JAPI spheroids reproducibly generated a single activated and inhibited region, in contrast to the uniform activation observed in spheroids of inhibitor-free control cells (**Fig. 3J-K**). This single-domain outcome is consistent with the arrest mechanism: when the geometry is small relative to the inhibitor diffusion length, the halo of the first activated domain reaches all other cells before independent nucleation events can occur. 2D simulations in a domain of 400 cells with continuous stochastic activation bursts recapitulated the experimental result (**Fig. 3L**, Supplemental Movie 3).

Together, these experiments demonstrate that JAPI patterning responds predictably to perturbations of inhibitor production rate, inhibitor diffusion length, and tissue geometry, in agreement with the theoretically derived arrest mechanism.

### Functionalizing JAPI circuits for patterned Wnt3a secretion to interface with chicken epidermis in recombination assays

Having established that JAPI patterns can be modulated through parameter and geometry perturbations, we next asked whether engineered JAPI cells could interface with and perturb a natural developmental patterning system. As a paradigmatic test, we turned to feather bud formation in embryonic chicken skin, a classical periodic patterning system that can be recapitulated in explant cultures by combining the dermal mesenchyme and competent epidermis (*33, 34*). Feather bud patterning is itself driven by a reaction-diffusion architecture in which Wnt acts as a short-range activator while Bmp and Dkk act as a long-range inhibitors (*15, 35*), making it a natural test system for interfacing with a synthetic RD architecture.

We functionalized the JAPI circuit to drive expression of Wnt3a, a signaling factor that participates in ectodermal appendage formation, under the same activation-responsive promoter that drives the GFP ligand and anti-GFP inhibitor (**Fig. 3M**, top; **Fig. S9A-B**). The resulting clonal cell lines produced Wnt3a specifically from activated cell clusters, as confirmed by transcriptional profiling (**Fig. S9C-D**). We adapted a classical reconstitution assay in which dissociated chicken dermal mesenchyme is combined with dissociated epidermis to drive feather bud formation, replacing the dermal mesenchyme with engineered fibroblasts (**Fig. 3M**, bottom). We compared three conditions: JAPI circuits without Wnt3a output, functionalized JAPI circuits with patterned Wnt3a output, and cells constitutively expressing Wnt3a without circuit-mediated patterning (**Fig. 3N**). In all three conditions, the chicken epidermis produced regular morphological ridges, as expected for reconstitution with a dermal cell layer (*34*). The geometry of the ridges varied across conditions: we observed a trend of decreasing ridge area and connectivity with increasing Wnt3a production (**Fig. S9E-F**), and Fourier analysis revealed subtle differences in ridge spatial frequency across conditions (**Fig. 3O**). Examining the engineered fibroblast layer, we observed that the GFP distribution of the JAPI circuits reorganized into a more regular pattern that aligned with the overlying ridges in all three conditions, including the JAPI control lacking Wnt3a output (**Fig. 3N, S9G**), an unexpected observation, indicating that co-culture with the epidermis also influences the spatial organization of the engineered fibroblast layer. These observations suggest that engineered JAPI cells can interface bidirectionally with developmental tissue: Wnt3a produced by the engineered cells reaches and qualitatively alters feather bud patterning in the overlying epidermis, while contact with the chicken tissue reorganizes the spatial pattern of the engineered fibroblast layer. These experiments illustrate that synNotch-JAPI patterning can be extended to produce self-organized patterns of endogenous signaling effectors that can interact with native developmental processes.

### Multiplexed JAPI circuits generate four-state patterns from independent signal propagation with tunable positive cross-correlation

Having established synNotch JAPI as a functional reaction-diffusion patterning circuit, we next asked whether the architecture could be multiplexed to generate patterns with multiple cell states in a single cell population, beyond the two state (activated or inhibited) partitioning achieved from single JAPI circuits. The compact single-receptor architecture of JAPI circuits, combined with the orthogonality of synNotch receptor-ligand pairs (*23*), enables us to implement two orthogonal JAPI circuits in a single cell population, extending the architecture to multi-state patterns. To this end, we designed a dual-circuit implementation in which two synNotch-based JAPI architectures, distinguished by their ligands (GFP and mCherry), inhibitors (anti-GFP and anti-mCherry), and transcriptional effectors (Tta and Gal4), operate in parallel; we termed it dual-JAPI (**Fig. 4A**). When simulated in 2D with parametrized Hill coefficients and assuming uncorrelated initiation noise between the two channels, these circuits produced two independent patterns, with domain sizes tunable through the same production and degradation parameters as in single circuits (**Fig. 4B**). Whereas a single JAPI circuit produces a binary pattern of activated and inhibited cells, dual circuits generate patterns of four cell states: off, one circuit activated, the other activated, or both activated.

**Figure 4.**
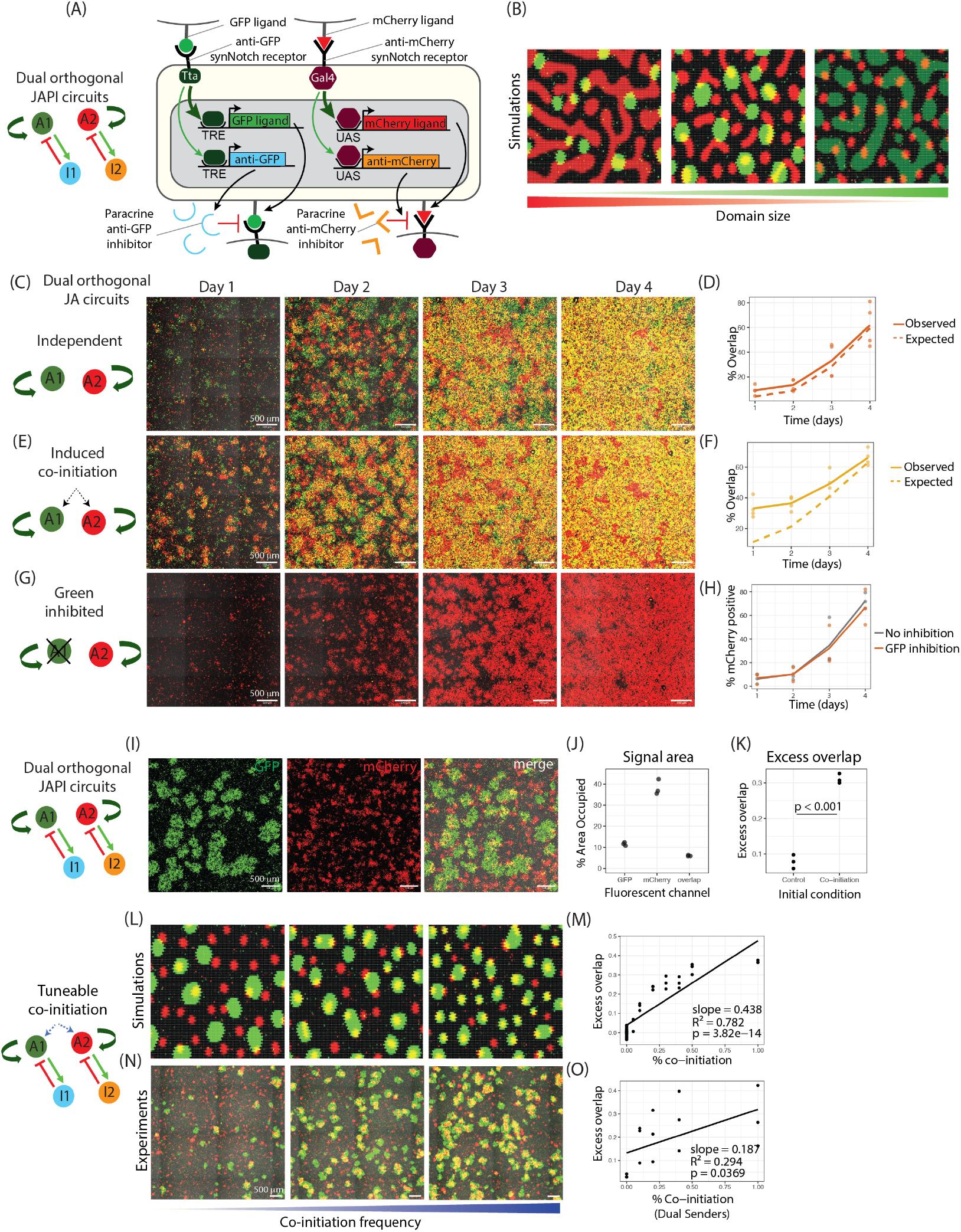
Multiplexing two JAPI circuits generates four-state patterns with tunable positive cross-correlation in a single cell population. (**A**) Schematic of dual orthogonal synNotch-based JAPI circuits engineered in a single cell line. Left, abstract conceptual depiction with two activators A1/A2 and two inhibitors I1/I2. Right, cellular implementation with parts combining the two individual JAPI circuits from Fig. 2A (anti-GFP-Tta) and Fig. 2E (anti-mCherry-Gal4). The anti-GFP-Tta synNotch receptor drives transcription from TRE promoters of GFP ligand (juxtacrine activator A1, green) and anti-GFP nanobody dimer (paracrine inhibitor I1, light blue). The anti-mCherry-Gal4 synNotch receptor drives transcription from UAS promoters of mCherry ligand (juxtacrine activator A2, red) and anti-mCherry nanobody dimer (paracrine inhibitor I2, orange). (**B**) Endpoint snapshots of two-dimensional simulations of two orthogonal JAPI circuits, parametrized with the Hill coefficients from Fig. 2J-K and S6H (n_a1_ = 10, n_i1_ = 4; n_a2_ = 4, n_i2_ = 4), shown for three sets of activator and inhibitor production rates that produce different relative domain sizes (left to right, indicated by the gradient bar below). Green intensity indicates dimensionless activator concentration for the GFP-based circuit; red intensity indicates dimensionless activator concentration for the mCherry-based circuit (see Supp. Note 3 for description of dimensionless units). Simulation setup and parameter values per each simulation is in Methods, Numerical Simulations. (**C-H**) Independent and co-induced initiation and signal propagation in dual JA circuits. (**C**,**E**,**G**) Fluorescence microscope images of the same field of view at the indicated timepoints of a timelapse experiment of cells containing two orthogonal juxtacrine self-activation circuits (JA), from a homogeneous inactivated cell lawn, either with: (**C**) no perturbations, (**E**) mixed with 1% cells constitutively expressing both GFP and mCherry ligands (dual senders), or (**G**) in medium containing tetracycline, which inhibits the GFP-based circuit. GFP signal in green, mCherry signal in red. Individual channels are shown in Fig. S10D,F,H. Scale bar = 500 µm. (**D**,**F**) Line plot of the percentage of cells positive for both GFP and mCherry over time, measured by FACS from the experiment in (**C**) and (**E**) respectively, comparing the observed value (solid line) to the value expected by independent probability (dashed line, computed as the product of the individual GFP-positive and mCherry-positive fractions). Each dot is an individual independent replicate (n = 4); lines connect replicate means. (**H**) Line plot of the percentage of mCherry-positive cells over time, measured by FACS from the experiments in (C) (‘No inhibition’, both circuits active) and (G) (‘GFP inhibition’, GFP-based circuit blocked by tetracycline). Each dot is an individual replicate; lines connect replicate means (n = 3 independent replicates). (**I-K**) Dual JAPI circuits operating in parallel in a single clonal cell line. Circuit architecture is shown to the left of the panel. (**I**) Fluorescence microscope endpoint images at day 4 of cells containing dual orthogonal JAPI circuits as designed in (A), showing the same field of view in three channels: GFP signal (left, green), mCherry signal (middle, red), and merge (right). Initial condition: homogeneous inactivated cell lawn. Scale bar = 500 µm. (**J**) Scatter plot of the percentage of area covered by each fluorescent signal in dual orthogonal JAPI cells from (I), measured at day 4. Each dot is an individual replicate (n = 3). Image processing in Methods, Image Analysis. (**K**) Scatter plot of excess overlap between GFP and mCherry signals in dual-JAPI cells initiated either spontaneously (Control) or with induced co-initiation from dual sender cells (Co-initiation), calculated as the difference between the measured DICE coefficient and the analytically estimated null value. Each dot is an individual replicate (n = 3). p-values calculated from Welch’s Two Sample t-test. DICE coefficient calculation in Methods, Image Analysis. (**L-O**) Tunable co-initiation in dual JAPI circuits. Left of the panel cluster, schematic of the circuit with the co-initiation step highlighted. (**L**) Endpoint snapshots of two-dimensional simulations of dual orthogonal JAPI circuits initiated with increasing co-initiation (left to right). Co-initiation is encoded by setting a defined number of cells co-expressing both activators at time zero. Green intensity indicates dimensionless activator concentration for the GFP-based circuit; red intensity indicates dimensionless activator concentration for the mCherry-based circuit (see Supp. Note 3 for description of dimensionless units). Simulation setup and parameter values per each simulation is in Methods, Numerical Simulations. (**M**) Scatter plot with linear regression of excess overlap as a function of co-initiation frequency, computed from simulations as in (L). Each dot is an individual simulation (n = 3 per condition); black line is the best-fit linear regression. Slope, R^2^, and p-value indicated in the figure. **(N)** Fluorescence microscope endpoint images at day 4 of cells containing dual orthogonal JAPI circuits (as in (I)), mixed with increasing numbers of dual senders (left to right: increasing co-initiation frequency, indicated by the gradient bar below). Each image shows the merged GFP (green) and mCherry (red) signals; separate channels are shown in Fig S11E. Scale bar = 500 µm. **(O)** Scatter plot with linear regression of excess overlap as a function of dual sender proportion, computed from experiments as in (N).. Each dot is an individual replicate (n = 3 per condition); black line is the best-fit linear regression. Slope, R^2^, and p-value indicated in the figure.

Although orthogonality in synNotch-mediated signaling has been demonstrated previously, it is unclear whether this independence extends to stochastic initiation of signal propagation, which underlies the JAPI patterning mechanism (**Fig. S10A**). To test this directly, without the confound of signal propagation arrest, we first examined activation dynamics in a cell line containing only the activator branches of two orthogonal juxtacrine feedforward circuits (GFP and mCherry), which we term dual-JA cells (**Fig. S10B-C**). Timelapse microscopy of clonal dual-JA cell populations showed that activation foci of the two channels arose spontaneously and in distinct cells (**Fig. 4C, S10D** for single channels). Quantitatively, the measured fraction of double-positive cells over time matched the prediction for two circuits initiating and propagating independently (**Fig. 4D, S10E**). We confirmed this independence with two complementary controls. First, forced co-initiation by addition of double sender cells (i.e. cells constitutively expressing both GFP and mCherry ligands) produced overlap above the independent prediction (**Fig. 4E-F, S10F-G**), demonstrating that elevated overlap is achievable under forced co-initiation and that the assay can detect overlap above the independent prediction. Second, suppressing selectively only the GFP-based circuit with tetracycline left propagation of the mCherry circuit unaffected (**Fig. 4G-H, Fig S10H-I**), ruling out cross-regulation through activation and revealing no measurable effects of transcriptional or translational saturation between the circuits.

With independence of stochastic signal initiation and propagation established, we built the full dual-JAPI cell line by adding both inhibitor branches: the paracrine anti-GFP and anti-mCherry inhibitor constructs, each driven by its respective transcriptional effector (**Fig. 4A, S11A**). From an initial homogeneous inactivated condition, these dual-JAPI cells spontaneously generated two-color, four-state stable patterns (**Fig. 4I**, Supplemental Movie 4). The two channels occupied different fractions of the total area and showed different autocorrelation profiles (**Fig. 4J, S11B**), likely reflecting differences in circuit parameters between channels. To quantify the spatial relationship between channels, we measured their overlap against a chance baseline that preserves each channel’s area fraction (Methods, “Image Analysis”); we term the deviation from chance the excess overlap, with zero indicating spatial independence. Dual-JAPI patterns were statistically quasi-independent, with only a slight positive excess overlap (**Fig. 4K**), and the radial cross-correlation decayed monotonically with no negative oscillations, ruling out spatial exclusion (**Fig. S11C**). The short-range positive elevation in dual-JAPI patterns likely reflected short-range cross-correlation between neighboring cells. In contrast, inducing co-initiation with dual sender cells markedly increased excess overlap (**Fig. 4K, S11D**).

Co-initiation therefore provides a handle for positive cross-correlation between dual-JAPI patterns, without genetic modification of the circuit. We tested whether this handle is tunable by varying the frequency of co-initiation events in simulations and experiments. In 2D simulations, seeding a defined number of cells with both channels active at the first timestep produced a proportional increase in cross-channel overlap (**Fig. 4L-M**). Titrating dual sender cells in dual-JAPI cultures produced the same trend: graded sender number generated graded overlap, reflecting an increased fraction of sender-driven co-initiation events relative to spontaneous independent initiations (**Fig. 4N-O, S11E** for single channels).

Together, dual-JAPI circuits generate spatial patterns of four cell states from two statistically quasi-independent patterning processes, with positive cross-correlation between them tunable through co-initiation frequency.

### A genetically encoded cross-inhibition library produces negative cross-correlation between coupled JAPI patterns

Having established that two JAPI circuits operate independently and that their cross-correlation can be manipulated through initial conditions, we next asked whether cross-correlations could be encoded in the circuit itself. Genetic encoding would enable autonomous appearance of correlated patterns intrinsically from a homogeneous initial condition rather than depending on external perturbations, extending the self-organization of dual-JAPI circuits to coordinated multi-channel patterns. To this end, we designed a cross-inhibitory dual-JAPI architecture in which activation of one circuit drives expression of the inhibitor of the other (**Fig. 5A**). The proposed implementation relies on introducing additional binding sites for the opposing circuit’s transcriptional effector into the promoter controlling inhibitor expression; in this way, activation of one circuit induces production of the inhibitor for the other. This logic can be realized as uni-directional coupling (one circuit inhibits the other) or bi-directional coupling (each circuit inhibits the other).

**Figure 5.**
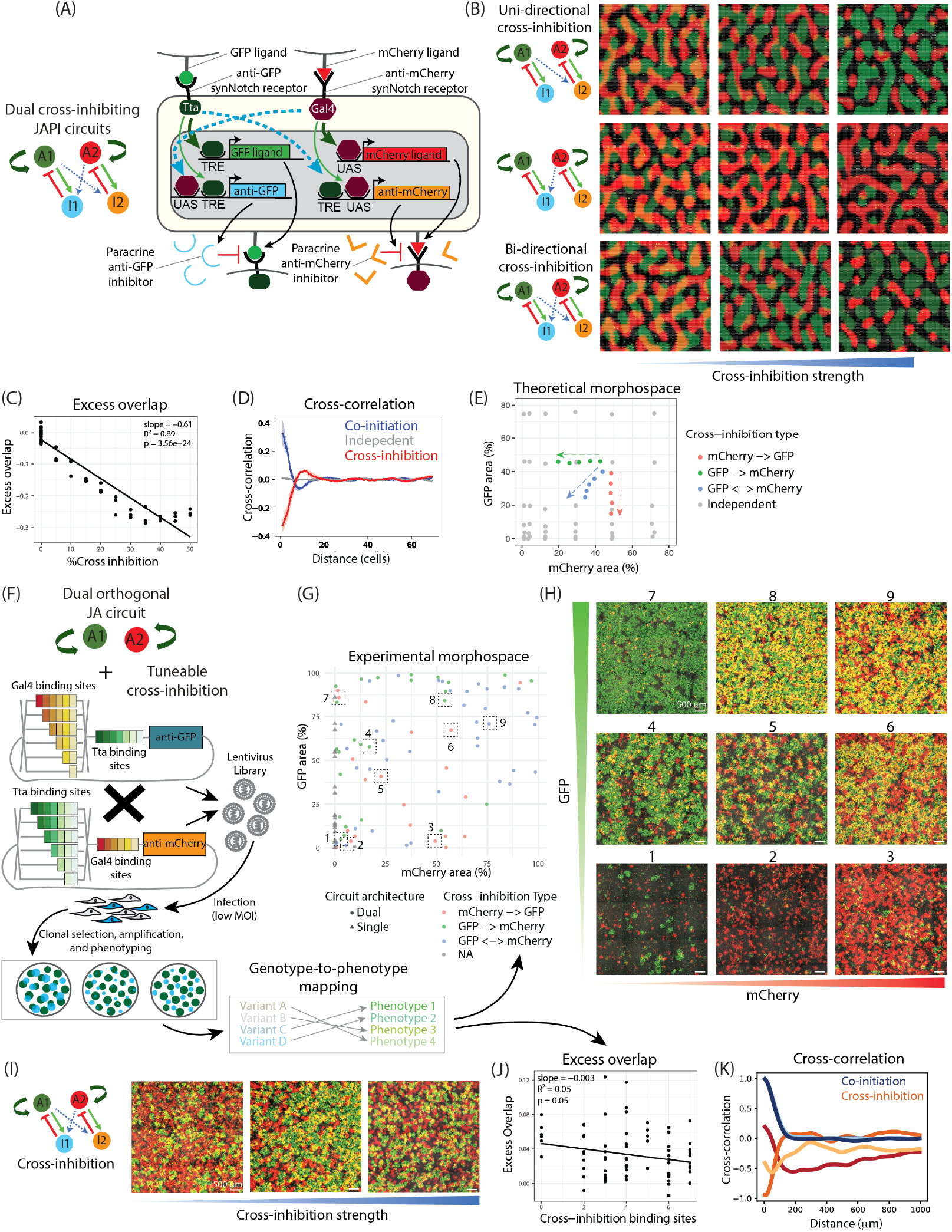
A genetically encoded cross-inhibition library produces negative cross-correlation between multiplexed JAPI patterns. (**A**) Schematic of dual cross-inhibiting synNotch-based JAPI circuits engineered in a single cell line. Left, abstract conceptual depiction with two activators A1/A2, two inhibitors I1/I2, and dashed blue arrows indicating cross-inhibitory coupling. Right, cellular implementation: each paracrine inhibitor is placed downstream of a dual-input promoter containing binding sites for both the Tta and Gal4 transactivation domains, allowing each inhibitor to be activated downstream of both synNotch receptors. The presence and arrangement of cross-binding sites determines the strength and directionality of cross-inhibition. (**B**) Endpoint snapshots of two-dimensional simulations of dual JAPI circuits with increasing cross-inhibition strength (left to right, indicated by the blue gradient bar at the bottom). Rows: top, uni-directional cross-inhibition from GFP circuit 1 to mCherry circuit 2; middle, uni-directional cross-inhibition from mCherry circuit 2 to GFP circuit 1; bottom, bi-directional cross-inhibition. Green intensity indicates dimensionless activator concentration for the GFP-based circuit; red intensity indicates dimensionless activator concentration for the mCherry-based circuit (see Supp. Note 3 for description of dimensionless units). Simulation setup and parameter values per each simulation is in Methods, Numerical Simulations. (**C**) Scatter plot with linear regression of excess overlap as a function of bi-directional cross-inhibition strength, computed from simulations as in the bottom row of (B). Each dot is an individual simulation (n = 3 per condition); black line is the best-fit linear regression. Slope, R^2^, and p-value indicated in the figure. (**D**) Line plot of the radially averaged cross-correlation between the GFP-based and mCherry-based activator signals, computed from simulations of dual JAPI circuits in three coupling regimes: independent (grey), induced co-initiation (blue), and cross-inhibiting (red). Curves are means with shading for standard deviation across replicates (n = 3 per condition). Simulation processing in Methods, Spatial analysis of 2D simulations. (**E**) Numerical morphospace scatter plot of the percentage of area covered by the mCherry-based circuit (x axis) versus the GFP-based circuit (y axis), from parametrized two-dimensional simulations of dual JAPI circuits. Grey dots, independent dual-JAPI circuits across a range of activator and inhibitor production rates (n = 36). Colored dots, dual-JAPI circuits with encoded cross-inhibition: red, mCherry → GFP; green, GFP → mCherry; blue, bi-directional (n = 5 each). Dashed colored arrows indicate the direction of movement in the morphospace as cross-inhibition strength increases, with arrow color matching the corresponding cross-inhibition type. Simulation setup and parameter values per each simulation is in Methods, Numerical Simulations. (**F**) Schematic of the experimental pipeline for constructing, testing and characterizing a library of dual-JAPI circuits with variable cross-inhibition. Starting from a parental cell line containing a dual orthogonal JA circuit (top), variants of the two inhibitor cassettes with different numbers of Tta and Gal4 binding sites in their dual-input promoters (color-coded barcodes) are combined to generate a lentiviral library. The library is delivered at low MOI to ensure single-construct integration per cell. Clonal cell lines are then generated, amplified, and phenotyped. They are then genotyped to map circuit variants to distinct patterning phenotypes. See Methods, Library screening of cross-inhibition circuits, for full pipeline. (**G**) Experimental morphospace of the percentage of area covered by the mCherry-based circuit (x axis) versus the GFP-based circuit (y axis), measured at day 4 for n = 85 independent dual-JAPI clonal cell lines. Dot color encodes the cross-inhibition type detected for each clone: grey, none (single-JAPI circuits); red, mCherry → GFP; green, GFP → mCherry; blue, bi-directional. Single GFP-based JAPI circuits are shown along the y axis (grey triangles, NA cross-inhibition type). Dashed boxes mark the clones whose patterns are shown in (H). (**H**) Fluorescence microscope endpoint images at day 4 of nine representative clonal cross-inhibiting dual-JAPI cell lines selected from the morphospace in (G) (dashed boxes), arranged to span the range of populated GFP-mCherry area combinations. Each image shows the merged GFP (green) and mCherry (red) signals; images of separate channels are found in Fig S14A. Initial condition: homogeneous inactivated cell lawn. Scale bar = 500 µm. (**I**) Fluorescence microscope endpoint images at day 4 of three cross-inhibiting dual-JAPI clonal cell lines with significant negative cross-correlation between GFP and mCherry signals, shown left to right with increasing cross-inhibition strength (indicated by the blue gradient bar below). Each image shows the merged GFP (green) and mCherry (red) signals; images for separate channels are found in S14B. Initial condition: homogeneous inactivated cell lawn. Scale bar = 500 µm. (**J**) Scatter plot with linear regression of excess overlap as a function of the number of cross-inhibitory binding sites in the inhibitor promoter across clonal cross-inhibiting dual-JAPI cell lines generated through the library approach, indiscriminately for uni- and bi-directional cross-inhibition. For cell lines with bi-directional cross-inhibition, the higher count between the two promoters is plotted. Each dot is an individual experimental observation (n = 76 measurements from n = 25 distinct genotyped clonal cell lines); black line is the best-fit linear regression. Slope, R^2^, and p-value indicated in the figure. (**K**) Line plot of the radially averaged cross-correlation between GFP and mCherry signals, computed from experimental patterns of dual-JAPI cell lines across the two non-independent coupling regimes shown in (D). Curves are individual replicates from independent initial conditions (shades of blue) or cross-inhibiting clonal cell lines (shades or red/orange). Image processing in Methods, Image Analysis.

We implemented this architecture computationally to ask whether cross-inhibition produces negative spatial coupling. In 2D simulations, increasing cross-inhibition strength progressively reduced overlap between the two patterns, in both uni- and bi-directional configurations (**Fig. 5B-C, Fig. S12A-B**). In uni-directional cases, the suppressed pattern was determined by the direction of inhibition; in bi-directional cases, both activated domains were reduced. Cross-correlation profiles revealed short-range negative correlation under cross-inhibition (**Fig. 5D, S12C-D**), in contrast to the positive correlation produced by co-initiation in **Fig. 4** and the near-zero correlation of independent dual-JAPI patterns. Cross-inhibition therefore generates a qualitatively new type of spatial coupling, negative cross-correlation.

To frame the simulation results, we placed them in a two-dimensional morphospace defined by the fractional area occupancy of each channel (**Fig. 5E**). Dual-JAPI patterns populate a large region of this morphospace from variation in circuit production rates alone, regardless of cross-inhibition (gray points). Cross-inhibition does not expand the morphospace – instead, it adds a directional axis within it: uni-directional GFP→mCherry cross-inhibition shifts patterns toward smaller mCherry domains; uni-directional mCherry→GFP cross-inhibition shifts toward smaller GFP domains; bi-directional cross-inhibition reduces both. The direction of the shift is set by the coupling geometry, and its magnitude increases with cross-inhibition strength (**Fig. 5E**, arrows). Cross-inhibition is therefore a directional intervention within the morphospace that stochastic parameter variation already populates, and because it is genetically encoded, the resulting shift arises autonomously from a homogeneous initial condition rather than being imposed externally.

To implement this architecture experimentally, we first tested whether dual-input promoters with binding sites for both Tta and Gal4 could functionally support cross-inhibition between the two circuits. Coating the culture surface with GFP forced activation of the GFP circuit and suppressed patterning by the mCherry circuit, and vice versa (**Fig. S12E-F**), confirming that the dual-input promoter responds to both transcriptional effectors and that activation of one circuit can drive expression of the inhibitor of the other at functional expression levels. With this design validated, we proceeded to systematically sample the cross-inhibition axis through a library approach. We generated a barcoded library of inhibitor-encoding constructs with tunable dual-input promoters (**Fig. 5F**). Each promoter contained a variable number of binding sites for the opposing synNotch receptor, enabling graded cross-inhibition strengths. The library was delivered as two lentiviral pools into cells containing dual independent JA circuits, allowing uni- or bi-directional cross-inhibition to be assembled combinatorially through infection with one or both pools alongside a single-promoter inhibitor construct. Infections were performed at low multiplicity (MOI ∼0.5) to favor single-copy integrations (**Fig. S13A**). Clonal cell lines were generated by FACS sorting and genotyped by targeted PCR amplification and barcode sequencing (see Methods, “Library Screening of cross-inhibition circuits”). As in single-circuit lines, self-organized patterns were visible in genetically identical colonies grown from single sorted cells (**Fig. S13B**). Individual clones reproduced their characteristic patterning across replicates (**Fig. S13C**), confirming that clone-level patterning is an intrinsic property of distinct integration sites rather than experimental variation. An initial screening of 86 clones populated a large region of the dual-activation morphospace (**Fig. 5G-H, S14A**), consistent with large sampling of parameters due to stochastic integration.

We then asked whether genetically encoded cross-inhibition translated into changes in spatial coupling between the two patterns. We recovered 17 unique genotypes from 25 successfully genotyped clones, covering ∼85% of the uni-directional and ∼20% of the bi-directional cross-inhibition configurations in the library. For each genotyped clone, we measured spatial overlap and short-range cross-correlation against the encoded cross-inhibition strength. Across the library, cross-inhibition correlated negatively with pattern overlap (**Fig. 5I-J, S14B**), with the trend conserved across uni- and bi-directional configurations and robust to different overlap metrics (**Fig. S14C**). Cross-inhibition clones generated patterns with negative short-range cross-correlation, distinct from the near-zero cross-correlation of independent dual-JAPI clones and the positive cross-correlation of co-initiated dual-JAPI (**Fig. 5K**). The library therefore validated both the substrate (morphospace breadth from stochastic variation, **Fig. 5G**) and the directional axis (cross-inhibition-driven shifts in overlap and cross-correlation), confirming that encoded cross-inhibition produces pattern correlation profiles qualitatively distinct from independent dual-JAPI circuits.

Together, the results in Figures 4 and 5 establish that synNotch-based JAPI circuits can be multiplexed to generate four-state patterns in a single cell population, and that the cross-correlation between the two channels can be controlled through two complementary handles: initial conditions, which tune positive cross-correlation without genetic intervention, and genetically encoded cross-inhibition, which produces autonomous negative cross-correlated patterns from a homogeneous initial condition. This represents, to our knowledge, the first synthetic implementation of multiple coordinated reaction-diffusion patterns in mammalian cells.

## Discussion

Two-species reaction-diffusion circuits have long admitted, in principle, several architectural implementations, including local activation and long-range inhibition, substrate depletion, and varied modes of spatial transport (*7, 10, 36–38*). The novel architecture introduced here, juxtacrine activation with paracrine inhibition (JAPI), produces self-organized patterning without requiring differential diffusion (**Fig. 1**). In comparison to classical paracrine activator paracrine inhibitor (PAPI) circuits, the change is structural rather than parametric: the diffusing activator is replaced in JAPI by a membrane-tethered activator that propagates by activating its neighbors through cell-cell contact. Mathematically, the activator diffusion coefficient is replaced by a juxtacrine coupling kernel: the linear stability operator retains the 2×2 structure of PAPI, with the kernel replacing the ratio of diffusion coefficients in determining, together with the reaction kinetics, the spatial scale of the emerging pattern. Surprisingly, despite this structural substitution, JAPI’s operator also admits Turing-type instabilities and accesses the same patterning regimes as the dual-diffusion case, with one fewer degree of freedom. Interestingly, this substitution changes how the activator’s spatial spreading is controlled. In PAPI, that spreading is set by the activator diffusion coefficient D_a_, a transport parameter that can in principle be tuned independently of the reaction kinetics. In JAPI, the activator does not diffuse; instead, an activated domain expands dynamically through contact relay, with an inhibitor-free maximum front velocity we denote v_a_ (Supp. Note 2). Unlike D_a_, v_a_ is not an independent parameter but an emergent property of the dynamics, jointly determined by the same reaction parameters that govern the rest of the circuit: the activator’s spreading is therefore entangled with the reaction kinetics in a way it is not in PAPI. A rigorous theoretical treatment of v_a_, only initiated here, including the conditions under which it admits a closed-form expression in terms of the underlying circuit parameters, would clarify the conditions under which the two architectures behave equivalently. Regardless, the functional interchangeability of diffusion and contact relay as modes of spatial propagation motivates the search for natural patterning circuits operating through the JAPI principle.

More broadly, the interchangeability of diffusion and contact relay suggests that, for multicellular patterning, juxtacrine and paracrine signaling are better treated within a common framework than as separate categories, as proposed independently by others (*21, 39–41*). In **Fig. S15** we sketch one possible formalism, in which each morphogen in a patterning circuit is specified as intracellular, membrane-tethered, or secreted, qualitatively setting its range and mode of transport, and each interaction is defined as acting on the same cell (cis), or on neighbors (trans). Within this framework, architectures typically treated as distinct, including PAPI, JAPI, lateral inhibition, single-morphogen reaction-diffusion circuits (*42*), and even positional-information readouts (*43*), emerge as variations on shared design principles, differing in the reactions between circuit components and in how those components are transported in space. Casting these architectures in a common representation renders their design space accessible to enumeration, an approach that has proven powerful within single transport modes (*27, 44–48*) but has not yet spanned diffusion- and contact-based circuits together. Whether this framework, or one derived from it, can support systematic enumeration across transport modes remains an open direction.

The JAPI circuits are an excellent target for synthetic biology efforts: they are compact, relying on only two effectors, yet can in principle generate complex emergent multicellular phenotypes without requiring two differentially diffusing species. In the implementation developed here (**Fig. 2)**, a single synNotch receptor drives expression of both its own membrane-tethered ligand, encoding juxtacrine self-activation, and a paracrine inhibitor. Both effectors are controlled by the same activation-responsive promoter downstream of synNotch, yielding a two-component circuit coupled through a single transcriptional channel. In L929 fibroblasts, this compact design is sufficient for generating self-organized multicellular patterns from initially homogeneous cell populations. Previous implementations have required either added molecular complexity through three-or-more-component circuits in prokaryotes, or additional components to process signals from an endogenous activator-inhibitor morphogen pair in mammalian cells (*16, 18, 49*). Or work here demonstrates that self-organized multicellular patterning in mammalian cells via reaction-diffusion circuits does not require complex regulatory networks or finely tuned differential diffusion. To our knowledge, this synNotch-based JAPI design represents the most compact engineered reaction-diffusion circuit reported so far. These synthetic, orthogonal circuit designs provide a powerful framework for testing whether minimal circuit architectures are sufficient to generate target phenotypes, in the absence of the dense, intertwined regulatory interactions characteristic of natural systems. They also pose a direct challenge to established theoretical and computational models: to what extent do their predictions hold when implemented in stochastic, complex living systems?

These specific implementations of JAPI with synNotch produce irregular, non-periodic patterns *in vitro*, a regime of reaction-diffusion patterning that has received limited theoretical and experimental attention. Linear stability analysis predicts three outcomes around the activated homogeneous steady state: homogeneous activation, homogeneous inhibition, and periodic Turing patterns. Numerical simulations reveal a fourth, consisting of irregular non-periodic domains that occupy a region of parameter space linear stability classifies as stable (*29, 30*) - the patterns we observe *in vitro* fall into this regime. Mechanistically, we hypothesize that these patterns are not driven by a diffusion-driven instability but by stochastic nucleation of finite-amplitude perturbations that propagate and arrest when inhibitor accumulation at the boundary suppresses further activation. This distinguishes the mechanism from stochastic Turing patterning (*16*), where noise seeds an underlying periodic instability: in JAPI, the homogeneous state is linearly stable, and patterns form only when perturbations are large enough to escape it, not by amplification of noise around an unstable mode. Building the synNotch-based JAPI implementation provides the first experimental realization, to our knowledge, of this regime in a synthetic patterning system, making it amenable to deliberate engineering.

Theoretically, within the irregular regime, activated domain size is set by the balance between activation front propagation, governed by v_a_, and inhibitor accumulation, rather than by wavelength selection. The model therefore predicts that domain size should decrease as inhibition is strengthened or its range shortened, with no role for a selected wavelength. Both predictions held experimentally (**Fig. 3**): increasing the inhibitor production rate, accessible through transgene copy-number, and reducing the inhibitor’s effective range by fusing the Nodal Finger 1 domain to the secreted inhibitor both systematically reduced domain size. In our numerical screens, this irregular regime occupies a substantially larger region of parameter space than the periodic Turing regime and is correspondingly less sensitive to fine parameter tuning - this raises the possibility that reaction-diffusion logic may be more widespread in biological contexts than the Turing case alone implies. Whether natural systems use the irregular regime, and what experimental signatures would distinguish arrest-driven from wavelength-driven patterning *in vivo*, is an open question. We note that our simulations indicate that JAPI should also access the classical periodic Turing regime under appropriate parameters; however, we did not observe periodic patterns experimentally in our implementation, and whether they can be produced from reconstructed circuits in the stochastic environment of living tissue remains an open question.

Given the tunability of the resulting pattern’s properties and the orthogonality of synNotch signaling to endogenous pathways, we explored whether synthetic JAPI circuits could interface with natural patterning systems through self-organized patterns of morphogen secretion (**Fig. 3**). Unlike current synthetic organizers, which typically function as an engineered point source of endogenous morphogens (*50–52*), JAPI circuits could generate self-organized morphogen patterns capable of interacting with endogenous patterning systems directly at the level of their own spatial organization, including wavelength, domain size, and spacing. To test this, we functionalized the circuit by coupling Wnt3a expression to synNotch activation and co-cultured these cells with embryonic chicken epidermis: in this configuration, Wnt3a-JAPI cells altered the spatial frequency of feather bud precursor ridges. The interaction was non-trivial and bidirectional: the synthetic output influenced the epidermis while contact with the epidermis reorganized the engineered cell layer, which may be due to the involvement of other endogenous signaling or non-signaling processes such as mechanical coupling or cell migration (*53, 54*). To our knowledge, this is the first demonstration that a synthetic reaction-diffusion circuit can operate at the interface with an endogenous developmental system. The L929 fibroblasts used here lack intrinsic differentiation potential, so future directions include implementing these circuits in cells that participate directly in morphogenesis, such as primary dermal fibroblasts for ectodermal appendage formation or pluripotent stem cells for controlled symmetry breaking during differentiation. More broadly, combining the self-organizing properties of synthetic patterning circuits with the organizing activity of morphogens may provide a route toward programmable pattern formation and morphogenesis in organoids and, ultimately, *in vivo*.

Finally, we demonstrate that synNotch-based synthetic JAPI circuits can be multiplexed in mammalian cells for self-organization of complex multi-state patterning systems (**Figs 4-5**). Two orthogonal JAPI circuits operating in parallel produced spatially independent patterns, and their cross-correlation could be tuned positively through co-initiation or negatively through engineered cross-inhibition. We constructed and screened a barcoded library of dual-JAPI variants with varying cross-inhibition strengths, reading out each clone by the spatial pattern it formed. The resulting morphospace was broadly populated through a combination of stochastic parameter variation and genetic cross-inhibition. To our knowledge, this is the first library screen of engineered patterning circuits by their emergent spatial output in mammalian cells - previous mammalian circuit libraries have screened scalar readouts such as reporter gene expression (*55, 56*). Building this library demonstrates that synthetic patterning circuits, once compact and orthogonal, become amenable to library-scale, phenotype-based screening that conventional design-build-test cycles cannot easily reach. Demonstrated here for two circuits of the same architecture and at sparse library coverage, this approach nonetheless hints at a reframing of synthetic multicellular patterning not as the construction of individual circuits, but as the systematic exploration of an architecture-level design space. Within this space, JAPI is one possible architecture and synNotch one potential molecular substrate. The theoretical framework sketched in **Fig. S15** maps a wider potential circuit design space: a natural extension of the library would screen not only parameter variations but also across distinct circuit architectures. Our work demonstrates the feasibility of phenotypic circuit-level screening; the framework points at the larger space such screening could one day traverse.

Combined with the functionalization strategies presented here, and extended to non-signaling modules controlling adhesion (*24, 57, 58*), proliferation (*59*), migration (*60*), and differentiation (*61*), these tools point toward engineering self-organized tissues composed of spatially structured and functionally specialized cell populations. Inspired by the conceptual ambition of understanding-by-building multicellular systems, this work aspires to contribute to the nascent field of synthetic developmental biology.

## Author contributions

L.M. and B.S. designed the project, acquired funding, and wrote the manuscript; L.M. supervised the research; L.M. performed the mathematical analysis with support from D.J.G.P and B.S.; B.S. performed the experiments with support from J.J.D, K.P., and S.M; B.S. performed the numerical simulations with help from S.B.T., M.K. and L.M.; B.S. performed data analysis and image analysis with help from P.S.B. and J.J.D; T.-X.J. and B.S. performed the chicken recombination experiments with supervision from C-M.C.; Z.A.K. constructed the plasmids for the library assay, with supervision from I.M.E.

## Acknowledgments

We thank all past and present Morsut lab members, Michael Elowitz and Sheng Wang for critical discussions on this project; Bernadette Masinsin and Jorge Contreras from the Flow Cytometry Facility, and Seth Ruffins from the Optical Imaging Facility of the Eli and Edith Broad CIRM Center; Justin Bois and Michael Elowitz for making their teaching materials for mathematical modeling of biological systems publicly available; Matt Thomson and Megan McCain for scientific discussion on the project and the manuscript.

## Data, code, and materials availability

All code used for reaction–diffusion simulations is availible at https://github.com/BenSwedlund/reaction_diffusion_simulations under a MIT open license. The parameter sets chosen for representation across all figures can be found in supplemental table 1 and 2 for single-pattern and dual-pattern simulations, respectively. Scripts use to analyze single and dual circuit simulation outcomes were placed in https://github.com/BenSwedlund/reaction_diffusion_simulations/tree/main/2D_simulations/res_analysis. Image analysis pipelines are available at https://github.com/BenSwedlund/Image_analysis_JAPI. All plasmids used to construct the transgenic cell lines presented in this study are listed in supplementary table 3, and will be deposited to Addgene, including detailed annotations and whole-plasmid sequencing. The complete sequences of the dual-input promoters engineered for the library assay are provided in supplementary table 4. Stable cell lines generated in this study are available from the corresponding authors upon reasonable request. The source data underlying the figures and raw imaging datasets will be made openly available upon acceptance and publication.

## Methods

### NUMERICAL SIMULATIONS

All simulations were performed using Python, and all code used was deposited at https://github.com/BenSwedlund/reaction_diffusion_simulations under an open MIT license. The parameter sets chosen for representation across all figures can be found in supplemental table 1 and 2 for single-pattern and dual-pattern simulations, respectively.

#### 1-dimensional simulations

We simulated the described activator-inhibitor equations using a finite-difference integration method on a domain comprised of 100 units. Activator and inhibitor concentrations were updated according to Hill-type reaction kinetics coupled with diffusion. Diffusion was implemented using a discrete second-order finite difference approximation of the Laplacian. In the case where the activator acted through juxtacrine signaling, the diffusion of the activator was set to zero, and the input of the reaction function became the average activator expression of its direct two neighbors instead of itself. We chose to use the average instead of the sum of neighbor activation to render the code more resilient to changes in the number of neighbors, such as when switching from 1D to 2D. Zero-flux (Neumann) boundary conditions were imposed at the domain edges. Simulations were initialized from activator concentrations either as random numbers between 0 and a defined maximal threshold, small fluctuations (5%) around the calculated steady state, or a point activation of a user-defined height in the center of the one-dimensional field. Steady states were obtained by numerically solving the fixed-point equation in the absence of diffusion and spatial heterogeneity in expression levels. Among the resulting solutions, we selected the non-zero root with negative reaction Jacobian eigenvalues, excluding the stable point (a,i) = (0,0). Root finding used a safeguarded Newton/bracketing scheme with refinement to 10^−3^ precision. Time evolution was computed using an explicit Euler integration scheme with fixed time step Δt = 0.01 and spatial step Δx = 1 until convergence, defined as the average change in concentration per cell between two consecutive recorded time points (interval = 200 steps) falling below a specified threshold (10^-4^). If convergence did not occur, the simulation was stopped after 50000 steps.

#### Defining Patterning Regimes

Batch simulations were run using USC’s Center for Advanced Research Computing (CARC) cluster. We performed permutations of the five non-dimensionalized parameters (Ω_a_, Ω_i_, Ψ, n_a_, n_i_) as defined in the supplemental text, for simplicity leaving the diffusion rate of the inhibitor for JAPI and PAPI simulations which we set to D_i_ = 10. Parameter ranges were 1-10 for both production rates, 0.1-10 for the degradation rate, and 1-12 for both hill coefficients, for a total of 273600 parameter permutations. Parameter values were spaced linearly with an interval of 1, except for an interval of 0.1 for the degradation rate between 0.1 and 1, to screen cases both where the activator or the inhibitor is more stable than the other. Regimes were defined using simulations started from (i) small random perturbations around the calculated activated steady state, and (ii) an initial central spike of activator concentration. Four regimes were thus defined: all off, all on, Turing, and non-Turing patterns. (i) was used as a primary classification of Turing patterns, defined by a stable steady state that becomes destabilized in the presence of diffusion as inferred from the simulations. (ii) was used to determine alternative outcomes: homogeneous outcomes were determined by either the decay of the local perturbation back to zero (all off) or its propagation until the edges of the field (all on). Irregular patterns were defined by a size-constrained signal propagation from a localized perturbation – meaning the result was a single, stable centered gaussian-like activator concentration profile. The width of these curves was quantified for each spatial profile as the standard deviation of the activator distribution along the domain after baseline subtraction. Width measurements were then averaged across simulations and summarized as a function of the activator and inhibitor Hill coefficients. To reduce boundary-dependent artifacts, we repeated the spike simulations in a longer domain and compared matched parameter sets between the shorter and longer domains (100 vs 200 domain length). For irregular patterns, we retained only those parameter sets whose interior spatial profiles were highly consistent across domain lengths, using a normalized root-mean-square error threshold computed on corresponding central regions of the profiles. Additional parameter sets that led to multiple regular peaks of activation from a local perturbation and were not yet categorized as Turing patterns were added to this category. The same classification scheme and simulation parameters were then applied to simulations using a paracrine activator with fixed diffusion rate D_a_ = 1, and the two sets of classified regimes were compared between the two types of activator-inhibitor circuits across all parameter permutations. Monostability and bistability were defined using the steady state solver defined above – if a second stable steady state was found outside the inactivated point (a,i) = (0,0), the system was determined as bistable for this specific parameter combination.

#### Defining sensitivity to initial conditions for the different patterning regimes

To quantify the sensitivity of patterning outcomes to initial conditions, we performed simulations across representative parameter sets of three discrete regimes (Turing, irregular monostable, irregular bistable) while varying the magnitude of initial perturbations. For stochastic initial conditions, activator concentrations were initialized independently at each grid point from a uniform distribution U (0,T), where the noise amplitude T was varied from 0.1 to 10 in increments of 0.1. For localized perturbations, simulations were initialized with a central activation peak of height T, with T varied over the same range and increments.

#### 2-dimensional simulations – single circuits

Two-dimensional simulations were performed in Python using a hexagonal lattice implemented in an even-r offset layout, where each cell interacted with up to six nearest neighbors. Boundary sites were updated using only in-bounds neighbors, with out-of-range neighbor slots masked to prevent flux across the domain edges. Reaction terms were identical to those used in the one-dimensional model and were integrated with an explicit forward-Euler scheme using fixed time step Δt = 0.01 and spatial step Δx = 1.0. Activator signaling was implemented in either a juxtacrine or paracrine manner. As above, in the juxtacrine case, the activator input to the reaction term was defined as the mean activator concentration of the six neighboring cells, and activator diffusion was set to zero. In the paracrine case, the activator input was the local activator concentration, and activator diffusion was included with diffusion coefficient Da using a discrete Laplacian on the hexagonal grid, similar to the implementation of the inhibitor diffusion. Simulations were run until the mean absolute change in concentration per cell over successive saved time points fell below a predefined stopping threshold (10^-4^), or up to 50000 steps for figure 1, and 5000 steps for all other panels. Representative panels for the four defined regimes were generated from random uniform activation between 0 and 2, for fixed Hill coefficients n_a_ = 3 and n_i_ = 3, inhibitor diffusion coefficient D_i_ = 10, and in the case of a paracrine activator, an activator diffusion coefficient D_a_ = 1. For the parametrized models with experimentally measured Hill coefficients (n_a_ = 10 and n_i_ = 4), instead of a fixed initial condition, the field was initialized as homogeneous inactivation. To capture more accurately the continuous nature of our experimental system, continuous stochastic activation was modelled as a Poisson process in which each cell independently fired with probability λ*Δt per time step. When a firing event occurred, the activator concentration in that cell was increased by a fixed increment: either the calculated steady-state value, or the value of the parameter spike_value.

#### 2-dimensional simulations – coupled circuits

To investigate interactions between competing patterning systems, we extended the two-dimensional hexagonal model to include two coupled activator-inhibitor pairs evolving on the same hexagonal lattice, sharing common neighborhood topology and boundary conditions. Each pair followed the same reaction-diffusion dynamics described above, governed independent parameter sets p_1_ and p_2_. Diffusion was implemented separately for each species and each pair, following the same discrete Laplacian scheme as in the single-pair simulations. Stochastic continuous Poisson-distributed nucleation events were incorporated independently for each pair, as described above. Parameter sets used to generate the figure panels are summarized in supplemental table 2. To model shared initial signal propagation foci in the presence of continuous stochastic noise, initial conditions were defined using a specific input number of shared activation foci, where the initial concentration of both activators was set to a high value (20). Cross-inhibitory coupling between the two systems was implemented at the level of inhibitor production: the inhibitor production rate of each pair included an additional reaction term, calculated by multiplying the Hill response function of the opposing pair by a defined cross-inhibition rate. The cross-inhibition rates were set as a fraction of each circuit’s own inhibitor production rate, so that a given percentage represents the same relative strength across circuits with different β_i_.

#### Spatial analysis of 2D simulations

Scripts use to analyze single and dual circuit simulation outcomes were placed in https://github.com/BenSwedlund/reaction_diffusion_simulations/tree/main/2D_simulations/res_analysis. Fourier power spectrum, radially averaged autocorrelation and cross-correlation were quantified using the same metrics as described in “Image Analysis” below, computed directly from the final activation concentration values at each point. To calculate the excess overlap as defined below, the final concentration profiles were first converted to a binary mask using Otsu thresholding method (*62*).

### CELL LINE ENGINEERING

Plasmids used in this study are listed in supplementary table 3 and will be made openly available through Addgene.

#### Molecular Cloning

All inserts were cloned using Gibson Assembly. Briefly, template vectors were cut using restriction enzymes, inserts were PCR amplified (Takara, 639298), and both were run on a 1% agarose gel. Amplified DNA bands were cut and subsequently purified using a column extraction kit (Takara, 740611) before Gibson assembly (NEB, E2621L). The assembly product was introduced into chemically competent bacteria (NEB, C3019H). Colonies were then grown overnight in the antibiotic corresponding to the resistance of the plasmid (usually ampicillin), and the plasmid was subsequently purified by miniprep (Zymo, D4016), quantified using a nanodrop and sequence verified using whole plasmid sequencing.

#### Cell culture

HEK 293-T cells (Takara 632180) and L929 fibroblasts (ATCC# CCL-1) were cultured on DMEM (Thermo 11965092) containing 10% FBS (Genesee Scientific 25-514) and 1% penicillin-streptomycin (Thermo 15140122). Passaging was performed every 2-4 days using PBS (Gibco, 14040133) rinse followed by incubation with TryPLE (Gibco, 12563029).

#### Virus production and cell line generation

Lentivirus particles were produced from 2nd generation transfer plasmids (pHR) co-transfected with the packaging and envelope plasmids (psPAX2 and pVSV-G). HEK 293-T cells were used to produce the viral particles. The transfection was performed using LTX (Thermo, 15338100) as per the manufacturer’s protocol. Specifically, 1250 ng of target lentiviral vector was mixed with 800 ng of psPAX2 and 450 ng of pVSV-G in 250 uL optiMEM supplemented with 2.5 uL plus reagent. Separately, 7.5 uL LTX was diluted in 250 uL optiMEM. Both solutions were then mixed together and left to incubate for ∼10 minutes, then added on top of 1.5 million HEK cells plated on the same day on a 6-well. Viral particles were collected at day 3 and filtered using a 0.45 um PES filter.

#### Clonal cell line generation

L929 transceivers were infected with two to three distinct doses of non-concentrated virus, ranging from 10 uL to 1 mL. Cells with constitutive expression of ligands or inhibitors were bulk-sorted based on the expression of the fluorescent reporters. For cell lines containing patterning circuits, clonal cell lines were derived from single FACS-isolated cells deposited in 96-well plates, selected based on the expression of fluorescent reporters. Additionally, self-activation was used to positively select for cells containing both a synthetic Notch receptor and a downstream juxtacrine activation cassette. For circuits comprising more than one construct, vectors were inserted in the cells two-by-two, generating a matrix of virus concentrations. Fluorescent reporter expression and self-activation were used to determine an optimal virus dosage, from which the clonal cell lines were amplified. After 10 days, clonal cell lines were screened by fluorescence microscopy in the 96-well plates for the presence of the desired self-organized phenotype, before being expanded and cryopreserved.

#### *IN VITRO* Patterning experiments

Cells containing GFP-based JA or JAPI circuits were continuously repressed during maintenance by the addition of 100 ng/mL Tetracycline to the medium, to repress the intracellular domain of the corresponding synNotch receptor, Tta-VP64. To maintain cells containing mCherry-ligand/Gal4-based circuits in an inactivated state, these cells were cultured in 5 uM DAPT (Sigma D5942), a g-secretase inhibitor that inhibits both synthetic and natural receptors. At the beginning of the experiment, cells were seeded at an initial density of 630 cells/mm^2^ (0.5X confluency) in medium containing 2% FBS and no inhibitors. The lower FBS concentration was used to reduce proliferation speed, as we have shown that over-confluent cultures signal through synNotch signaling less efficiently^1^. For single GFP circuit patterning experiments, a 3 mg/mL fibrin gel was added on top of the cells by mixing thrombin (Sigma, T4648) diluted in 2% FBS medium and fibrinogen (Sigma, F8630) 6mg/mL diluted in PBS at a 1:1 ratio. The presence of the gel was used to minimize disruptions in inhibitor diffusion when multiple imaging steps were taken. Cells were imaged every day using a Zeiss Axio Observer.Z1 microscope. No gel was added in the medium for cells containing the mCherry circuit, and the plates were images as a single snapshot at day 4. Live imaging was performed using a Cellcyte 1 microscope, enabling continuous imaging without inducing movement of the plate (as it is the objective that moves).

#### Global Inhibition of JA cells using inhibitor-containing conditioned media

To globally test the ability of a paracrine anti-GFP nanobody dimer to inhibit signal initiation and propagation in a feedforward juxtacrine activator circuit, 1 million cells constitutively secreting the inhibitor were cultured for one week in a T75. This medium was then transferred onto JA cells during a signal propagation experiment as described above. Quantification of GFP positive cells was performed by Flow cytometry.

#### FACS-sorting and replating experiments

To extract cells from a fibrin gel at the endpoint of an experiment, the gel was dissociated using Nattokinase (Amazon, B09LRLLGCZ1). One capsule was diluted in 10 mL PBS and filtered using a 0.45 um filter. Cells were incubated with the Nattokinase for 30 minutes at 37°C, then manually dissociated. GFP and mCherry were used to gate activated and inactivated cells, and 60000 cells of each population was replated in separate wells of a 24-well plate. After 24 hours, the cells were imaged and a 3 mg/mL fibrin gel was added on top. Cells were imaged again 4 days after sorting. Quantification of signal area was performed using the pipeline described in “Image analysis”.

#### Localized inhibition source experiments

To form a localized source of inhibitor production, cells constitutively expressing a paracrine inhibitor were seeded as a 10 uL drop containing 10000 cells at the edge of a 24-well. After one hour in the incubator to let the cells settle, JA cells were seeded globally in the well and over the drop at 630 cells/mm^2^ (0.5X confluency). A 3 mg/mL fibrin gel was added the next day, and the wells were subsequently imaged with both a 2X and 10X objective on a Keyence BZ-X710. Quantifications were performed on images treated with Fiji’s subtract background (rolling ball = 50), by manually drawing a line from the edge of the well containing the locally seeded cells until the opposing end of the well, and computing the fluorescence signal over this line using Image J.

#### Dual localized cell seeding experiments

JA and JAPI were manually seeded as 10 uL drops as close as possible without leading to the fusion of both droplets. Drops were placed in the incubator for one hour to let the cells settle down. Then, the well was filled with culture medium. The next day, the medium was replaced with a 3 mg/mL fibrin gel. Signal quantification was performed as described below, in “Image Analysis”.

#### Spheroid experiments

Cells were aggregated in U-bottom 96-well plates (Greiner, 650970) at a concentration of 500 cells per well, in 100 uL 10% FBS medium, avoiding the edge wells and instead filling them with PBS. Spheroids were imaged at day 4 using a Zeiss Axio Observer.Z1 microscope. GFP signal over the diameter of the spheroids was measured by drawing a 50 pixel-width line across the spheroids and measuring the intensity of GFP signal across this line. We quantified this signal intensity over the diameter for a total of 100 spheroids, spread over 2 circuits (JA, JAPI) and 3 independent experiments.

### IMAGE ANALYSIS

Scripts for image pre-processing and analysis were deposited at https://github.com/BenSwedlund/Image_analysis_JAPI. For quantification of GFP signal over time in live imaging, the GFP signal was simply thresholded and the resulting area covered by positive signal was computed over each timeframe. For all other patterning experiments, fluorescence images were processed using a custom ImageJ macro. Binary masks were generated through two sequential thresholding and Gaussian smoothing steps, followed by iterative dilation and erosion to smooth object boundaries and remove very small objects. The binary masks were used for all image analysis, including calculating the fraction of the area occupied by a signal. The equivalent diameter of separated features was calculated from the measured area as 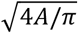. The 2D autocorrelation of an image was calculated using the fast Fourier transform method. To calculate the mean autocorrelation as a function of distance from the origin, referenced above as the radially averaged autocorrelation, we calculated each pixel’s distance from the origin and binned the pixels by distance into 100 bins, up to a maximum distance of 1 mm. The mean autocorrelation and its confidence interval was calculated from the values inside each bin. The cross-correlation measures the spatial covariance between different species in the same sample and was calculated similarly using the Fourier transform method. Radial cross-correlation was calculated similarly to radial autocorrelation, using 100 bins up to a maximum distance of 1 mm.

To further characterize the spatial colocalization between two fluorescence signals, we first quantified fractional occupancies of each signal and their overlap, which we denote:

– p_X_ = fraction of space occupied by signal X
– p_y_ = fraction of space occupied by signal Y
– p_XY_ = fraction of space where X and Y colocalize
– p_X_ p_y_ = fractional overlap of X and Y expected under pure independence

From these metrics, we calculated a chance-corrected version of the Dice-Sörensen coefficient (*D*_*cc*_), defined as the difference between the observed Dice coefficient and its expectation under spatial independence of the two masks:

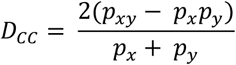

This metric is a rescaling of the covariance between masks Cov(X,Y) = *p*_*xy*_ − *p*_*x*_*p*_*y*_. D_cc_ > 0 indicates colocalization, D_cc_ < 0 indicates spatial exclusion, and D_cc_ = 0 indicates spatial independence. The Dice coefficient is used commonly to measure the overlap between binary masks, particularly in image segmentation (*63*). As an additional metric for quantifying the relationship between encoded cross-inhibition and signal overlap, we computed the colocalization coefficient (*64*):

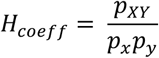

Conceptually, the colocalization coefficient is the ratio of the observed overlap between X and Y to the expected overlap if X and Y were independent. Thus, H_coeff_ > 1 indicates colocalization between the two signals, H_coeff_ < 1 indicates mutual exclusion, and H_coeff_ = 1 indicates complete independence.

### SYN-NOTCH PARAMETRIZATION EXPERIMENTS

GFP and mCherry were purified as described previously (*61*). The inhibitor-containing conditioned media was prepared as above (see “Global Inhibition of JA cells…”). For activation experiments, the ligand (GFP or mCherry) was dried overnight as the minimum volume possible covering the whole well at a concentration of 100 ug/mL. For inhibition curves, cells were seeded on 50 ug/mL GFP in a medium containing a specific dilution factor of inhibitor-containing conditioned media. Cells were seeded at 630 cells/mm^2^ (0.5X confluency). Results were analyzed via live imaging using a Keyence BZ-X710 or via Flow Cytometry (Attune) at 48 hours post-seeding. Dose–response relationships for both activator and inhibitor conditions were modeled using Hill functions. For the activator, the response was fit to 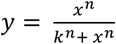; for the inhibitor, the response was fit to 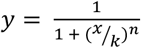, where y represents the fraction of activated cells, x denotes activator or inhibitor concentration, k is the half-maximal constant, and n is the activation or inhibition Hill coefficient. Parameters were estimated by nonlinear least-squares regression using the ‘nls’ function in R, with initial parameter values chosen to facilitate convergence and a maximum of 1000 iterations. Each experiment was first fit individually to estimate its half-maximal constant k; the mean k across experiments was computed, and each experiment’s concentrations were rescaled by its deviation from this mean so all curves shared a common inflection point. The normalized data were then pooled and fit once, and the reported Hill coefficient n was extracted from this pooled fit. This prevents inter-experiment k-variability from influencing the measured cooperativity. Fitted k and n values were extracted from these normalized models and used to plot the corresponding fitted curves. The Hill coefficients were then used to parametrize the 2D simulation models described above. The high activator cooperativity is independently supported by the ultrasensitive switch near the inflection point measured and parametrized using the same pipeline from timelapse imaging data (Fig. S6D-F). The primary measurement extracted from the imaging data was the mean fluorescence intensity over time within a segmented mask corresponding to the cell-covered area at each time point.

### CHICKEN RECOMBINATION EXPERIMENTS

L929 fibroblasts with a Wnt3a expression cassette placed downstream of synNotch activation were verified for Wnt3a expression by RNAseq using Plasmidsaurus’ ultrafast RNA-seq service. 3 days prior to recombination with chicken epidermis, L929 fibroblasts were seeded in 2% FBS medium at a density of 630 cells/mm^2^ on cell culture inserts (VWR, 353090). These cells were imaged at day 0 prior to recombination on these inserts. The day of the recombination experiment, the medium on top of the insert was aspirated just before addition of the chicken epidermis. Dorsal embryonic chicken skin was obtained from fertilized eggs at Hamilton and Hamburger stage 29-32 (*65*), when the skin is competent for feather bud formation. Dermis and epidermis were dissociated using 2% trypsin for 15-20 minutes at room temperature, then the epidermis was manually separated from the underlying dermis (*34*). A piece of dorsal epidermis was then placed onto the cell culture inserts plated with the layer of engineered fibroblasts. The resulting recombined explants were imaged at day 1. Characterization of GFP signal was performed as above, with a 2-step blurring and binarization process. Similarly, epidermal ridges were segmented from the brightfield channel using a manually defined intensity threshold, as ridges consistently appeared darker and more optically dense relative to the surrounding epidermis. Fourier transformation and area percentage were performed as above. Binary masks were skeletonized, and branch points were defined as skeleton pixels with three or more neighbors. Branch-point density was calculated as the number of such pixels divided by the total foreground area.

### LIBRARY SCREENING OF CROSS-INHIBITION CIRCUITS

Thirteen plasmids in total were constructed by restriction digestion and ligation, each placing a variable number of binding sites for the opposing circuit’s activator and a unique 70 bp barcode upstream of the gene encoding the circuit’s inhibitor. The list of constructed promoter sequences, barcodes and primers used to identify the individual promoter sequences is listed in supplementary table 4.

Six anti-GFP plasmids were generated, containing between one and six GAL4 binding sites. To produce these plasmids, a dsDNA fragment containing a single GAL4 site was PCR amplified from a 90 bp IDT oligonucleotide. The fragment and recipient plasmid were then digested with PaqCI (NEB, R0745S) to expose compatible overhangs, ligated with T4 ligase (NEB, M0202S), and transformed into NEB chemically competent 5-alpha *E. coli* (NEB, C2987H). Correct plasmids were identified by colony PCR and verified by Sanger sequencing. After completion of the initial plasmid, iterative rounds of restriction digestion and ligation into each preceding construct were used to produce additional plasmids carrying two to six GAL4 sites. To add barcodes, plasmids were opened with PaqCI and ligated with a dsDNA fragment containing a unique 70 bp barcode flanked by library-specific primer sites. These plasmids were sequence-verified using Oxford Nanopore Technologies sequencing (Plasmidsaurus).

Seven anti-mCherry plasmids were produced (pZK0024–0030). For these plasmids, the barcode and TetO binding sites were co-installed in a single restriction digestion and ligation step. Specifically, two dsDNA fragments, one containing the 70 bp barcode and the other the TetO insert, were prepared from IDT oligonucleotide templates or PCR-amplified from the TRE-anti-GFP template, respectively. Each fragment and the recipient plasmid (UAS-anti-mCherry) were digested with compatible restriction enzymes, ligated with T4 ligase, and transformed into NEB chemically competent 5-alpha competent E. coli. To construct the plasmid containing seven TetO sites, an additional TetO site was incorporated directly onto the barcode insert using BsaI-mediated type IIS assembly to circumvent a PCR amplification failure of the corresponding template region on the TRE-anti-GFP template. Correct plasmids were identified by colony PCR and verified by Sanger sequencing. These plasmids were also sequence-verified using Oxford Nanopore Technologies sequencing (Plasmidsaurus).

Constructs encoding the cross inhibition in one direction were pooled at an equimolar ratio before generation of lentiviral particles. The resulting two lentiviral libraries were titrated, and L929 cells containing a dual independent JA circuit were infected with these libraries at a multiplicity of infection (MOI) lower than 0.5. Cells expressing the fluorescent reporters for the two constructs were then sorted into single cells in 96-well plates. In parallel, bulk infected cells were FACS-sorted, lysed, PCR-amplified and NGS sequenced to verify the presence of all barcodes for both individual libraries. To genotype the resulting clones, cells were lysed using DirectPCR (Viagen, 301-C) and PCR amplified using CloneAmp (Takara, 639298), then sent to Sanger sequencing. A total of 251 clones were screened in 96-well plates, out of which 85 were characterized through from the same inhibited initial condition as described above and placed on a 2D morphospace after image processing as described above. 22 clones were selected to assess reproducibility of the patterning phenotype (see S13C). In total, 25 clones were selected for genotyping, representing 16 different genotypes, which is 29% of all possible genotype combinations (5/6 uni-directional GFP -> mCherry, 4/7 uni-directional mCherry -> GFP, 7/42 bi-directional GFP <-> mCherry). These genotyped clones were used for replicated experiments, for a total of 76 observations used to draw the correlative features between genotype and signal overlap between both circuits.

### Description of Supplementary Materials Attached to this Preprint

Supplementary Movie 1: Timelapse microscopy of L929 mouse fibroblasts engineered with a synthetic reaction-diffusion circuit, composed of synNotch-mediated juxtacrine activation and paracrine inhibition (JAPI), showing the activation of the ligand (GFP) over time.

Supplementary Movie 2: Two-dimensional simulation of a juxtacrine activator–paracrine inhibitor (JAPI) reaction–diffusion circuit, parameterized to recapitulate the phenotype of an equivalent synthetic circuit implemented in mammalian fibroblasts using synNotch-based components.

Supplementary Movie 3: Two-dimensional simulation of a juxtacrine activator–paracrine inhibitor (JAPI) reaction–diffusion circuit, parameterized to recapitulate the phenotype of an equivalent synthetic circuit implemented in mammalian fibroblasts using synNotch-based components. Here, the simulation domain size is reduced (20 × 20), mimicking experimental conditions in which fibroblasts carrying the JAPI circuit are aggregated into small spheroids generated from 500 initial cells.

Supplementary Movie 4: Time-lapse imaging of L929 mouse fibroblasts engineered with two orthogonal synthetic reaction–diffusion circuits, each composed of synNotch-mediated juxtacrine activation coupled to paracrine inhibition (JAPI). The video shows the spatiotemporal evolution of the expression of the two membrane-tethered activators, GFP and mCherry, over four days.

Table 1. Parameter sets used to generate the 1D and 2D simulation outcomes shown in the main and supplementary figures for single-circuit reaction–diffusion systems implementing either juxtacrine activator–paracrine inhibitor (JAPI) or paracrine activator–paracrine inhibitor (PAPI) architectures. The table specifies initial conditions, activator type, simulation dimensionality (1D or 2D), step number limit, and all simulation parameter values.

Table 2. Parameter sets used for 2D simulations of dual interacting reaction–diffusion circuits. For each circuit, the table lists the full parameter set of both circuits, together with additional variables specific to the dual-circuit architecture, including co-initiation conditions and cross-inhibition parameters.

Table 3. Engineered cell lines generated and used in this study, including where they appear in the figures, a description of the circuit, and the lentiviral vectors used for assembling the circuit. Vector descriptions and corresponding Addgene accession numbers, where available, are provided.

Table 4. Dual-input promoters engineered in this study, including information on the downstream the target transgene, the number and arrangement of binding sites for each of the two transcriptional inputs, the associated barcode for recognition in a pooled library assay, and the full promoter sequence.

## Supplementary Figures

**Fig. S1.**
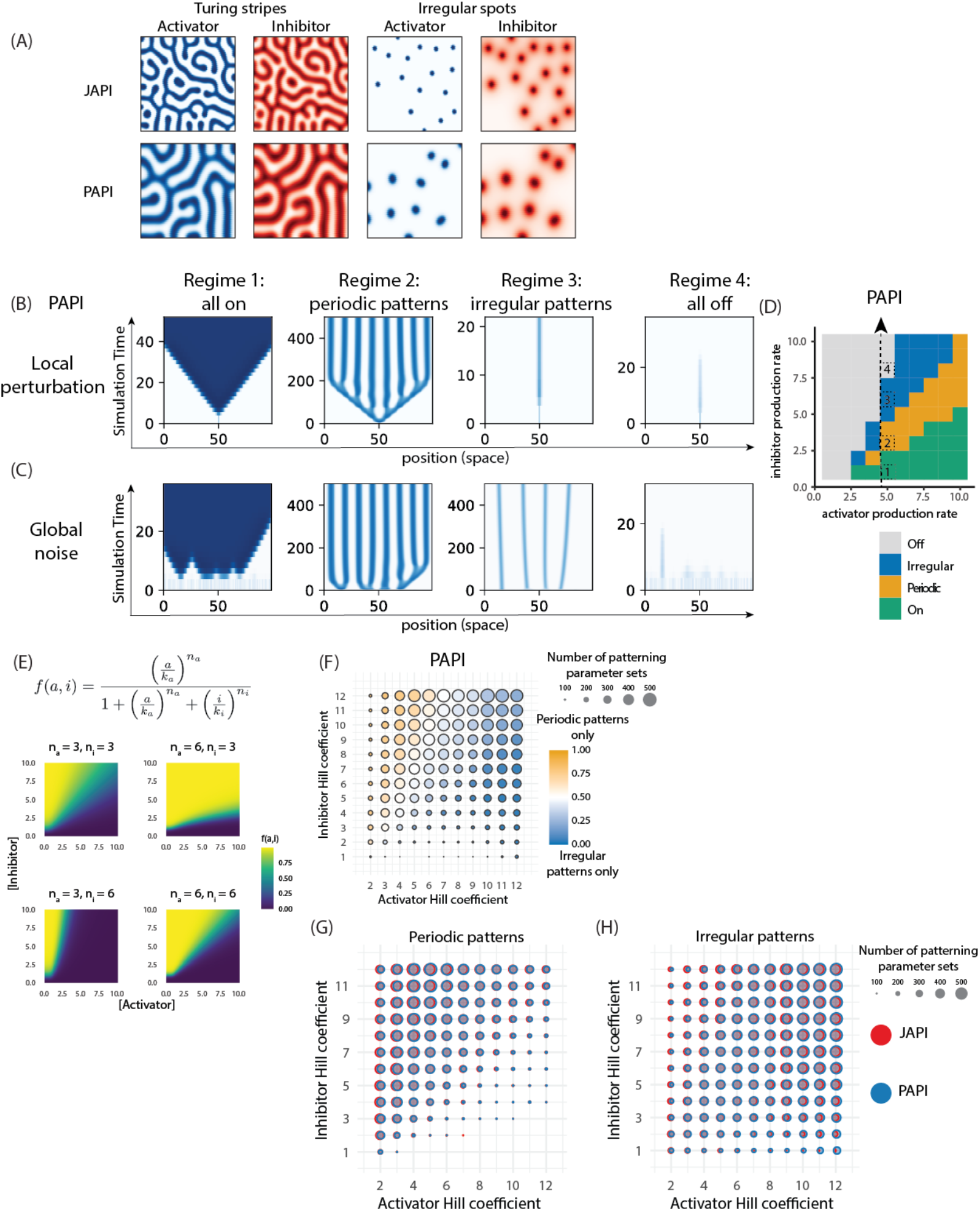
related to Fig. 1. JAPI and PAPI architectures access the same patterning regimes. (**A**) Endpoint snapshots of two-dimensional simulations for JAPI (top row) and PAPI (bottom row) circuits, shown separately for the activator (blue) and inhibitor (red) channels, in two non-homogeneous regimes labeled above each pair of columns (Turing stripes, irregular spots). Intensity indicates dimensionless activator or inhibitor concentration. Simulation setup and parameter values per each simulation is in Methods, Numerical Simulations. (**B-C**) Kymographs of one-dimensional PAPI simulations from a centrally located activator burst (B) or from spatially uniform noise (C), for parameter sets representative of the four regimes, columns labeled at the top. Blue intensity indicates dimensionless activator concentration (see Supp. Note 3 for description of dimensionless units). Initial conditions and parameter values per each simulation is in Methods, Numerical Simulations. (**D**) Phase diagram of regime classification for a PAPI circuit as a function of activator production rate (x axis) and inhibitor production rate (y axis), with the remaining dimensionless parameters held constant (n_a_ = n_i_ = 3, γ = 0.5, D = 10). Tiles are colored by regime as indicated in the legend on the figure. The four dashed boxes labeled 1 to 4 mark the parameter values used for the corresponding regimes in (B) and (C). Classification rule in Methods, Defining Patterning Regimes. (**E**) Heatmaps showing the outcome of a competitive-inhibition Hill function shown above, as a function of activator concentration (x axis) and inhibitor concentration (y axis), for four (n_a_, n_i_) combinations indicated above each heatmap. (**F**) Dotplot of PAPI patterning outcomes from a parameter sweep with 1900 per activator (x axis) and inhibitor (y axis) Hill coefficient parameter combinations, taken by independently varying the activator and inhibitor production rates and inhibitor degradation rate. Compare to Fig.1J where the same graph is shown for the JAPI architecture. Dot size is proportional to the number of parameter combinations producing patterns at each (n_a_, n_i_) pair; dot color indicates the proportion of those producing periodic patterns (orange) versus irregular patterns (blue). Sweep ranges and classification settings in Methods, Defining Patterning Regimes. (**G-H**) Dotplots of JAPI (red) and PAPI (blue) overlaid for parameter sets producing periodic patterns (G) or irregular patterns (H), projected onto the activator Hill coefficient (x axis) and inhibitor Hill coefficient (y axis). Compare to Fig. 1J and S1F, where the same dotplot is shown for each architecture separately with periodic-vs-irregular composition by color. Dot size is proportional to the number of parameter combinations producing the indicated pattern type in each architecture. The horizontal separation between paired circles was set so that their intersection area is proportional to the observed overlap between JAPI and PAPI parameter sets giving rise to the same regime. Sweep ranges and JAPI/PAPI-specific settings in Methods, Defining Patterning Regimes.

**Fig. S2.**
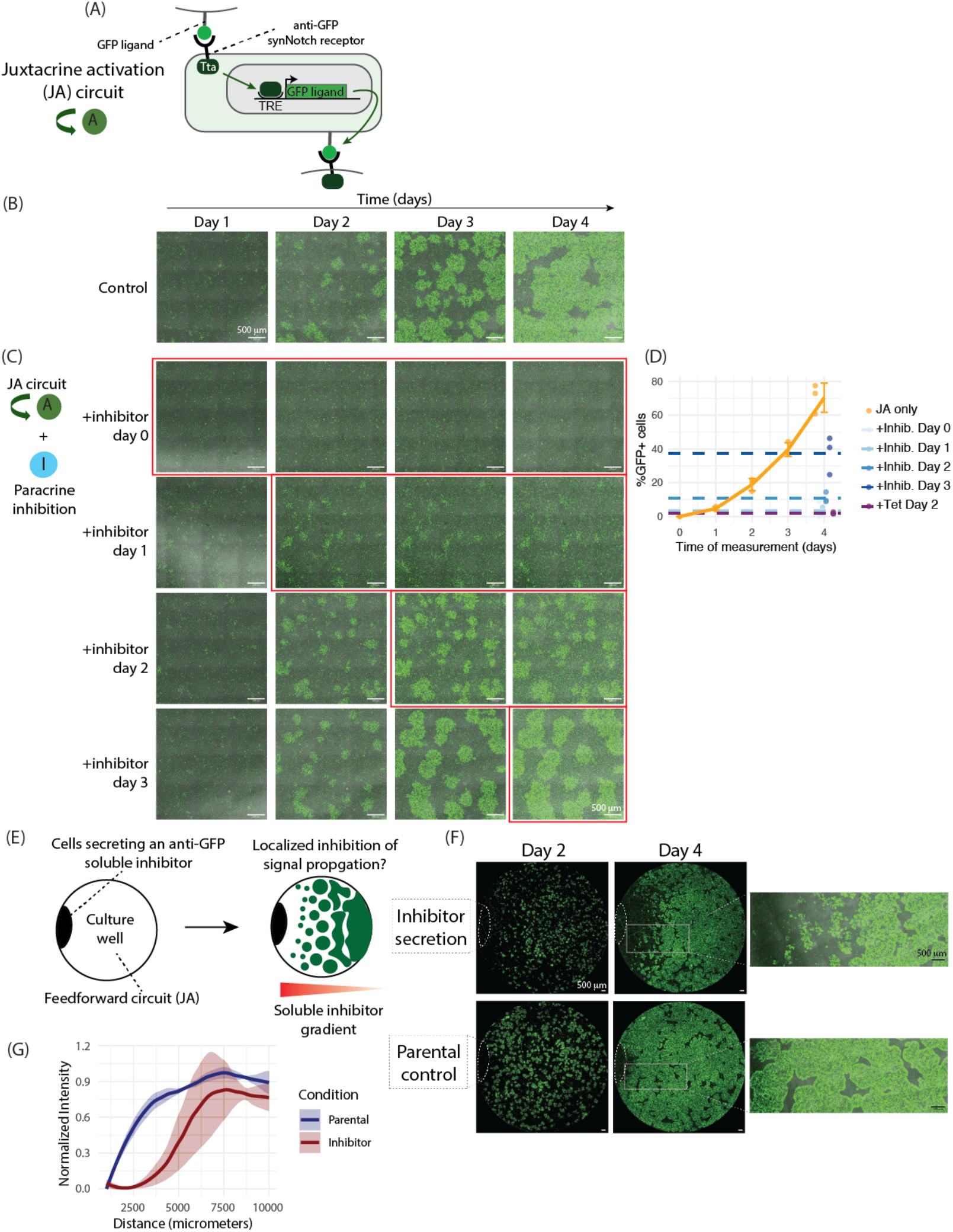
related to Fig. 2. The juxtacrine activation and paracrine inhibition branches of JAPI are functional and locally acting in isolation. (**A**) Schematic of the feedforward juxtacrine self-activation (JA) circuit. An anti-GFP synNotch receptor with an intracellular Tta transactivation domain (dark green) detects GFP ligand presented on a neighboring cell, drives transcription from a TRE promoter of a single downstream cassette encoding the membrane-tethered GFP ligand (juxtacrine activator A, green). JA circuit first introduced by Santorelli et al., Nat Comm 2024. (**B**) Fluorescence microscope images at the indicated timepoints of a timelapse experiment, showing the same field of view across time. Cells are L929 fibroblasts containing the JA circuit from (A). Green indicates activated cells (GFP signal); brightfield in grey. Initial condition: homogeneous inactivated cell lawn. Scale bar = 500 µm. (**C**) Fluorescence microscope images at the indicated timepoints of a timelapse experiment of JA cells, showing the same field of view per each row. In the four rows, a soluble anti-GFP inhibitor is added to the culture medium at the day indicated by the red box. Green indicates activated cells (GFP signal); brightfield in grey. Initial condition: homogeneous inactivated cell lawn. Scale bar = 500 µm. (**D**) Line and dot plot of the percentage of GFP-positive cells from experiments as in (C), measured by FACS. The orange line and dots show the uninhibited JA-only propagation timecourse (n = 3). Dots in shades of blue correspond to conditions where the soluble inhibitor was added at the indicated timepoint, all measured at day 4; dashed horizontal lines correspond to the average for each condition (n = 3), drawn across the full axis as a reference level. Cells inhibited with tetracycline, an inhibitor of the Tta intracellular domain of the synNotch receptor, are indicated in purple. (**E**) Schematic of the localized inhibition experimental setup. Left, condition at time zero: cells constitutively secreting soluble anti-GFP inhibitor (black) are locally seeded as a drop at the left edge of the well, and the rest of the well is plated with JA cells (white). Right, cartoon of the predicted outcome after 4 days inside the well: the inhibitor-secreting drop stays on the left edge, a graded distribution of activated GFP cells (green) across the well with fewer activated cells closer to the source. The inferred underlying inhibitor gradient is abstractly depicted as the red triangle at the bottom. (**F**) Fluorescence microscope images at the indicated timepoints of a timelapse experiment of the setup in (E); the same entire well of the experiment is shown at the two time points. Green indicates activated cells (GFP signal); brightfield in grey. Top row, the locally seeded edge cells secrete soluble anti-GFP inhibitor; bottom row, the locally seeded edge cells are non-secreting parental cells. Dotted lines indicate the position of the locally seeded edge cells. Right-most panels show a zoom of the day 4 images at the indicated positions. Scale bars = 500 µm. (**G**) Line plot of normalized GFP intensity as a function of distance from the locally seeded edge cells, for the two conditions in (F). x axis is distance in µm from the seeded cells; y axis is normalized GFP intensity. Curves are means of independent replicates with shading for standard deviation (n = 3). Image processing in Methods, Image Analysis.

**Fig. S3.**
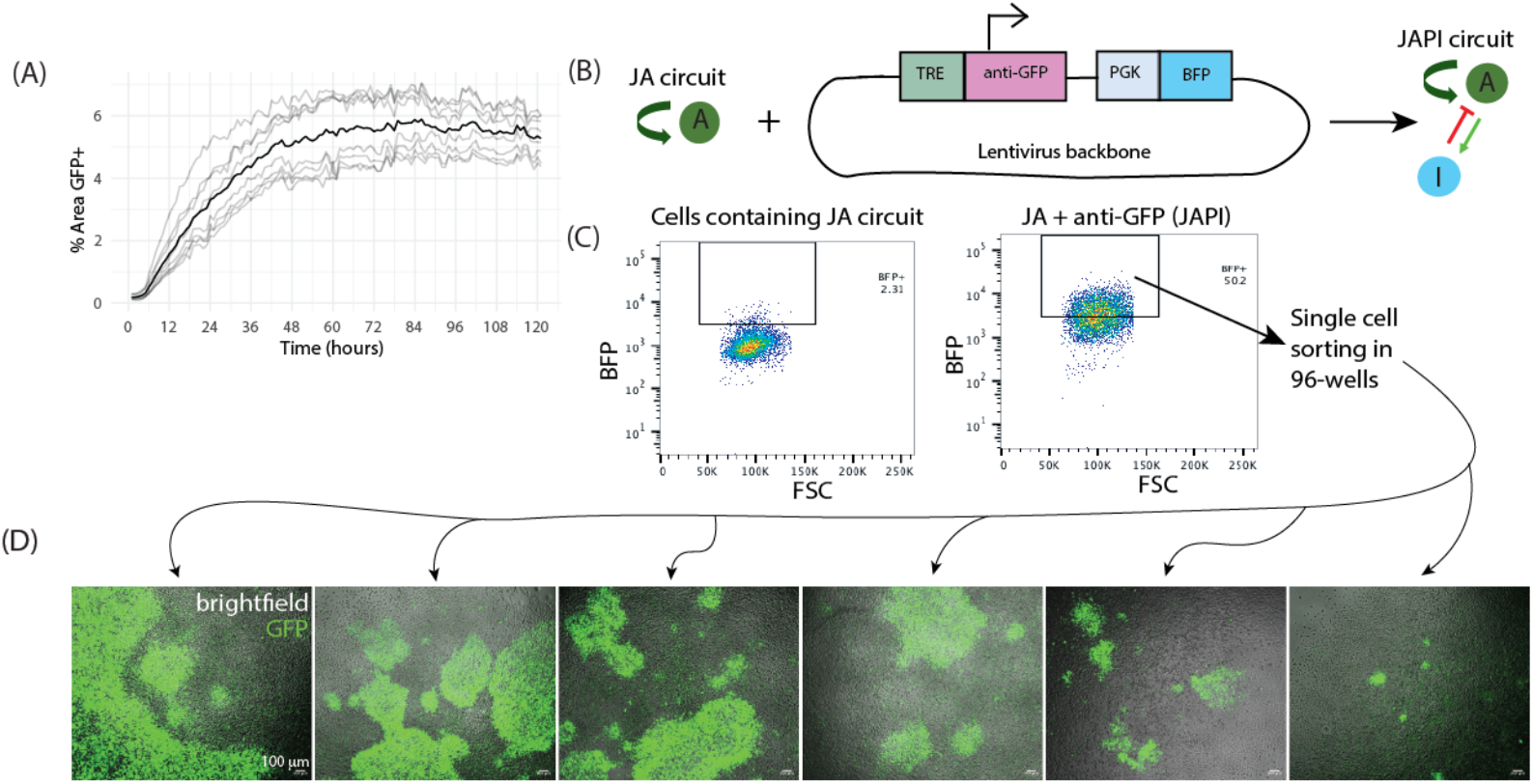
related to Fig. 2. patterning spontaneously emerges from single sorted JAPI cells. (**A**) Line plot of the area fraction covered by GFP signal over time, measured by live imaging of cells containing the JAPI circuit, from experiments such as shown in Supplemental Movie 1. Light grey curves are individual fields of view (n = 6) captured from two independent experiments; black curve is the average. Initial condition: homogeneous inactivated cell lawn. Image processing in Methods, Image Analysis. (**B**) Schematic of the lentiviral plasmid used to add the paracrine inhibitory module to cells already containing a juxtacrine activation (JA) circuit. The construct encodes the anti-GFP nanobody dimer (paracrine inhibitor) downstream of a TRE promoter, placing inhibitor expression under control of the anti-GFP-Tta synNotch receptor. A constitutive PGK promoter drives expression of a BFP reporter from the same construct, used for sorting in (C). (**C**) FACS plots showing the gating strategy for sorting BFP-positive cells, used to identify cells where the construct from (B) has been successfully integrated. x axis is forward scatter (FSC); y axis is BFP signal. Left, plot read from parental cells containing only the JA circuit (BFP-negative baseline). Right, plot read from cells from the same parental line infected with the construct from (B) (JAPI cells, BFP-positive). The black rectangle marks the gate used to sort BFP-positive cells as single cells into 96-well plates to generate clonal JAPI lines. Percentages indicate the fraction of cells inside the gate. (**D**) Fluorescence microscope images of six independent clonal JAPI cell lines derived from (C), taken directly from 96-well plates used for single-cell sorting, and imaged two weeks after. Green indicates activated cells (GFP signal); brightfield in grey. Scale bar = 100 µm.

**Fig. S4.**
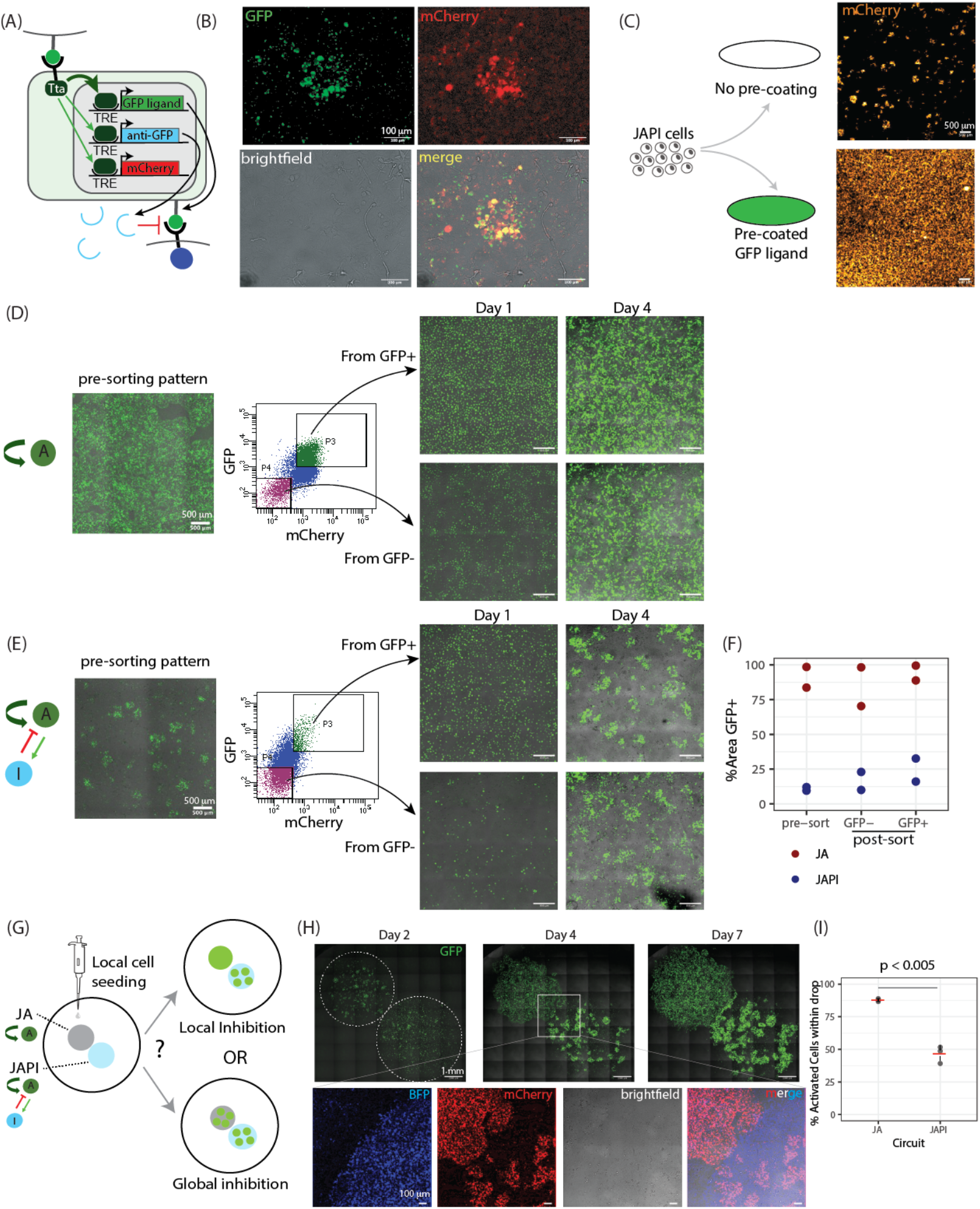
related to Fig. 2. JAPI patterning reflects synNotch activity, is dynamically maintained, and arises from locally acting inhibition. (**A**) Schematic of the JAPI circuit, modified with an additional intracellular mCherry reporter of synNotch activation. The anti-GFP-Tta synNotch receptor drives transcription from three TRE-promoted cassettes: the membrane-tethered GFP ligand (juxtacrine activator, green), the soluble anti-GFP nanobody dimer (paracrine inhibitor, light blue), and the cytoplasmic mCherry reporter (red). (**B**) Fluorescence microscope images of a clonal cell line containing the circuit from (A), showing the same field of view in four channels after 4 days of culture: GFP signal (top left, green), mCherry signal (top right, red), brightfield (bottom left, grey), and merge (bottom right). Initial condition: homogeneous inactivated cell lawn. Scale bar = 100 µm. (**C**) Test of whether JAPI cells retain the ability to activate from external ligand presentation. Left, schematic of the experimental setup: initially inactivated JAPI cells containing the circuit from (A) are plated at day zero either on a non-coated culture well (top) or on a culture well pre-coated with GFP ligand (bottom). Right, fluorescence microscope images at day four of each experiment, showing expression of the intracellular mCherry reporter (orange). Scale bar = 500 µm. (**D-E**) Pipeline of testing the reversibility of activation and inhibition for cells containing a JA (D) or JAPI (E) circuit, shown left to right as three stages: (i) the pre-sorting pattern at day 4 after onset of patterning, (ii) FACS sorting of the patterned cell population into activated (P3, GFP and mCherry double-positive) and inactivated (P4, GFP and mCherry double-negative) sub-populations, and (iii) re-plating and re-imaging of each sub-population at day 1 and day 4 after sorting. The circuit schematic (JA in D, JAPI in E) is shown at the far left. Re-plated images are labeled “From GFP+” (top, from P3) and “From GFP-” (bottom, from P4). Green indicates activated cells (GFP signal); brightfield in grey. Scale bar = 500 µm. (**F**) Dot-plot graph of the area fraction covered by GFP signal across the conditions in (D-E), for cells containing the JA circuit (red) or the JAPI circuit (blue). Each dot is an independent replicate (n = 2). Image processing in Methods, Image Analysis. (**G**) Schematic of the experimental design testing whether JAPI-mediated inhibition acts locally or at the well scale. Left, JA cells and JAPI cells are seeded as adjacent drops (gray and light blue circles respectively) in close proximity within a single well, with the rest of the well empty. Right, two possible outcomes drawn as cartoons of the well at a later timepoint: top, local inhibition, where JA cells in the JA drop activate completely and independently of the JAPI drop; bottom, global inhibition, where JAPI cells suppress JA cell activation in their drop. (**H**) Top, fluorescence microscope images at the indicated timepoints of a timelapse experiment of the setup in (G); the same field of view is shown across timepoints. Green indicates activated cells (GFP signal). Dotted lines on the day 2 image indicate the position of the two locally seeded cell drops (JA on top left, JAPI on bottom right). White rectangle on the day 4 image indicates the position of the zoom shown in the bottom panels. Bottom, zoom of the contact zone between the two cell drops at day 4, showing four channels of the same field of view: BFP (blue, marking the JAPI cells), mCherry (red, intracellular synNotch reporter in both JAPI and JA cells), brightfield (grey), and merge. (**I**) Dot-plot graph of the percentage of activated cells within each cell drop in the experiment of (H), separated by circuit (JA, JAPI),measured at day 4. Each dot is an individual replicate (n = 3); red bars indicate replicate means. p-values calculated from Welch’s Two Sample t-test.

**Fig. S5.**
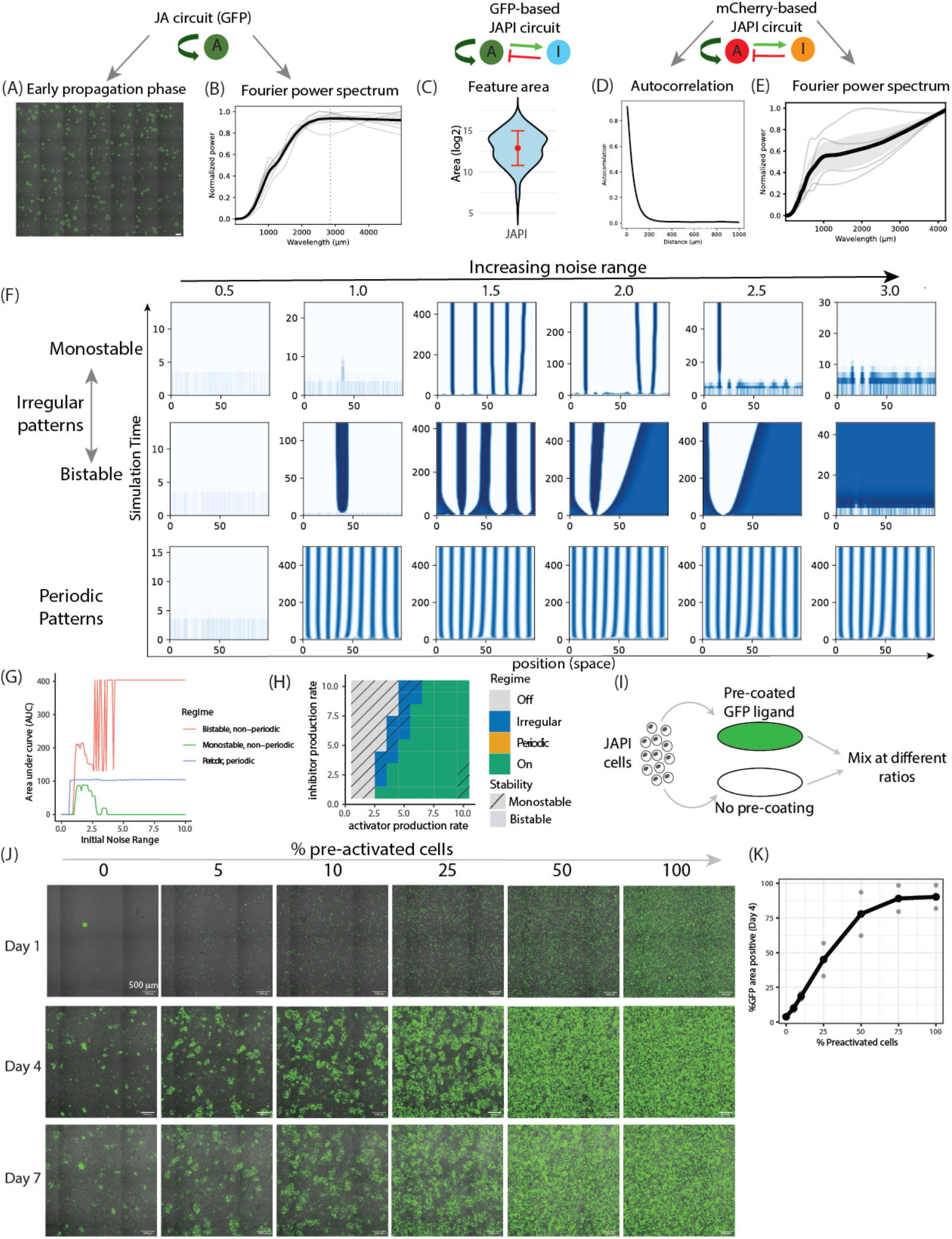
related to Fig. 2. JAPI patterns show non-periodic spatial features and initial-condition sensitivity characteristic of the irregular regime. (**A-B**) Representative microscope image and Fourier power spectrum of a cell line with the JA circuit depicted above. (A) Fluorescence microscope image of cells containing the GFP-based JA circuit at day 3, capturing the active propagation phase. Green indicates activated cells (GFP signal); brightfield in grey. Scale bar is 500 um. (B) Line plot of the Fourier power spectrum of processed GFP signal during the propagation phase, computed from images such as (A). Light grey curves are individual experiments, black curve is the average, grey shading is standard deviation (n = 5). Dotted vertical line marks an identified local maximum. Image processing in Methods, Image Analysis. (**C**) Violin plot of the distribution of activated domain areas measured from a clonal cell line containing the GFP-based JAPI circuit in the schematic above. Red dot indicates the mean, red bar indicates the standard deviation. Image processing in Methods, Image Analysis. (**D-E**) Statistical features of patterns measured at endpoint for a clonal cell line containing the mCherry-based circuit shown above and in Main Fig. 2E. (D) Line plot of the radially averaged autocorrelation of processed mCherry signal. Light grey curves are individual experiments, black curve is the average, grey shading is standard deviation (n = 4). (E) Line plot of the Fourier power spectrum of processed mCherry signal. Light grey curves are individual experiments, black curve is the average, grey shading is standard deviation (n = 4). Image processing in Methods, Image Analysis. (**F**) Grid of kymographs of one-dimensional simulations across three parameter regimes (rows, labeled at left: monostable irregular, bistable irregular, and periodic patterns) and six values of initial noise range (columns). The noise range n indicates the range of the random noise initial condition, as every cell receives an initial random activation value between 0 and n. Within each kymograph, x axis is one-dimensional position, y axis is simulation time progressing upward (note the time axis scale differs between panels to capture the relevant dynamics), and blue intensity indicates dimensionless activator concentration (see Supp. Note 3 for description of dimensionless units and stability regimes). Simulation setup and parameter values per each simulation is in Methods, Numerical Simulations. (**G**) Line plot of the area under the curve of the final activator concentration profile across position as a function of initial noise range, for the three parameter regimes shown in (F). Curves are color-coded by regime as indicated in the figure legend. Simulation setup in Methods, Numerical Simulations. (**H**) Phase diagram of regime classification for a JAPI circuit on the activator/inhibitor production rate axes, with an overlaid hatching pattern indicating monostable (single hatching) vs bistable (no hatching) regions. Tiles are colored by regime as indicated in the legend on the figure. Note that, for this parameter set, no combination of production rates generates periodic patterns, and that the irregular patterning regime spans both monostable and bistable parameter sets. Classification rule and bistability determination in Methods, Defining Patterning Regimes. (**I**) Schematic of the experimental design testing the sensitivity of JAPI patterning to initial conditions. Cells from a pre-coated GFP-ligand well (activated JAPI cells, green) and cells from a non-coated well (inactivated JAPI cells, white) are mixed at varying ratios and re-plated. (**J**) Fluorescence microscope images at the indicated timepoints of a timelapse experiment of the setup in (I); each column shows a different starting fraction of pre-activated cells; each row shows the same field of view across time. Green indicates activated cells (GFP signal); brightfield in grey. Scale bar = 500 µm. (**K**) Dot plot of the percentage of GFP-positive area at day 4 as a function of the percentage of pre-activated cells at time zero, from experiments as in (J). Each dot is an individual replicate (n = 2); black curve connects the means. Image processing in Methods, Image Analysis.

**Fig. S6.**
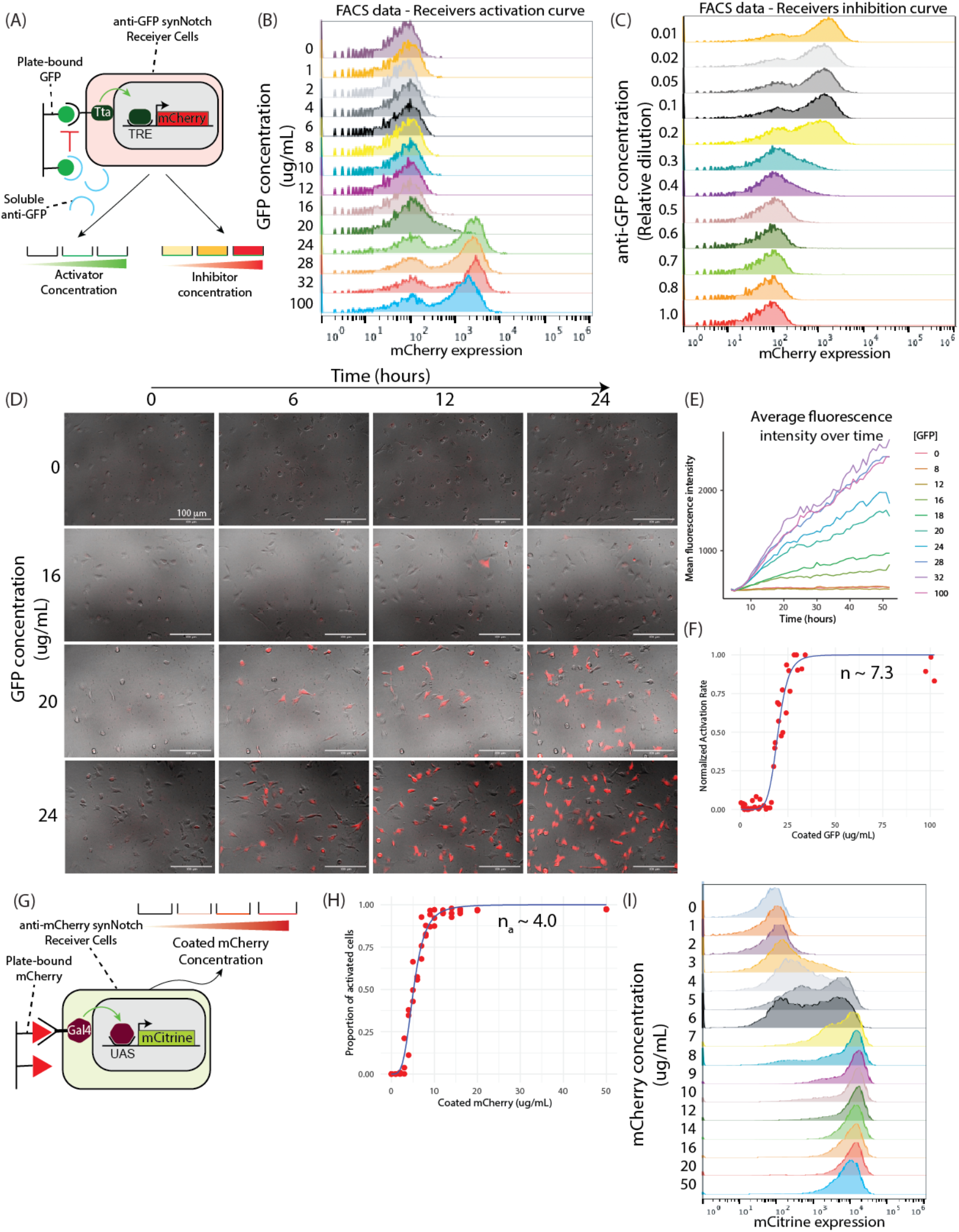
related to Fig. 2. synNotch receptors show non-linear activation and inhibition, with Hill coefficients depending on the receptor architecture. (**A**) Schematic of the experimental setup for measuring synNotch dose-response curves. A receiver cell line constitutively expressing an anti-GFP-Tta synNotch receptor with a downstream TRE-promoted mCherry reporter is plated at low confluency under two conditions: left, on a culture well coated with a varying concentration of GFP ligand (activator) to measure activation; right, in culture medium containing a varying concentration of soluble anti-GFP inhibitor in the presence of constant activator concentration, to measure inhibition. In both cases, the output of activation is read as mCherry expression. (**B**) FACS histograms of mCherry expression in receiver cells from (A) 48 hours after plating on increasing concentrations of plate-bound GFP ligand (rows, labeled at left). (**C**) FACS histograms of mCherry expression in receiver cells from (A) 48 hours after plating on a constant concentration of plate-bound GFP ligand and increasing concentrations of soluble anti-GFP inhibitor (rows, labeled at left, in relative dilution of inhibitor-containing conditioned media). (**D**) Fluorescence microscope images at the indicated timepoints of a timelapse experiment of receiver cells from (A) plated on increasing concentrations of plate-bound GFP ligand (rows, labeled at left). Red indicates synNotch-activated cells (mCherry reporter); brightfield in grey. Scale bar = 500 µm. (**E**) Line plot of mean mCherry fluorescence intensity over time from the experiments in (D), for each plate-bound GFP concentration (curves color-coded by concentration as indicated in the figure legend). Curves indicate the means of n = 3 independent experiments. Image processing in Methods, Image Analysis. (**F**) Scatter plot with fitted Hill curve of normalized activation rate as a function of plate-bound GFP concentration, computed from the slopes of the curves in (E). Red dots are individual experimental data points from n = 3 independent replicates; blue curve is the fitted activating Hill function. The fitted Hill coefficient is indicated in the figure. Fitting procedure in Methods, synNotch parametrization experiments. (**G**) Schematic of the experimental setup for measuring dose-response curves of a second synNotch receptor with an orthogonal architecture constructed with a mCherry ligand binding domain and Gal4 intracellular transactivation domain. A receiver cell line constitutively expressing an anti-mCherry-Gal4 synNotch receptor with a downstream UAS-promoted mCitrine reporter is plated on a culture well coated with mCherry ligand (varying concentration). The output of activation is read as mCitrine expression. (**H**) Scatter plot with fitted Hill curve of the proportion of activated receiver cells from (G) as a function of plate-bound mCherry concentration, computed from FACS data such as shown in (I). Red dots are individual experimental data points from n = 3 independent experiments; blue curve is the fitted activating Hill function. The fitted Hill coefficient is indicated in the figure. Fitting in Methods, synNotch parametrization experiments. (**I**) FACS histograms of mCitrine expression in receiver cells from (G) plated on increasing concentrations of plate-bound mCherry ligand (rows, labeled at left, in µg/mL).

**Fig. S7.**
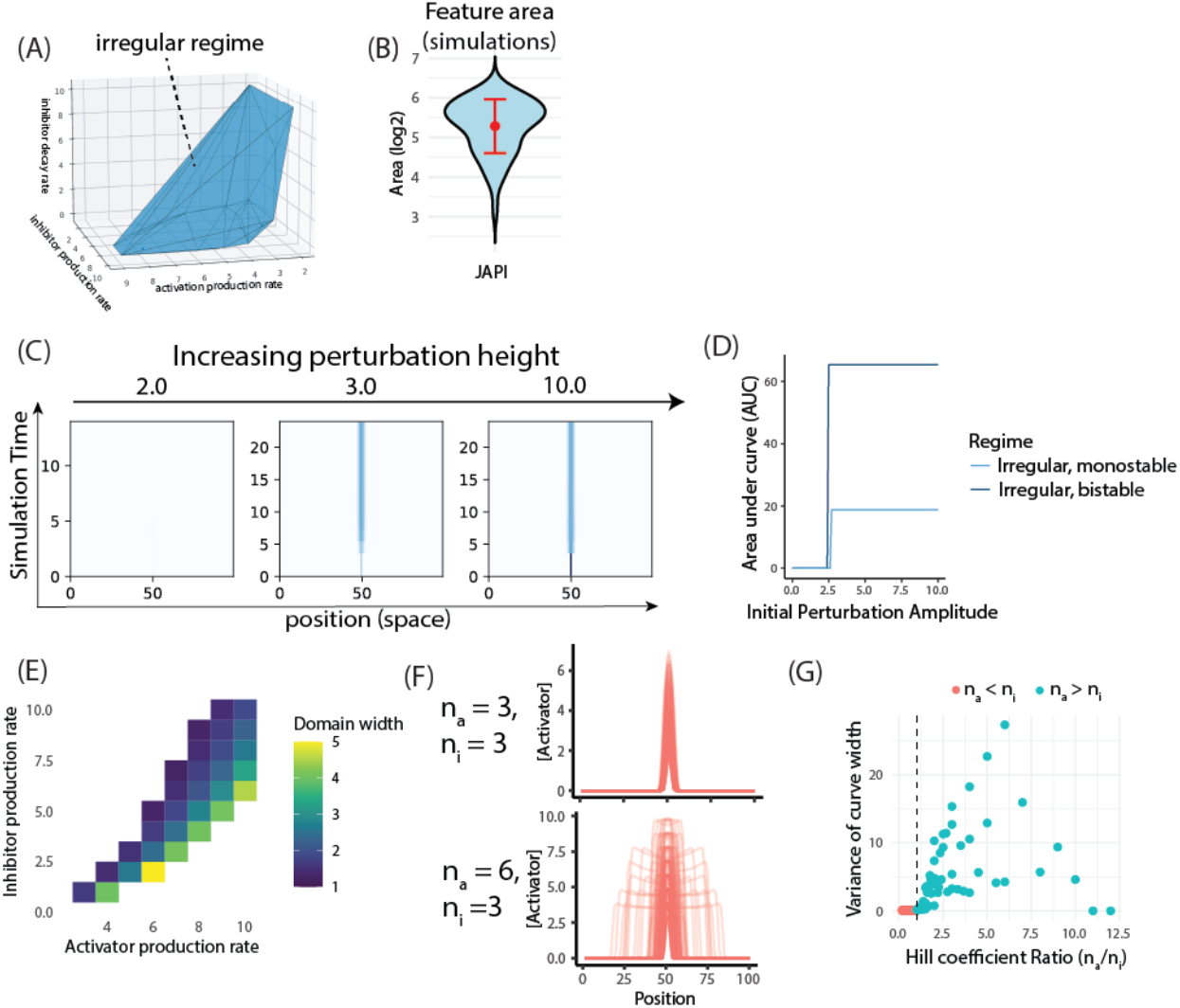
related to Fig. 2. Domain size in the irregular regime is set by activator-inhibitor balance and depends on relative activator and inhibitor Hill cooperativity. (**A**) 3D plot showing the volume of parameter space within the irregular regime for a JAPI circuit with parametrized Hill coefficients (n_a_ = 10, n_i_ = 4). Axes are the three remaining non-dimensionalized parameters: activator production rate β_a_, inhibitor production rate β_i_, and inhibitor degradation rate γ. Blue points mark parameter combinations classified as irregular; non-blue points mark homogeneous regimes (all on or all off). Classification rule in Methods, Defining Patterning Regimes. (**B**) Violin plot of the distribution of feature area from parametrized JAPI simulations (n_a_ = 10, n_i_ = 4), initiated from continuous stochastic activation bursts. Red dot indicates the mean, red bar indicates the standard deviation (n = 3 replicated simulations). Compare to S5A, which shows the same metric measured experimentally. Simulation setup in Methods, Numerical Simulations. (**C**) Kymographs of one-dimensional JAPI simulations initiated from a centrally located activator burst of increasing amplitude (columns), for a parameter set within the irregular regime. Blue intensity indicates dimensionless activator concentration (see Supp. Note 3 for description of dimensionless units). Simulation setup and parameter values per each simulation is in Methods, Numerical Simulations. (**D**) Line plot of the area under the curve of the final activator concentration profile across position as a function of initial perturbation amplitude, for two parameter sets within the irregular regime: monostable (light blue) and bistable (dark blue). Simulation setup in Methods, Numerical Simulations. (**E**) Heatmap of the width of the stable activated domain emerging from a centrally located activation burst, as a function of activator production rate (x axis) and inhibitor production rate (y axis), with the remaining non-dimensionalized parameters held constant (n_a_ = 6, n_i_ = 3, γ = 0.1) within the irregular regime. Color represents domain width quantified by the standard deviation of the activator spatial distribution, expressed in cell units (see colorbar). Simulation setup and parameter values in Methods, Numerical Simulations. (**F**) Line plot of the activator concentration profile across one-dimensional position, overlaying the outcome of ∼160 simulations from a centrally located activation burst across varying activator and inhibitor production rates and inhibitor degradation rate, for two pairs of Hill coefficients shown separately. Each light red curve is an individual parameter combination outcome; the spread illustrates the parameter sensitivity of domain width. Simulation setup and parameter values in Methods, Numerical Simulations. (**G**) Scatter plot of the variance of activator peak width across parameter sets within the irregular regime as a function of the Hill coefficient ratio n_a_ / n_i_ (n = 12 x 12 = 144 total points). Red dots, parameter sets with n_a_ < n_i_ ; teal dots, parameter sets with n_a_ > n_i_ . Dashed vertical line marks the n_a_ / n_i_ = 1 boundary. Simulation setup and parameter values in Methods, Numerical Simulations.

**Fig. S8.**
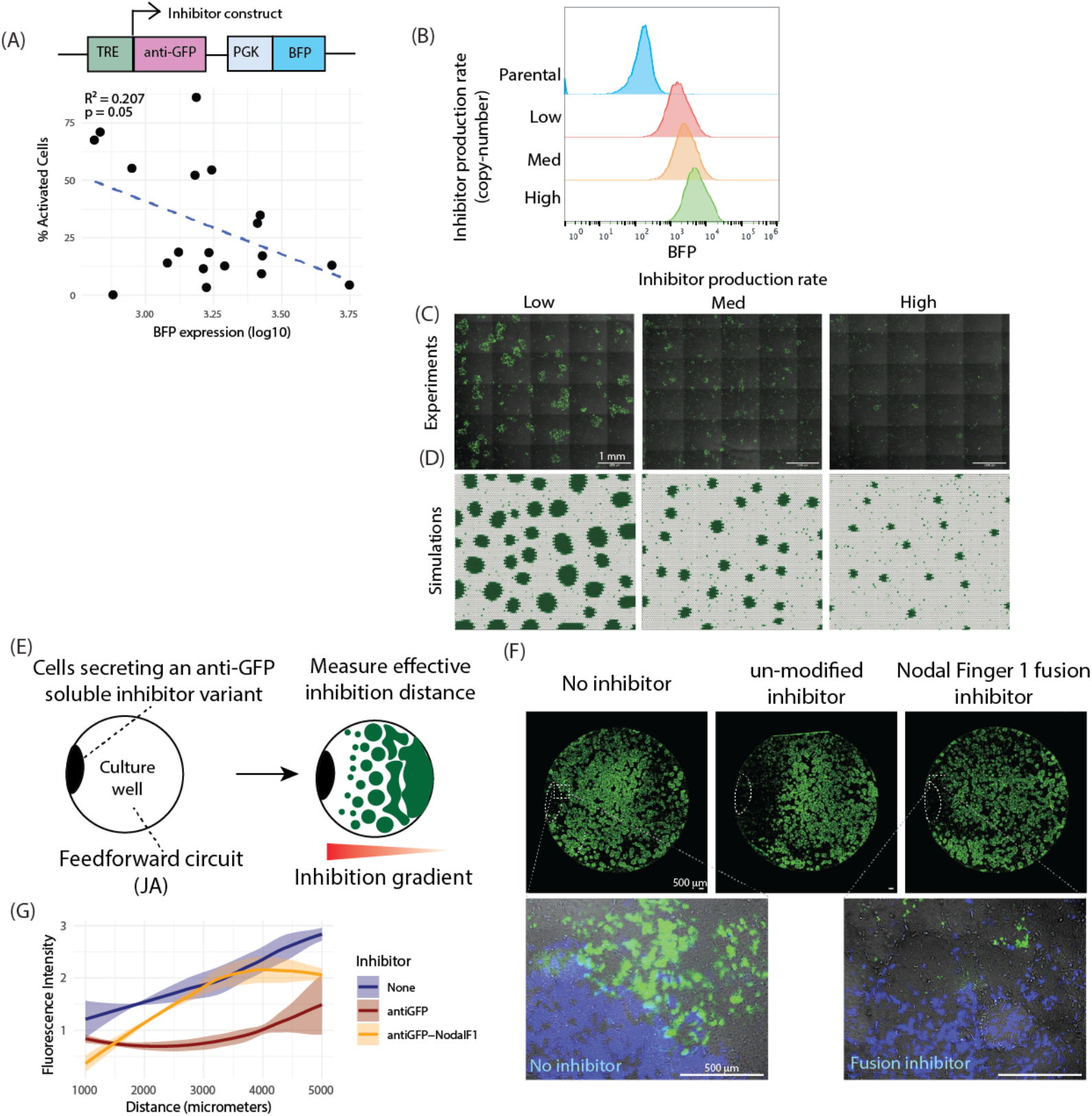
related to Fig. 3. Validation of inhibitor production rate and diffusion length as valid parameter knobs for tuning domain size in JAPI-mediated irregular patterning. (**A**) Top, schematic of the lentiviral construct used to add downstream activation of a paracrine inhibitor in a JAPI circuit. Bottom, scatter plot with linear regression of the percentage of activated cells as a function of BFP expression (log10 scale). Each dot is a measurement made from an individual clonal line (n = 19); dashed line is the best-fit linear regression. R^2^ and p-value indicated in the figure. (**B**) FACS histograms of BFP expression in three polyclonal cell lines bulk-sorted from a population of JAPI cells based on distinct BFP expression profiles (rows, labeled at left). (**C**) Fluorescence microscope endpoint images at day 4 of the three bulk-sorted JAPI cell lines from (B), with increasing inhibitor production rate β_i_ (left to right). Green indicates activated cells (GFP signal); brightfield in grey. Initial condition: homogeneous inactivated cell lawn. Scale bar = 500 µm. Quantification of these data is shown in Fig. 3C. (**D**) Endpoint snapshots of two-dimensional JAPI simulations parametrized with the Hill coefficients from Fig. 2J-K (n_a_ = 10, n_i_ = 4), with three increasing values of the inhibitor production rate β_i_ (left to right). Green intensity indicates dimensionless activator concentration (see Supp. Note 3 for description of dimensionless units). Simulation setup and parameter values per each simulation is in Methods, Numerical Simulations. Quantification of these data is shown in Main Fig. 3D. (**E**) Schematic of the experimental setup for measuring the effective diffusion length of paracrine anti-GFP inhibitor variants. Left, cells constitutively secreting a soluble anti-GFP inhibitor variant are locally seeded as a drop at the left edge of the well, and the rest of the well is plated with JA cells. Right, cartoon of the expected outcome inside the well at a later timepoint: the inhibitor-secreting drop on the left edge, a graded distribution of activated GFP cells (green) across the well with fewer activated cells closer to the source. The inferred underlying inhibitor gradient is abstractly depicted as the red triangle at the bottom. (**F**) Fluorescence microscope endpoint images at day 4 of the setup in (E), for three conditions in columns (labeled at top). No inhibitor corresponds to a condition where parental cells were locally seeded. Top row, full-well view; bottom row, zoom of the indicated regions showing the boundary between the locally seeded cells (BFP, blue, expressed in all three locally seeded lines as a tracking marker) and activation of the JA cells (GFP, green). Scale bar = 500 µm. (**G**) Line plot of normalized fluorescence intensity of the JA-cell GFP signal as a function of distance from the locally seeded drop, for the three conditions in (F). Curves are means of independent replicates with shading for standard deviation (n = 3). Image processing in Methods, Image Analysis.

**Fig. S9.**
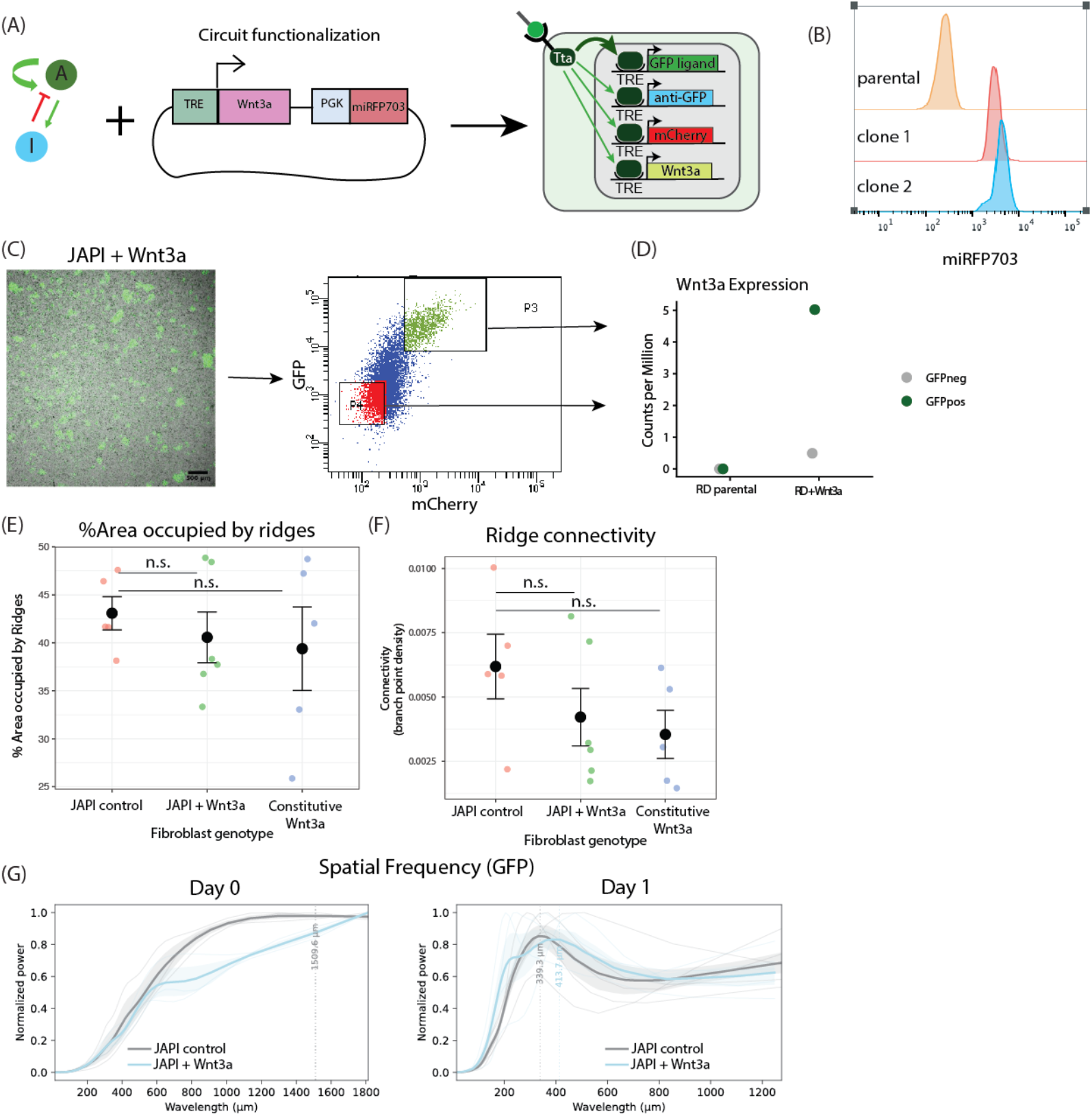
related to Fig. 3. Functionalization of a JAPI circuit for downstream patterned production of Wnt3a, and characterization of their interaction with dorsal chicken embryonic epidermis. (**A**) Schematic of the functionalization of a JAPI circuit by addition of a Wnt3a expression cassette downstream of synNotch activation, showing the lentiviral construct added to the circuit. Right, the resulting functionalized JAPI circuit with four cassettes downstream of synNotch receptor activation. (**B**) FACS histograms of miRFP703 expression in three cell populations (rows, labeled at left): parental JAPI cells, and two clonal lines generated from infection with the construct in (A). (**C**) Left, fluorescence microscope image and FACS gating strategy for transcriptomic analysis of clonal cells containing a functionalized JAPI circuit. Green indicates activated cells (GFP signal); brightfield in grey. Scale bar = 500 µm. Right, FACS plot showing the gating used to sort activated (P3, GFP and mCherry double-positive) and inactivated (P5, GFP and mCherry double-negative) cell populations for transcriptomic profiling by RNA-seq. (**D**) Dot plot of Wnt3a transcript expression measured by RNA-seq, for the two sorted populations from (C) (“RD + Wnt3a”) in comparison to sorted populations from parental JAPI cells (no Wnt3a transgene, “RD parental”). Each dot is an individual replicate (n = 1). (**E**) Dot plot of the percentage of epithelial area covered by ridges 24 hours after recombination, separated by underlying engineered fibroblast genotype. Each dot is an individual replicate; black bars indicate mean and standard deviation (n = 5). p-values (n.s. = not significant) calculated from Welch’s Two Sample t-test. Image processing in Methods, Chicken recombination experiments. (**F**) Scatter plot of the ridge connectivity (branch-point density on a skeletonized binary image) of the epithelial ridges 24 hours after recombination, separated by underlying engineered fibroblast genotype. Each dot is an individual replicate (n = 5); black bars indicate mean and standard deviation. p-values (n.s. = not significant) calculated from Welch’s Two Sample t-test. Skeletonization procedure in Methods, Chicken recombination experiments. (**G**) Line plot of the Fourier power spectrum of the GFP signal of JAPI control and JAPI + Wnt3a engineered fibroblasts, shown at day 0 (left, before recombination) and day 1 (right, 24 hours after recombination with embryonic dorsal chicken epithelium). Light grey curves are individual experiments, dark curves are the average for each condition, shading is standard deviation (n = 5 for JAPI control, n = 3 for PAI + Wnt3a). Dotted vertical lines mark identified local maxima. Image processing in Methods, Chicken recombination experiments.

**Fig. S10.**
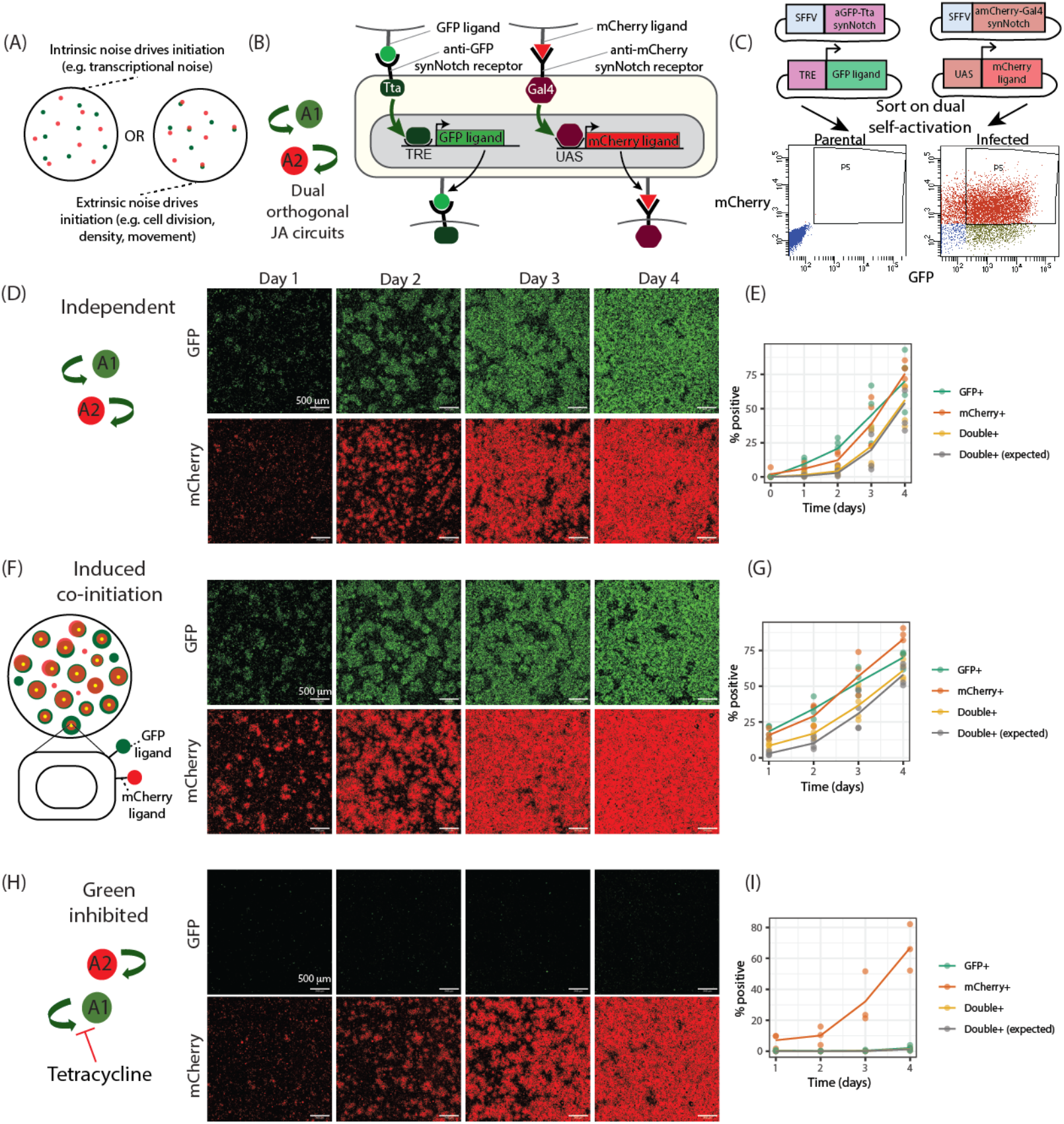
related to Fig. 4. Construction and characterization of dual orthogonal JA circuits in a single cell line. **(A)** Schematic of two possible outcomes for the initiation of dual orthogonal synNotch-based juxtacrine activation (JA) circuits, drawn as cartoons of a cell well at an early timepoint. Left, intrinsic noise drives initiation: each circuit initiates independently from transcriptional noise (green and red dots in distinct cells). Right, extrinsic noise drives initiation: shared cellular events (cell division, density, movement) co-initiate signal propagation of both circuits in the same cells. **(B)** Schematic of a circuit encoding dual orthogonal synNotch-mediated juxtacrine self-activation (JA), without paracrine inhibition. The anti-GFP-Tta synNotch receptor drives TRE-promoted GFP ligand; the anti-mCherry-Gal4 synNotch receptor drives UAS-promoted mCherry ligand. The two circuits are theoretically completely orthogonal. (**C**) Infection strategy for generating the dual-JA cell line. Top, schematic of the four lentiviral constructs used: two SFFV-promoted synNotch receptors (anti-GFP-Tta, anti-mCherry-Gal4) and two response-promoted ligands (TRE-GFP ligand, UAS-mCherry ligand). Bottom, FACS plots showing the sorting strategy: parental cells (left, baseline) and infected cells (right, post-infection); the P5 gate marks cells positive for both GFP and mCherry, used for single-cell sorting to generate clonal dual-JA lines. (**D**) Fluorescence microscope images at the indicated timepoints of a timelapse experiment of cells containing the dual-JA circuit from (B); the same field of view is shown across timepoints. Scale bar = 500 µm. The merge of the two channels is shown in Fig. 4C. (**E**) Plot of the percentage of single and dual activated cells over time, measured by FACS from the experiment in (D). Curves color-coded by population, with grey representing double positive cells expected by independent probability. Each dot is an individual replicate (n = 4); lines connect replicate means. (**F**) Left, schematic of the induced co-initiation experiment: dual-JA cells from (B) are mixed with 1% cells constitutively expressing both GFP and mCherry ligands (dual senders). Right, fluorescence microscope images at the indicated timepoints of a timelapse experiment of this setup; the same field of view is shown across timepoints. Scale bar = 500 µm. The merge of the two channels is shown in Fig. 4E. (**G**) Plot of the percentage of single and dual activated cells over time, measured by FACS from the experiment in (F) (n = 4). Curves color-coded by population as in (E). (**H**) Fluorescence microscope images at the indicated timepoints of a timelapse experiment of the dual-JA cells from (D) in medium containing tetracycline, which inhibits the Tta domain of the anti-GFP synNotch receptor and selectively blocks the GFP-based circuit; the same field of view is shown across timepoints. Scale bar = 500 µm. The merge of the two channels is shown in Fig. 4G. (**I**) Line plot of the percentage of GFP-positive, mCherry-positive, and double-positive cells over time, measured by FACS from the experiment in (H) (n = 3). Curves color-coded by population as in (E).

**Fig. S11.**
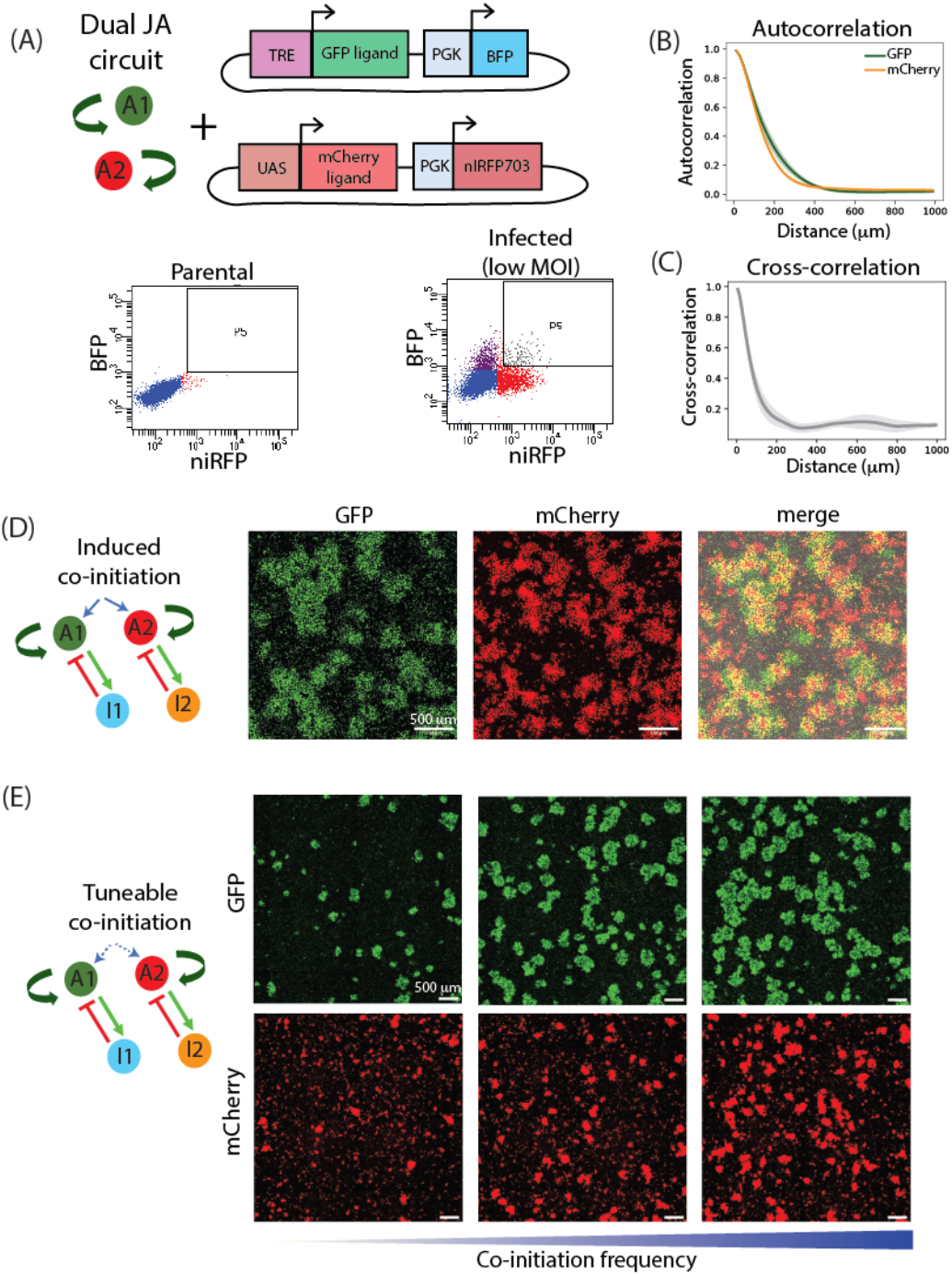
related to Fig. 4. Construction and characterization of dual orthogonal JAPI circuits in a single cell line. (**A**) Top, schematic of the implementation of dual orthogonal JAPI circuits by adding two inducible inhibitor transgenes to the dual-JA cell line from (S11B). Left, the parental dual-JA circuit. Right, the two added lentiviral constructs. Bottom, FACS sorting strategy for dual-JA cells receiving both inhibitor modules. The P5 gate marks cells positive for both BFP and niRFP, sorted to generate clonal dual-JAPI lines. (**B**) Line plot of the radially averaged autocorrelation of GFP signal (green) and mCherry signal (orange) from patterns generated by a dual-JAPI cell line. Curves are means with shading for standard deviation across replicates (n = 3). Image processing in Methods, Image Analysis. (**C**) Line plot of the radially averaged cross-correlation between GFP and mCherry signals from patterns generated by a dual-JAPI cell line. Curve is the mean with shading for standard deviation across replicates (n = 3). Image processing in Methods, Image Analysis. (**D**) Single-channel fluorescence microscope endpoint images at day 4 of cells containing dual orthogonal JAPI circuits mixed with 1% of cells constitutively expressing ligand GFP and ligand mCherry (dual senders). (**E**) Single-channel fluorescence microscope endpoint images at day 4 of cells containing dual orthogonal JAPI circuits mixed with increasing numbers of dual senders (left to right). Each image shows the separate channels corresponding to the merged images shown in Fig 4N. Scale bar = 500 µm.

**Fig. S12.**
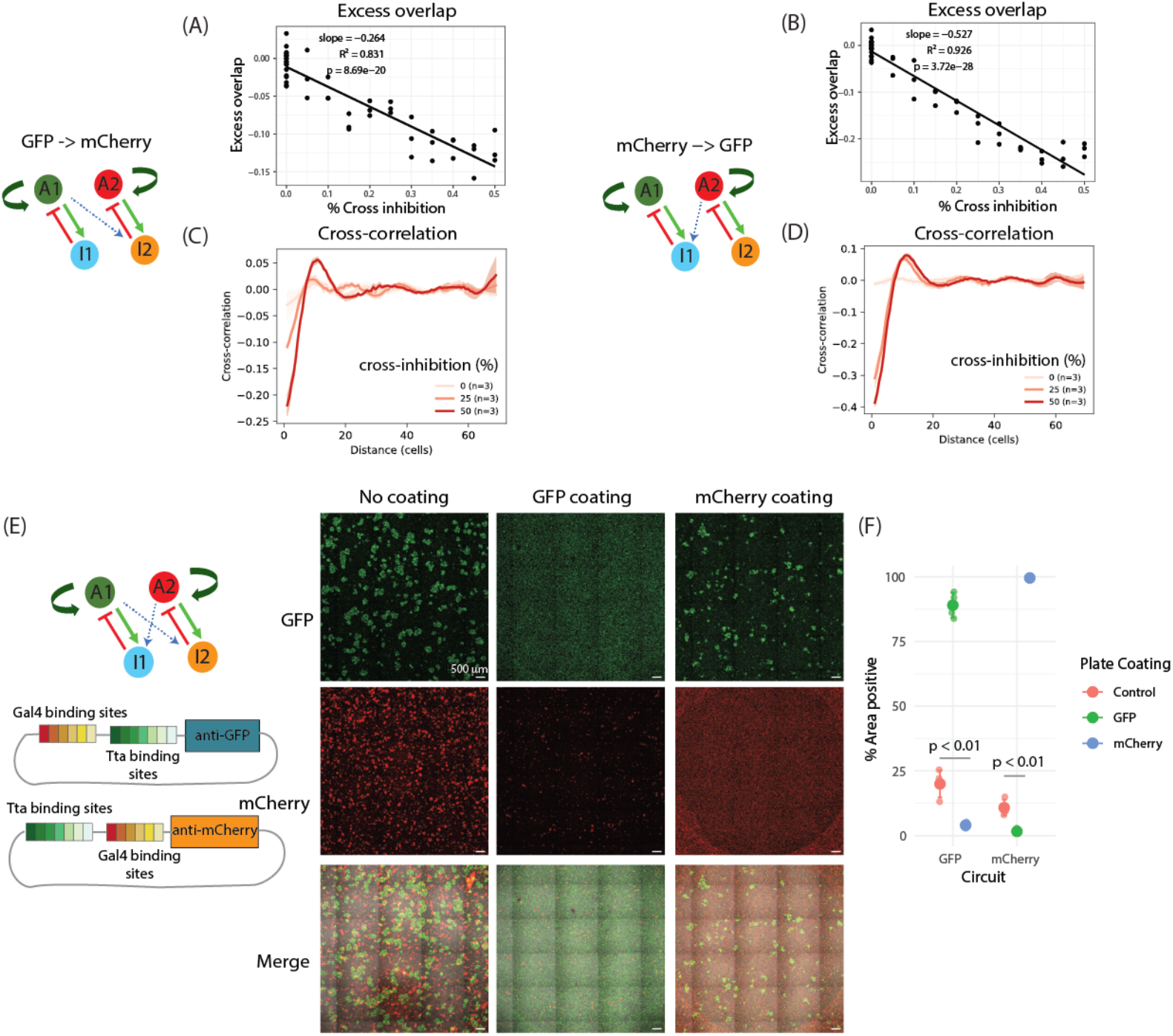
related to Fig. 5. Characterization of uni-directional and bi-directional cross-inhibition in dual-JAPI circuits. (**A-D**) Simulated uni-directional cross-inhibition. Left of the results panels, schematic of the dual JAPI circuit with uni-directional cross-inhibition GFP → mCherry, indicated by a dashed blue arrow. (**A**) Scatter plot with linear regression of excess overlap as a function of cross-inhibition strength, computed from two-dimensional simulation. Each dot is an individual simulation (n = 3 per condition); black line is the best-fit linear regression. Slope, R^2^, and p-value indicated in the figure. Simulation setup and parameter values in Methods, Numerical Simulations. (**B**) Same as (A) for uni-directional cross-inhibition in the opposite direction (mCherry → GFP). (**C**) Line plot of the radially averaged cross-correlation between GFP and mCherry signals, for three representative cross-inhibition strengths from the simulations in (A) (curves color-coded by strength as indicated in the figure legend). Curves are means with shading for standard deviation across replicates (n = 3 per condition). Image processing in Methods, Image Analysis. (**D**) Same as (C) at three matching representative cross-inhibition strengths for uni-directional cross-inhibition in the opposite direction (mCherry → GFP). (**E**) Left, schematic of the bi-directional cross-inhibiting dual-JAPI circuit design. The promoter of each paracrine inhibitor cassette is built as a tandem of the two synNotch-inducible promoter elements (TRE and UAS), with the binding sites of the cross-inhibiting synNotch placed further from the transcription start site than the cognate sites. Right, fluorescence microscope endpoint images for cells plated on culture wells with three coating conditions (no coating, GFP coating, mCherry coating), in order to force activation of one circuit and assess the consequence of cross-inhibition on the opposing signal. (**F**) Dot plot of the percentage of area positive for each fluorescent signal, separated by coating condition and fluorescent circuit reported. Dot colors encode plate coating condition. Each dot is an individual replicate (n = 3). p-values calculated using Welch’s Two Sample t-test.

**Fig. S13.**
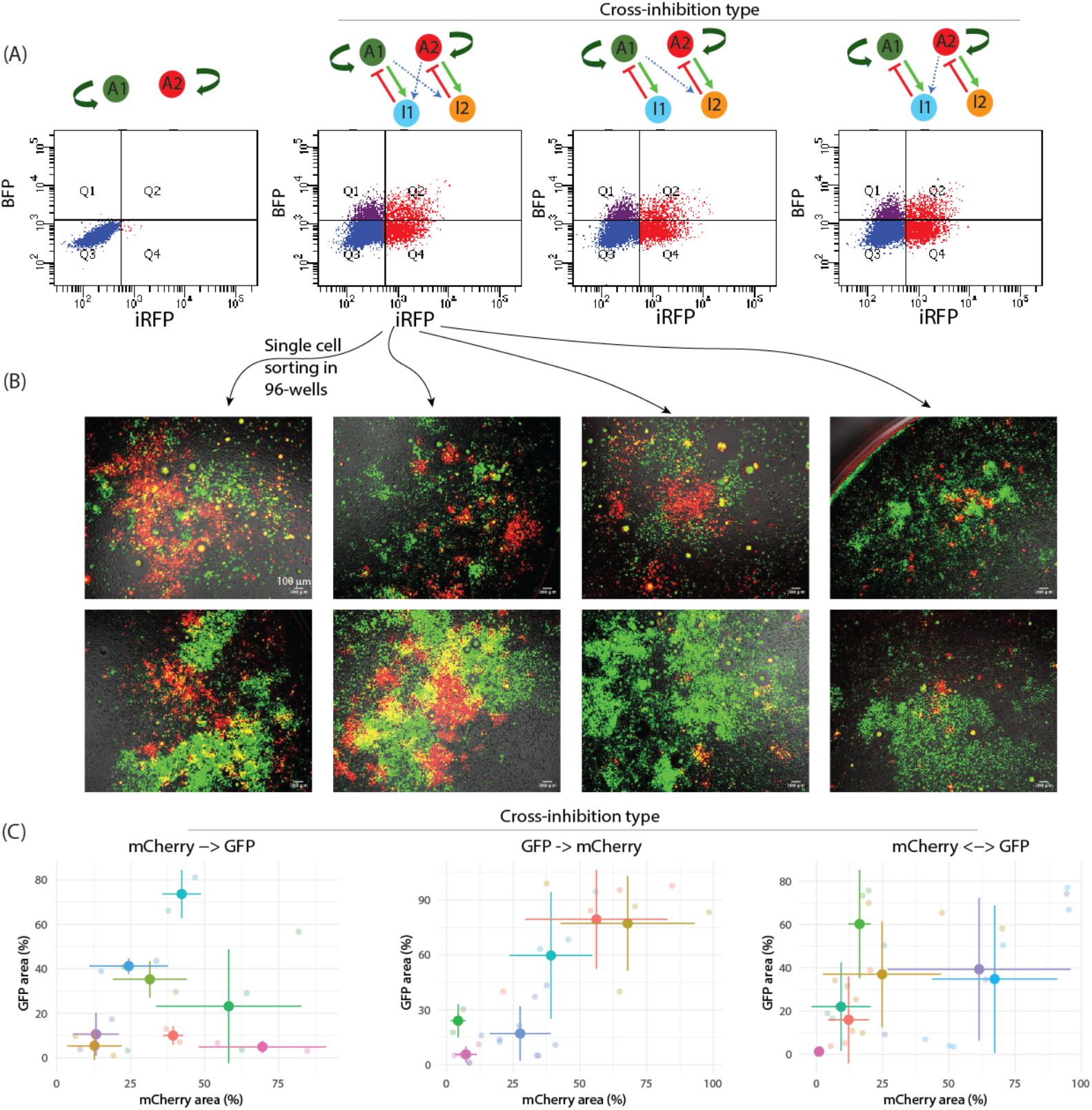
related to Fig. 5. Clonal characterization of the cross-inhibiting dual-JAPI library. (**A**) FACS sorting strategy for generating clonal dual-JAPI cell lines with bi-directional or uni-directional cross-inhibition. Four plots are shown, where quadrant Q2 was used to generate clonal populations through single-cell sorting of dual reporter-positive cells in 96-well plates. Left, parental dual-JA cells (BFP-negative, iRFP-negative baseline). The next three plots show dual-JA cells infected at low MOI (MOI < 0.5) with three types of cross-inhibition library constructs (schematic above each plot): bi-directional cross-inhibition, uni-directional GFP → mCherry, and uni-directional mCherry → GFP. (**B**) Fluorescence microscope images of eight representative clonal cross-inhibiting dual-JAPI cell lines derived from the sorting in (A), taken in the same 96-well well around two weeks after single-cell sorting. Green indicates activated cells (GFP signal); red indicates activated cells (mCherry signal); brightfield in grey. Scale bar = 100 µm. (**C**) Scatter plot of the percentage of area covered by mCherry signal (x axis) versus the percentage of area covered by GFP signal (y axis), measured at day 4 across clonal dual-JAPI cell lines, separated by cross-inhibition type. Three sub-panels are shown, from left to right: mCherry → GFP (n = 18 replicates across n = 8 clonal cell lines), GFP → mCherry (n = 27 replicates across n = 6 clonal cell lines), mCherry ↔ GFP (bi-directional; n = 29 replicates across n = 7 clonal cell lines). Dot colors group replicates by clonal cell line; bold dot with crosshairs indicates the mean and standard deviation across replicates for each clone. Image processing in Methods, Image Analysis.

**Fig. S14.**
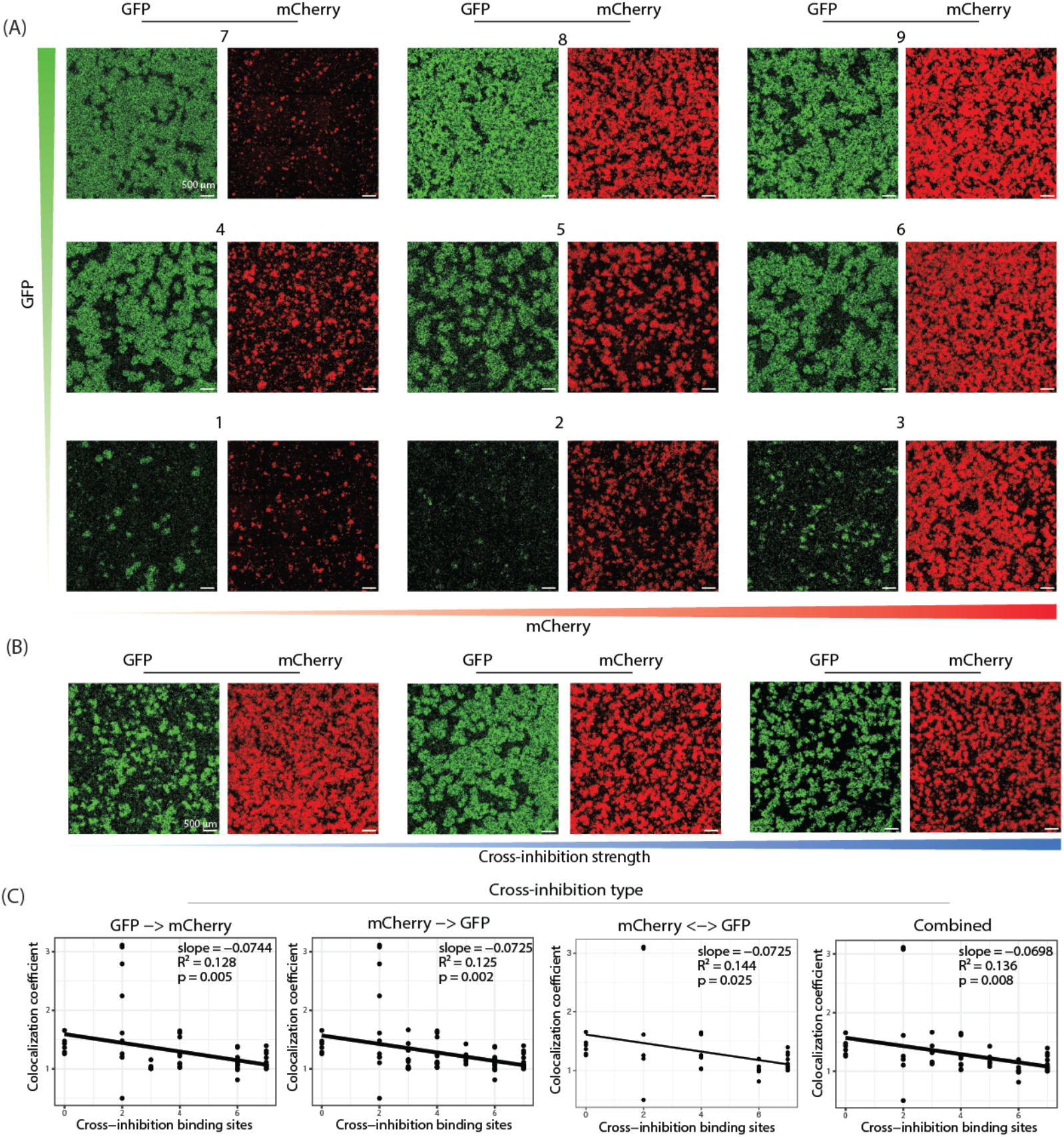
related to Fig. 5. Characterization of the resulting morphospace from tuneable cross-inhibiting dual-JAPI circuits. Single-channel fluorescence microscope endpoint images at day 4 of nine representative clonal cross-inhibiting dual-JAPI cell lines, corresponding to the merged images shown in Fig. 5H. Initial condition: homogeneous inactivated cell lawn. Scale bar = 500 µm. (**B**) Single-channel fluorescence microscope endpoint images at day 4 of three cross-inhibiting dual-JAPI clonal cell lines with significant negative cross-correlation between GFP and mCherry signals, shown left to right with increasing cross-inhibition strength (indicated by the blue gradient bar below), corresponding to the merged images shown in Fig. 5I. Initial condition: homogeneous inactivated cell lawn. Scale bar = 500 µm. (**C**) Scatter plots with linear regression of the colocalization coefficient, computed as the ratio of the measured DICE coefficient to the null DICE coefficient, as a function of the number of cross-inhibitory binding sites in the inhibitor promoter, separated by cross-inhibition type. Four sub-panels are shown, from left to right: GFP → mCherry (n = 60 observations from n = 13 genotyped clonal cell lines), mCherry → GFP (n = 51 observations from n = 20 genotyped clonal cell lines), mCherry ↔ GFP (bi-directional, n = 31 observations from n = 7 genotyped clonal cell lines), and combined across all three types (n = 76 observations from n = 25 genotyped clonal cell lines). Each dot is an individual clone; black line is the best-fit linear regression. Slope, R^2^, and p-value indicated in the figure for each sub-panel.

**Fig. S15.**
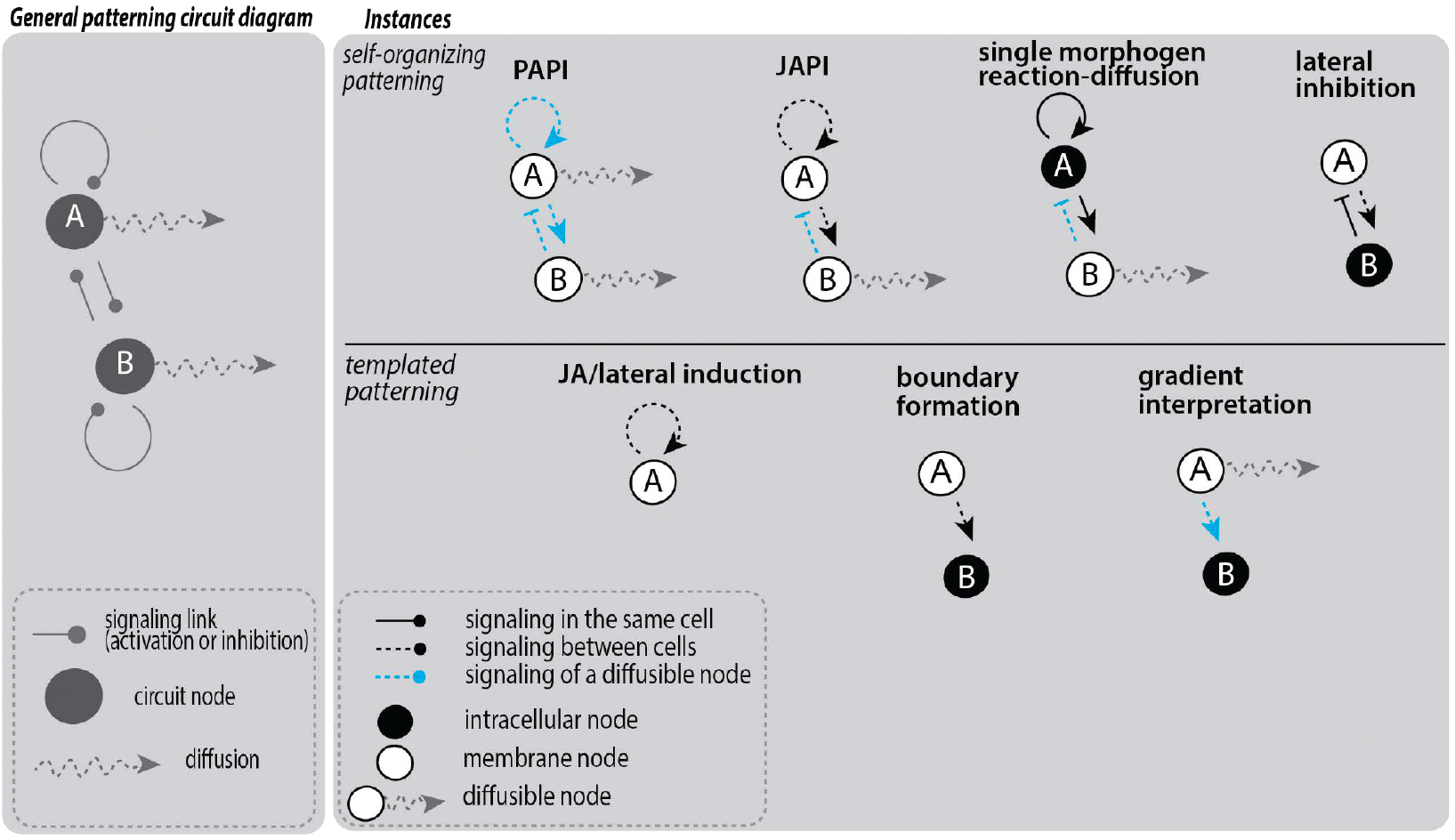
related to Discussion. A common framework for multicellular patterning circuit architectures. This scheme represents many patterning architectures as variations on a single template. As in standard graph-based representations of patterning circuits, each architecture is built from two nodes (A, B) connected by signaling links. To this we add two independent annotations: one on the nodes, specifying where each species is localized (intracellular, membrane-tethered, or secreted), and one on the links, specifying how each interaction is transmitted (within the same cell, between cells by direct contact, or by diffusion). To define a specific architecture, one specifies which species are present and where they are localized, then which interactions connect them and their type. Because every interaction requires a mode of transport, this view generalizes reaction-diffusion: classical reaction-diffusion (PAPI) is the special case in which all transport occurs by diffusion. The same structure accommodates other reaction-diffusion architectures, such as JAPI, where A is transported by juxtacrine cell-to-cell auto-activation, as well as architectures not classically associated with reaction-diffusion, such as gradient interpretation. Node fill indicates localization: filled, intracellular; open, membrane-tethered; open with a wavy arrow, secreted, where the wavy arrow denotes that the species is itself diffusible. Line style indicates the transport mode of each interaction: solid, within the same cell (cis); dashed black, between cells by direct contact (trans); dashed blue, by diffusion, denoting that a diffusible species acts on a target by diffusing to it. Arrowheads indicate activation and flat bars indicate inhibition. In the general patterning circuit diagram (left), grey nodes and links indicate unspecified localization, transport mode, and sign. On the right, instances are grouped into self-organizing patterning, in which the pattern emerges from the circuit’s own dynamics, and templated patterning, in which a pre-existing source instructs the pattern.

## Supplementary Notes

### Supplementary Note 1

#### Linear Stability Analysis of JAPI

##### Model

This note analyzes the linear stability of a juxtacrine-activator paracrine-inhibitor (JAPI) reaction-diffusion system, in which the activator is membrane-tethered and propagates by cell-cell contact, while the inhibitor is paracrine and diffuses. JAPI is contrasted with classical paracrine-activator paracrine-inhibitor (PAPI) systems, the prototypical implementation of local-activation/lateral-inhibition (LALI) reaction-diffusion patterning, in which both species diffuse. We work in units where the lattice spacing is unity, and assume periodic boundary conditions on a one-dimensional lattice (taken either infinite or sufficiently large that finite-size effects are negligible; small finite-lattice corrections are discussed where relevant).

We consider a JAPI system in which the activator is membrane-tethered and propagates through neighbor-mediated relay, while the inhibitor is diffusible. The system is formulated on a one-dimensional lattice of cells indexed by integer *j* along the lattice (so that *j* serves simultaneously as cell label and, in unit-spacing units, as spatial coordinate), with the understanding that all cells are geometrically identical and that the lattice is translationally invariant. The juxtacrine interaction kernel is written as *K*_*jl*_ = *K*(*j* − *l*), where *K*(*r*) is a symmetric, nonnegative, row-normalized function of the cell-to-cell displacement *r*; its specific form (nearest-neighbor cosine kernel) is given below. In the main text we use the shorthand *Ka* for the kernel-weighted activator input; here we work with the explicit lattice form (*Ka*)_*j*_ ≡ ∑_*l*_ *K*_*jl*_ *a*_*l*_. Under these assumptions the dynamics take the form

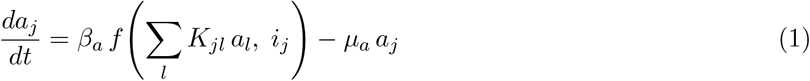

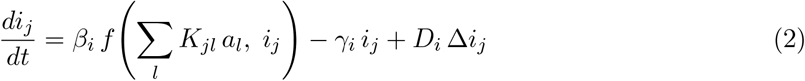

where *a*_*j*_ and *i*_*j*_ denote activator and inhibitor levels in cell *j*; *β*_*a*_, *β*_*i*_ are production rates; *µ*_*a*_, *γ*_*i*_ are degradation rates; and *D*_*i*_ is the inhibitor diffusion coefficient. The symbol Δ denotes the nearest-neighbor discrete Laplacian on the cell index,

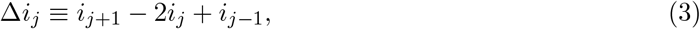

a finite-difference operator between cells (not a continuum derivative within a cell). The Fourier representation of Δ on the lattice is given in the Fourier Decomposition section below. Both species share the same nonlinear production function *f*, evaluated at the juxtacrine-weighted activator input (*Ka*)_*j*_ and the local inhibitor level *i*_*j*_. This coupling structure ensures that the Fourier transform of the kernel, 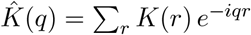 (defined precisely in the Fourier Decomposition section), appears in both rows of the linear operator derived below (Turing, 1952; Murray, 2003).

###### Production function

For the steady state analysis below we take *f* to be a two-input Hill function representing competitive inhibition,

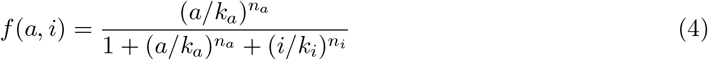

where *k*_*a*_ and *k*_*i*_ are activation and inhibition thresholds and *n*_*a*_, *n*_*i*_ are Hill coefficients. We take *f* to be shared between activator and inhibitor because this reflects the experimental implementation (see Fig. 2 in main text), in which both species are driven by a single transcriptional channel: the synNotch receptor activation directly drives expression of both the membrane-tethered activator and the secreted inhibitor through a common promoter. The form of the mode-dependent linear operator extends to the case of distinct production functions *f*_*a*_≠ *f*_*i*_ with notational changes; the compact determinant identity in the existence proof below is written for the shared-promoter case used here.

###### Scope of the linear stability analysis

The specific functional form of the production functions determines the location and number of homogeneous steady states, which we analyze for the competitive inhibition case in Supplementary Note 3. The linear stability analysis that follows is, however, agnostic to the specific form of the production functions: it requires only that the steady state (*a*_0_, *i*_0_) exists and that the partial derivatives *f*_*a*_ and *f*_*i*_ evaluated there have the appropriate signs, positive for self-activation and negative for inhibition. The results therefore apply to any reaction-diffusion system with this architecture, not only the competitive inhibition case (Gierer & Meinhardt, 1972; Kondo & Miura, 2010).

##### Steady State Analysis

Before linearizing, we establish the fixed point structure of the system. Seeking spatially homogeneous solutions *a*_*j*_ = *a, i*_*j*_ = *i*, the discrete Laplacian vanishes and the juxtacrine sum reduces to ∑_*l*_ *K*_*jl*_ *a*_*l*_ = *κ a*, where

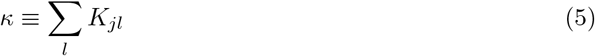

is the total juxtacrine coupling weight, independent of *j* by translational invariance. With the kernel *K*_*jl*_ = *K*(*j* − *l*) assumed symmetric (*K*(*r*) = *K*(−*r*)), nonnegative (*K*(*r*) ≥ 0), and row-normalized (∑_*r*_ *K*(*r*) = 1), the Fourier transform 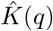 is real, even, and bounded above by 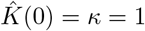. For unnormalized kernels, the local activator amplification at *q* = 0 becomes *κ β*_*a*_ *f*_*a*0_, where *f*_*a*0_ ≡ ∂*f/*∂*a* evaluated at the activated homogeneous steady state (its explicit form for the competitive Hill function is given in Eq. (15) below), and the conditions derived below should be replaced accordingly with the substitution *α* → *κα*. The system reduces to the single-cell ODE:

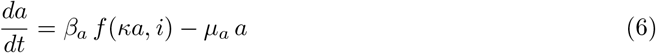

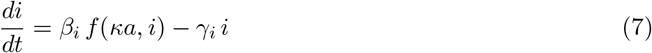

###### Trivial fixed point

The origin (*a, i*) = (0, 0) is always a fixed point. For *n*_*a*_ *>* 1, *f* (0, 0) = 0 and ∂*f/*∂*a*|_(0,0)_ = 0, so the Jacobian at the origin is diagonal with eigenvalues −*µ*_*a*_ and −*γ*_*i*_, both strictly negative. The off state is therefore linearly stable and does not support Turing instability whenever *n*_*a*_ *>* 1. (For *n*_*a*_ = 1 the Jacobian acquires a finite off-diagonal contribution and origin stability becomes parameter-dependent; we do not consider this case here.)

###### Nontrivial fixed points

Existence of a nontrivial activated fixed point (*a*_0_, *i*_0_) with *a*_0_ *>* 0 depends on both activator and inhibitor parameters in the coupled two-species system. A useful lower bound on the activator production rate required for such a fixed point follows from the reduction of the system to the activation branch alone (called JA, see also main text). A nontrivial fixed point of the JA system satisfies:

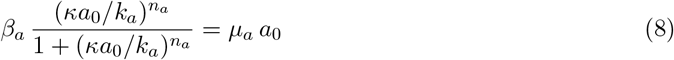

The onset of bistability in the JA reduction, the parameter value at which a pair of nontrivial fixed points appears, is found by requiring this equation to hold simultaneously with the tangency condition that the production and degradation curves have equal slopes. Solving these two conditions jointly gives the critical activator concentration:

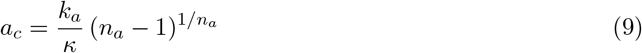

and the critical production rate:

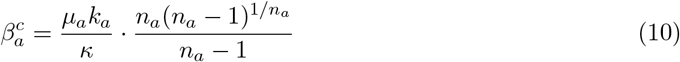

Bistability of the JA system requires *n*_*a*_ *>* 1 and 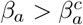. We note that the full bifurcation condition for the coupled (*a, i*) system is more complex and depends on the inhibitor parameters as well; the JA bistability condition above provides a useful lower bound. A complete analysis of the two-species fixed point structure in dimensionless parameter space, including the dependence on Hill coefficients and the full two-species phase portrait, is provided in Supplementary Note 3.

When the JA bistability condition holds, the one-dimensional JA reduction has three fixed points along the activator axis: the stable off state (0, 0), an unstable threshold state, and a stable activated state. In the full two-species system, the inhibitor tracks the activator at steady state through the ratio *i*_0_*/a*_0_ = *β*_*i*_*µ*_*a*_*/*(*β*_*a*_*γ*_*i*_), which follows from dividing the two steady-state equations.

All subsequent linear stability analysis is performed around the nontrivial activated fixed point (*a*_0_, *i*_0_), whose existence we assume in what follows.

##### Homogeneous Steady State

We now identify the activated steady state (*a*_0_, *i*_0_) to linearize around. The steady-state values satisfy:

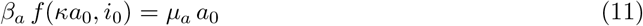

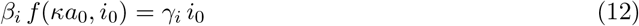

##### Partial Derivatives of the Production Function

Linearization requires the partial derivatives of *f* with respect to each argument. Writing 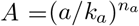 and 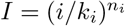, so that *f* = *A/*(1 + *A* + *I*), direct differentiation gives

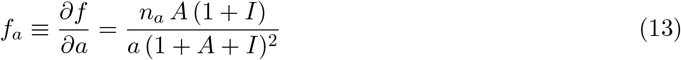

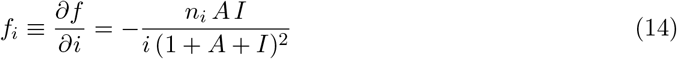

Evaluated at the homogeneous state, with 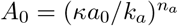 and 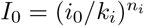,

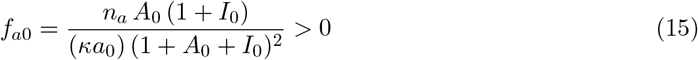

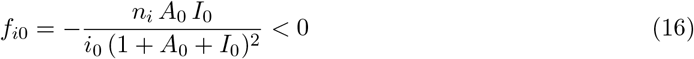

Note that *f*_*a*0_ is evaluated at the juxtacrine input *κa*_0_ rather than *a*_0_ alone, reflecting the fact that the effective activation signal experienced by each cell in the homogeneous state is amplified by the total coupling weight *κ*.

##### Linearization

We perturb around the homogeneous state, writing *a*_*j*_ = *a*_0_ + *δa*_*j*_ and *i*_*j*_ = *i*_0_ + *δi*_*j*_. The perturbed juxtacrine input is

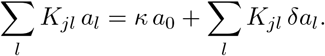

Expanding *f* to first order and canceling steady-state terms using the steady-state equations yields the linearized system

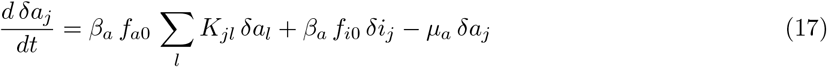

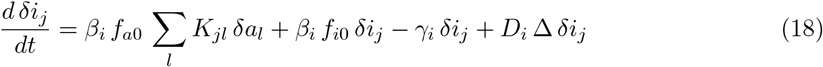

The architecture-specific structure is already apparent: the activator couples spatially through the juxtacrine kernel *K*, while the inhibitor couples through diffusion.

##### Fourier Decomposition

Cells are indexed by integers *j* along the 1D lattice, so *j* serves simultaneously as cell label and (in unit-spacing units) spatial coordinate; both the discrete Laplacian defined above and the Fourier ansatz below rely on this 1D ordered labeling. Extensions to higher-dimensional lattices replace *j* with a multi-index without changing the structure of the analysis.

We seek normal-mode solutions of the form

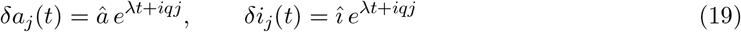

where *q* ∈ (−*π, π*] is the lattice wavenumber and *λ* is the complex growth rate. Under this ansatz the convolution with *K* diagonalizes,

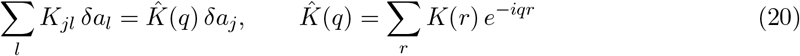

At *q* = 0 this reduces to 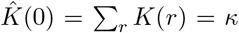, consistent with the homogeneous-state analysis above.

###### Discrete Laplacian

For nearest-neighbor diffusion on a 1D lattice the discrete Laplacian acting on a Fourier mode gives

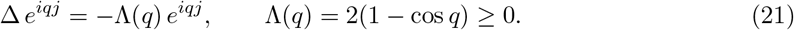

In the long-wavelength limit *q* → 0 we have Λ(*q*) = *q*^2^ + *O*(*q*^4^), recovering the continuum Laplacian eigenvalue and ensuring consistency with reaction-diffusion theory at large spatial scales.

###### Nearest-neighbor juxtacrine kernel

For a symmetric nearest-neighbor kernel 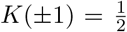, *K*(*r*) = 0 otherwise, the Fourier transform evaluates to

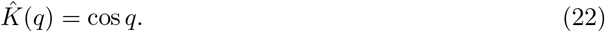

This is not a delta function (whose transform would be flat); rather it is a finite-range kernel whose Fourier transform is positive at low *q* (coherent amplification of long-wavelength modes) and becomes negative near *q* = *π* (suppression of checkerboard modes). The sign change at high wavenumber is a qualitative feature of discrete juxtacrine coupling that has no analog in diffusive activator spread, though at low wavenumbers the cosine kernel admits an effective-diffusion rewriting (analyzed in the comparison section below). The nearest-neighbor kernel is the natural minimal model for juxtacrine signaling in which cells interact only with immediate neighbors; more extended or graded kernels (e.g. a Gaussian 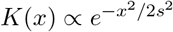) can be introduced if a continuously tunable interaction range is desired, but introduce an additional phenomenological length scale not analyzed here.

###### Eigenvalue problem

Substituting the Laplacian and kernel transforms into the linearized dynamics, the system reduces for each mode *q* to the 2 × 2 eigenvalue problem

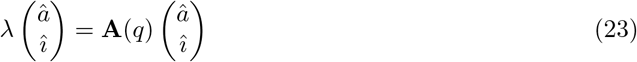

with linear operator

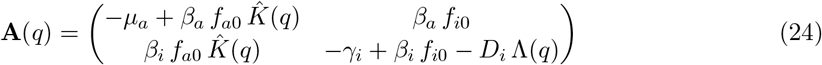

This is the central linear operator for JAPI.

##### Dispersion Relation

The eigenvalues of **A**(*q*) are found from the characteristic equation

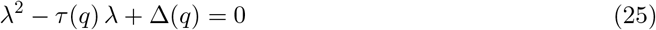

where the trace and determinant are

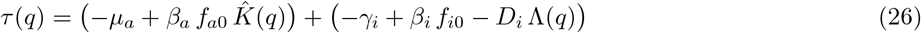

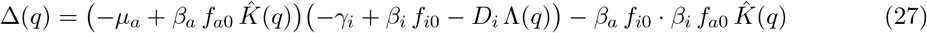

The two eigenvalues are

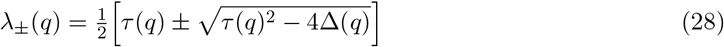

and the relevant dispersion relation is *λ*_max_(*q*) = max{Re(*λ*_+_), Re(*λ*_−_)}. A patterned state arises when *λ*_max_(*q*) *>* 0 for at least one mode *q* 0 while the *q* = 0 mode remains stable, and the dominant pattern wavelength at onset is set by the mode *q*^∗^ that maximizes *λ*_max_(*q*).

##### Stability Conditions

###### Homogeneous stability

The *q* = 0 mode corresponds to spatially uniform perturbations. Since Λ(0) = 0 and 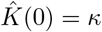, stability requires

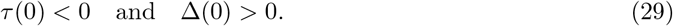

These are the standard Routh–Hurwitz conditions for the local reaction system, ensuring that the homogeneous steady state is stable to uniform perturbations.

##### Spatial instability

A patterned state emerges when the homogeneous conditions hold yet *λ*_max_(*q*) *>* 0 for some *q*≠ 0. Under the trace-negative conditions verified in the next subsection, this is equivalent to Δ(*q*) becoming negative at some nonzero wavenumber, here controlled by the interplay between 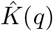 and Λ(*q*) rather than by two diffusion coefficients. The existence of parameter regimes where this occurs is proven in the next subsection.

##### Existence of Turing Instabilities in JAPI

We show that the JAPI operator **A**(*q*) derived above admits finite-wavenumber instabilities of the homogeneous on state under the standard local activator–inhibitor prerequisites, establishing that the Turing regime is accessible to the JAPI architecture. The closed-form threshold, mode selection, and unstable band are the object of future theoretical work; here we establish existence.

###### Setup and prerequisites

Introduce the abbreviations

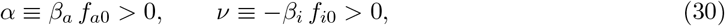

where positivity follows from *f*_*a*0_ *>* 0 and *f*_*i*0_ *<* 0 at the activated steady state. *Sign convention*. The minus sign in the definition of *ν* is chosen so that *ν* is positive; with this convention *β*_*i*_*f*_*i*0_ = −*ν*, so the (2,2) entry of **A**(*q*) reads −*γ*_*i*_ − *ν* − *D*_*i*_Λ(*q*). All expressions below (in particular the determinant identity in Eq. (32)) follow this convention. An alternative convention in which *ν* is defined as *β*_*i*_*f*_*i*0_ (negative-valued) would give equivalent results with sign flips wherever *ν* appears; e.g., the leading term of Eq. (32) would read *µ*_*a*_(*γ*_*i*_ − *ν*) rather than *µ*_*a*_(*γ*_*i*_ + *ν*), with the same numerical value because of the sign flip in *ν*. The local prerequisites for a Turing-type instability of the on state are:

i. *α > µ*_*a*_ (local activator self-amplification at *q* = 0)
ii. *α* − *µ*_*a*_ − *γ*_*i*_ − *ν <* 0 (Routh–Hurwitz trace condition at *q* = 0)
iii. *µ*_*a*_*ν > γ*_*i*_(*α* − *µ*_*a*_) (Routh–Hurwitz determinant condition at *q* = 0) Conditions (ii) and (iii) together ensure homogeneous stability of the on state.

We additionally assume that the lattice kernel and Laplacian eigenvalue satisfy the normalization properties

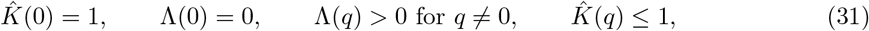

and that the lattice spectrum contains at least one allowed nonzero mode in a sufficiently small neighborhood of *q* = 0 where the continuity arguments below apply. This is automatic for an infinite lattice or for a sufficiently large finite lattice, but should be checked mode-by-mode on small finite lattices. For the cosine kernel 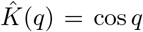, this holds in the continuous-*q* or sufficiently large finite-lattice limit whenever *α > µ*_*a*_; on small finite lattices the allowed modes must be checked explicitly.

###### Determinant structure

Direct expansion of det **A**(*q*) using Eq. (24), after using *β*_*a*_ *f*_*i*0_ · *β*_*i*_ *f*_*a*0_ = −*αν*, yields

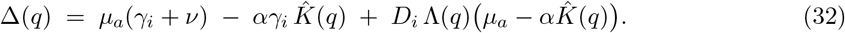

The right-hand side separates into a *q*-dependent local part, 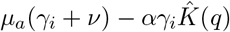, and a spatial-coupling part 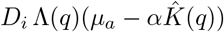 that is linear in *D*_*i*_. The coefficient of *D*_*i*_ is determined by the sign of 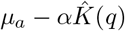: for modes where 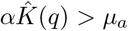, the coefficient is negative, and Δ(*q*) decreases linearly as *D*_*i*_ increases.

###### All modes are stable at *D*_*i*_ = 0

Before showing that finite *D*_*i*_ can destabilize a nonzero mode, we verify that no spatial instability exists in the absence of inhibitor diffusion. At *D*_*i*_ = 0,

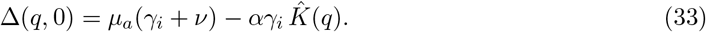

Using 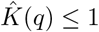,

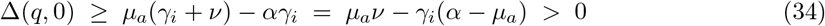

by condition (iii). The trace is

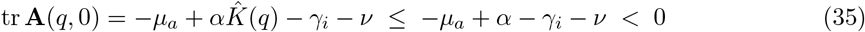

by condition (ii). All modes are therefore linearly stable at *D*_*i*_ = 0, regardless of wavenumber. Spatial instability in JAPI is consequently *diffusion-driven*: it requires *D*_*i*_ *>* 0 and emerges as *D*_*i*_ increases beyond a finite threshold, in the standard Turing sense.

###### Existence of a destabilizing finite-wavenumber mode

By continuity of 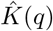 and condition (i), there exists *q*^∗^≠ 0 sufficiently close to *q* = 0 such that both

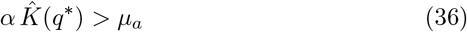

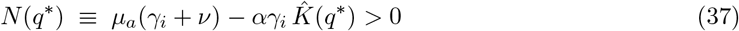

hold. The first follows from 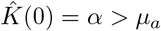, and the second follows from *N* (0) = *µ*_*a*_*ν* −*γ*_*i*_(*α*−*µ*_*a*_) *>* (condition iii), both by continuity.

For such a *q*^∗^, write

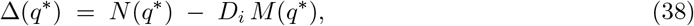

where 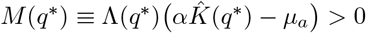. The determinant Δ(*q*^∗^) is therefore a strictly decreasing linear function of *D*_*i*_ with positive intercept *N* (*q*^∗^) and positive slope magnitude *M* (*q*^∗^). It vanishes at the finite positive threshold

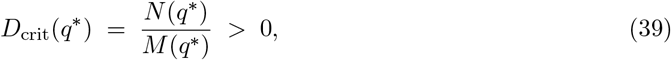

and becomes negative for all *D*_*i*_ *> D*_crit_(*q*^∗^). At this mode, the JAPI operator therefore acquires a positive-real-part eigenvalue, while the homogeneous mode remains stable because *q* = 0 is unaffected by *D*_*i*_.

###### Existence statement

Under conditions (i)–(iii) and the lattice spectral assumptions stated above, the JAPI operator **A**(*q*) admits at least one wavenumber *q*^∗^≠ 0 at which the homogeneous on state is destabilized by sufficiently strong inhibitor diffusion, while the homogeneous mode remains stable. This is the defining signature of a Turing-type instability.

Because the off state (0, 0) remains linearly stable for *n*_*a*_ *>* 1 (see Trivial fixed point analysis above), this argument establishes linear instability of the activated homogeneous branch only, not global convergence of the nonlinear system to a patterned state. Nonlinear pattern selection, saturation, and the possibility of basins of attraction containing the off state require numerical simulation or weakly nonlinear analysis, and are not the object of this note.

**The JAPI architecture therefore admits Turing instabilities under the same type of local activator–inhibitor prerequisites familiar from classical PAPI**, a comparison developed in detail in the next subsection, with the destabilizing spatial threshold provided by *D*_*i*_ alone rather than by a ratio of two diffusion coefficients. The precise threshold

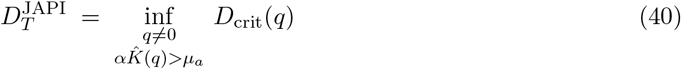

(replaced by a minimum over allowed lattice modes *q*_*m*_ = 2*πm/N* on a finite periodic lattice of *N* cells) depends on the lattice geometry through 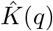 and Λ(*q*), and on the lattice spectrum on finite lattices. For the 1D nearest-neighbor cosine kernel in the continuous-*q* limit, the infimum is attained in the interior of the allowed band under the stated conditions and admits a direct closed-form expression by minimization of *D*_crit_(*q*) over *q*. We do not pursue this in detail here; closed-form analysis of 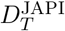 for general kernels, finite-lattice corrections, higher-dimensional geometries, and the relationship to the classical PAPI threshold, are the object of future theoretical work.

###### Worked example

The following parameter set is presented as a *local-Jacobian* demonstration of existence: it satisfies conditions (i)–(iii) and the kernel assumptions, and so admits a Turing instability of the operator **A**(*q*). We do not claim it corresponds to a particular set of Hill-function parameters (*β*_*a*_, *β*_*i*_, *k*_*a*_, *k*_*i*_, *n*_*a*_, *n*_*i*_) satisfying the homogeneous steady-state equations; for the competitive Hill production function, the steady-state constraint imposes *α/µ*_*a*_ *< n*_*a*_ and *ν/γ*_*i*_ *< n*_*i*_, so realizing the example below from the Hill model would require *n*_*a*_ *>* 5 and *n*_*i*_ *>* 10, which are higher than the experimentally measured Hill coefficients in the implementation analyzed in the main text. A Hill-realizable parameter set in the irregular regime relevant to the experimental implementation is analyzed in future theoretical work; the example here serves only to exhibit a Jacobian-level parameter set in which the existence proof applies.

Take

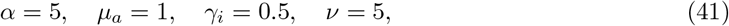

with the 1D nearest-neighbor kernel 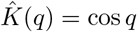. Verification of (i)–(iii):

i. *α* − *µ*_*a*_ = 4 *>* 0 ✓
ii. *α* − *µ*_*a*_ − *γ*_*i*_ − *ν* = −1.5 *<* 0 ✓
iii. *µ*_*a*_*ν* − *γ*_*i*_(*α* − *µ*_*a*_) = 3 *>* 0 ✓

Pick *q*^∗^ with cos *q*^∗^ = 0.8. Then 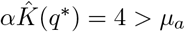, Λ(*q*^∗^) = 0.4, and

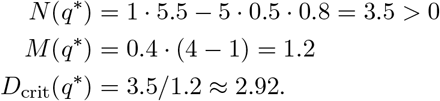

For any *D*_*i*_ *>* 2.92, the mode *q*^∗^ satisfies Δ(*q*^∗^) *<* 0. At *q* = 0, both Routh–Hurwitz conditions hold regardless of *D*_*i*_: *τ* (0) = − 1.5 *<* 0 and Δ(0) = 3 *>* 0, so the homogeneous mode remains stable. The system is therefore in a Turing regime for *D*_*i*_ *>* 2.92 at this parameter set. The chosen *q*^∗^ (with cos *q*^∗^ = 0.8) is illustrative; direct numerical minimization of *D*_crit_(*q*) over the allowed band gives the true onset *D*_*T*_ ≈ 2.46 for this parameter set, at the optimal mode cos *q*_*T*_ ≈ 0.65, *q*_*T*_ ≈ 0.86 rad (wavelength ≈ 7.3 cells).

###### Comment on the geometric threshold

For the 1D nearest-neighbor cosine kernel, the expression

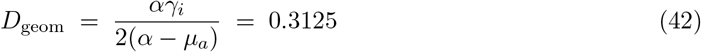

for the parameters above, obtained by requiring that the determinant minimum lie within the physical interval of accessible wavenumbers, has appeared in informal analyses as a candidate threshold. *D*_geom_ is strictly necessary for finite-*q* instability (it gives the lower bound on *D*_*i*_ for the minimum of Δ(*c*) over *c* = cos *q* to fall in the physical band) but is *not* the threshold itself: 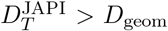 in general. A direct numerical minimization of *D*_crit_(*q*) gives *D*_*T*_ ≈ 2.46 for the parameter set above. A closed-form expression for 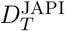, including for general kernels, is developed in future theoretical work.

##### Structure of the Linear Operator: Comparison with Classical LALI

We now compare the JAPI linear operator derived above with the corresponding operator for a classical paracrine activator–paracrine inhibitor (PAPI) system. The comparison clarifies which features are shared between the two architectures and which are specific to JAPI.

For PAPI, an identical linearization procedure yields

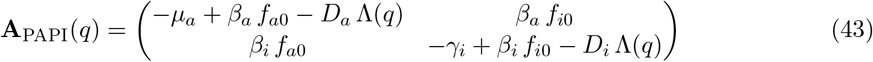

where *D*_*a*_ is the activator diffusion coefficient and Λ(*q*) is the same discrete Laplacian eigenvalue as in the JAPI case. A note on notation: *f*_*a*0_ in the PAPI operator is evaluated at the homogeneous state *a*_0_ (no juxtacrine input), and so is formally distinct from *f*_*a*0_ in JAPI which is evaluated at the juxtacrine input *κa*_0_. For the symmetric nearest-neighbor kernel *K*(±1) = 1*/*2 used here, *κ* = 1 and the two evaluations coincide; the structural comparison below holds in the general case as well.

Placing the two operators side by side:

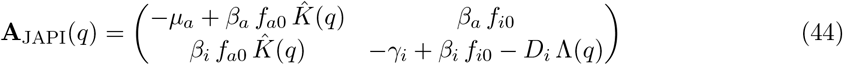

The two operators are identical except in the entries involving the activator’s spatial coupling. Specifically:

- **Entry (1**,**1)**. In PAPI, the activator self-term carries −*D*_*a*_Λ(*q*), a diffusive penalty whose *q*-dependent shape Λ(*q*) is fixed by the lattice geometry and whose amplitude *D*_*a*_ is an independent biophysical parameter, tunable without affecting the local reaction kinetics. In JAPI, this is replaced by 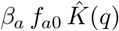, where the *q*-dependent shape 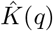 is again fixed by the lattice geometry but the amplitude is set by *β*_*a*_ *f*_*a*0_, the same coupling that determines the local self-amplification at *q* = 0. The kernel shapes are fixed by lattice geometry in both architectures; what differs is whether the amplitude is independent of the reaction kinetics (*D*_*a*_ in PAPI, yes) or entangled with them (*β*_*a*_ *f*_*a*0_ in JAPI, no).
- **Entry (2**,**1)**. In PAPI, the inhibitor cross-term *β*_*i*_ *f*_*a*0_ carries no 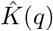 factor because inhibitor production responds to the local activator perturbation rather than to a juxtacrine-weighted activator input. In JAPI, this entry carries 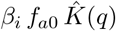 because the inhibitor is produced in response to the juxtacrine activator input, which is itself mode-dependent. The kernel therefore modulates not only the activator self-term but also the inhibitor cross-coupling.
- **Entries (2**,**2) and (1**,**2)**. Identical in both systems. The inhibitor diffusion term −*D*_*i*_Λ(*q*) and the inhibitory feedback *β*_*a*_ *f*_*i*0_ are unchanged.

The key consequence is that *D*_*a*_ drops out of the JAPI patterning problem entirely (Marcon et al., 2016). The instability mechanism is qualitatively preserved: both architectures combine local activator self-amplification with inhibitor-mediated long-range suppression. However, JAPI implements the activator-side spatial structure through the juxtacrine kernel rather than through an independently tunable activator diffusion coefficient.

For convenience in the discussion that follows, we denote the diagonal entries of the JAPI operator as 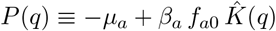 and *Q*(*q*) ≡ −*γ*_*i*_ + *β*_*i*_ *f*_*i*0_ − *D*_*i*_ Λ(*q*) (activator and inhibitor self-couplings, respectively). The inhibitor self-coupling *Q*(*q*) is identical in both architectures and always negative: *f*_*i*0_ *<* 0 ensures the inhibitor self-loop is stabilizing, and −*D*_*i*_Λ(*q*) makes *Q*(*q*) more negative at higher *q*. The instability therefore requires *P* (*q*) *>* 0 at some nonzero wavenumber, meaning 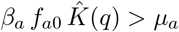 at those modes.

What differs between the two systems is the parametric structure of *P* (*q*). In PAPI, *P*_PAPI_(*q*) = −*µ*_*a*_ + *β*_*a*_ *f*_*a*0_ − *D*_*a*_Λ(*q*), and the amplitude on the spatial term is *D*_*a*_, an independent molecular parameter that can be tuned without affecting the local reaction kinetics: decreasing *D*_*a*_ keeps *P*_PAPI_(*q*) positive over a wider range of wavenumbers and widens the unstable band. In JAPI, 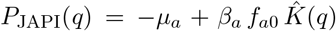, and the amplitude on the spatial term is *β*_*a*_ *f*_*a*0_, which is not independent of the reaction kinetics: the same coupling sets both the *q* = 0 self-amplification (via *β*_*a*_ *f*_*a*0_ directly) and the *q*-dependent activator coupling (via 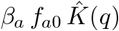). This entanglement is the precise sense in which JAPI is a more constrained system: the qualitative instability mechanism is the same, but the independent tuning of activator spatial spread through a parameter decoupled from the reaction kinetics, which *D*_*a*_ provides in PAPI, has been removed by construction.

For the nearest-neighbor kernel, 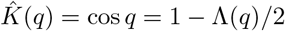. Substituting into the JAPI activator self-coupling gives

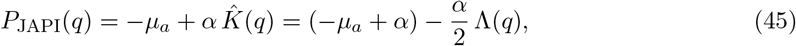

which has exactly the same *q*-dependence as a PAPI activator self-coupling with an effective activator diffusion coefficient 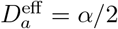. In this sense, the JAPI (1,1) entry is diffusion-like in the nearest-neighbor case, but with the effective diffusion amplitude slaved to the local activator gain *α* = *β*_*a*_ *f*_*a*0_ rather than tunable independently. The genuinely architecture-specific structural distinction is therefore not the form of the (1,1) entry, but the kernel filtering of the (2,1) off-diagonal entry: in PAPI the activator-to-inhibitor coupling carries no *q*-dependence, while in JAPI it carries a 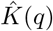 factor inherited from the juxtacrine activator input. For more extended kernels, the (1,1) entry can also deviate from the standard diffusive *q*^2^ form at intermediate and high wavenumbers; in the 1D nearest-neighbor case analyzed here, the deviation is small at low *q* but the sign change of cos *q* near *q* = *π* remains a qualitative difference from Fickian diffusion. Closed-form analysis of selected wavenumber, instability bandwidth, and parameter space geometry that follow from these distinctions, including the consequences of the (2,1) entry’s kernel filtering, are the object of future theoretical work.

##### Summary

Linear stability analysis of JAPI yields a 2 × 2 mode-dependent growth operator **A**(*q*) in which juxtacrine relay enters through the Fourier transform of the neighbor kernel 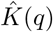, while inhibitor diffusion contributes the stabilizing penalty −*D*_*i*_Λ(*q*). Under the standard local activator–inhibitor prerequisites for Turing patterning, we prove that the JAPI operator admits finite-wavenumber instabilities, establishing that the Turing regime is accessible to the JAPI architecture. The result is a sufficient condition stated at the Jacobian level: it establishes that the JAPI architecture admits Turing-type instability whenever the local activator–inhibitor prerequisites are met at some activated steady state, independent of whether a given Hill parameterization realizes those prerequisites. Direct comparison with the classical PAPI operator shows that the two systems differ in exactly two entries, both involving the activator spatial coupling, while sharing identical inhibitor diffusion, inhibitory feedback, and local reaction structure.

The instability mechanism is qualitatively preserved in JAPI: Turing-type patterning requires local activator self-amplification (*P* (*q*) *>* 0 at some nonzero wavenumber) combined with long-range inhibitory suppression (*Q*(*q*) *<* 0 driven by *D*_*i*_Λ(*q*)). What JAPI loses relative to PAPI is *D*_*a*_ as an independent tuning parameter for the width of the unstable band. The activator spatial term is instead fixed by kernel geometry and circuit gain *β*_*a*_ *f*_*a*0_, which are entangled with the rest of the reaction network. The closed-form threshold expression for the JAPI Turing onset, its precise relationship to the classical PAPI threshold, and the structure of the unstable band are the object of future theoretical work.

## Supplementary Note 2

### Physical Interpretation of the Juxtacrine Activator Velocity Scale

Supplementary Note 1 establishes that the activator diffusion coefficient *D*_*a*_ does not appear in the JAPI patterning problem, and proves the existence of Turing-type finite-wavenumber instabilities under the standard local prerequisites. The Note 1 analysis identifies the conditions under which spatially patterned states can arise. The present note addresses a complementary question: how rapidly an activated domain expands through the tissue before inhibitor-mediated arrest. To this end, we develop the physical interpretation of what governs the activator’s spatial propagation in JAPI: an effective propagation velocity *v*_*a*_ characterizing the cell-to-cell relay of the membrane-tethered activator. We derive a minimal threshold-crossing estimate for *v*_*a*_ in an inhibitor-suppressed limit; a full traveling-wave analysis incorporating the coupled neighbor dynamics, lattice-discreteness effects, and inhibitor-mediated corrections is the subject of future theoretical work.

#### Setup

For reference, the full PAPI and JAPI dynamics take the forms:

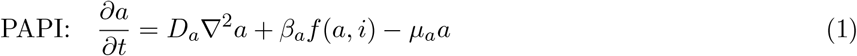

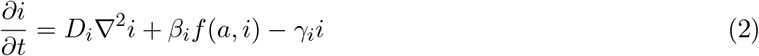

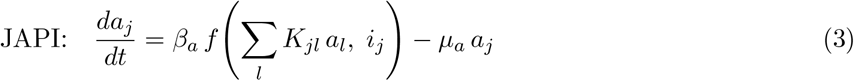

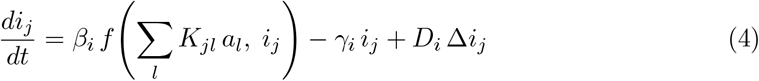

The PAPI activator equation contains the diffusion term *D*_*a*_∇^2^*a*; the JAPI activator equation replaces it with the discrete kernel sum over neighboring cells. The PAPI equations are written in continuum, appropriate for the diffusive regime; the JAPI equations are written on a lattice, reflecting the cell-by-cell architecture of the relay (Murray, 2003). The juxtacrine kernel *K*_*jl*_ is taken to be nonnegative, symmetric, row-normalized, and self-input free (*K*_*jj*_ = 0), as stated in Supplementary Note 1.

The full coupled four-equation system is analyzed in Supplementary Note 1 (linearization and stability) and Supplementary Note 4 (arrest dynamics). For the velocity-scale analysis that follows, we work in an inhibitor-suppressed limit appropriate to the early expansion phase of an activated domain: before significant inhibitor accumulation, the inhibitor argument of *f* contributes negligibly to activator production. Operationally, we set *f* (·, *i*_*j*_) ≈ *f* (·, 0), so the activator equation reduces to a one-species front-propagation problem driven by activator self-amplification and degradation alone. This limit also describes exactly the transceiver configuration where the inhibitor branch is genetically absent. The circuit parameters governing *v*_*a*_ in this limit are derived and discussed below; the inhibitor parameters re-enter the dynamics when the system transitions from expansion to arrest, which is the subject of Supplementary Note 4.

#### Physical picture of juxtacrine relay

In PAPI, the activator spreads through a tissue by diffusion. Its reach at any position is characterized by the diffusion length 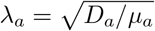, which captures the steady-state balance between diffusion and degradation in the linearized reaction-diffusion equation.

In JAPI, the membrane-tethered activator does not diffuse. Instead, an activated cell presents activator ligand to its immediate neighbors through cell-cell contact. When a neighbor receives a juxtacrine signal, it begins producing its own activator ligand, which it presents to its own neighbors. The sensitivity of a cell to its kernel-weighted input *s*_*j*_ = ∑_*l*_ *K*_*jl*_ *a*_*l*_ is set by *k*_*a*_, the input concentration at which production is half-maximal. The activator “spreads” through this relay, advancing one cell at a time as activator accumulates and drives production in successive neighbors.

The rate of this relay defines an effective propagation velocity *v*_*a*_ for the activation front. Rather than a length scale set by diffusion, JAPI has a velocity set by relay kinetics. Note that, unlike paracrine diffusion, the relay in JAPI is intrinsically discrete: the cell is the unit of advance, and the front advances in integer cell-diameter steps rather than as a continuous spatial process. This discrete relay mode is a lattice analog of front propagation in reaction-diffusion systems (Kolmogorov et al., 1937). The precise classification (Fisher-KPP, Zeldovich, or other type), and how it differs from the corresponding classical PAPI fronts given JAPI’s lattice discreteness and the absence of an independent activator diffusion coefficient, is the subject of future theoretical work.

#### Threshold-crossing estimate

To make *v*_*a*_ concrete, we consider a single relay step: a previously inactive cell *j* exposed to juxtacrine signal from already-activated neighbors. Writing the kernel-weighted incoming signal to cell *j* as *s*_*j*_(*t*) = ∑_*l*_ *K*_*jl*_ *a*_*l*_(*t*), the internal activator *a*_*j*_(*t*) evolves (in the inhibitor-suppressed limit) as

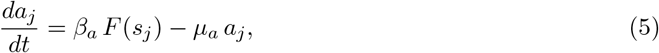

where 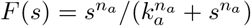 is the activation-side Hill function and *k*_*a*_ is the half-maximal input concentration of the receiver.

Two distinct thresholds appear naturally in a relay step, because the input to a downstream cell is kernel-weighted rather than equal to the activator level of the upstream cell. Cell *j* + 1 activates when its input *s*_*j*+1_ = ∑_*l*_ *K*_*j*+1,*l*_ *a*_*l*_ crosses *k*_*a*_. If the only contribution to *s*_*j*+1_ comes from cell *j* (worst case, no contribution from the other side), then cell *j* must accumulate an activator level *a*_*j*_ = *k*_*a*_*/K*_*j*+1,*j*_ before *s*_*j*+1_ reaches *k*_*a*_; this elevated level is the *output threshold θ*_*a*_. More generally, *θ*_*a*_ ∼ *k*_*a*_*/*∑ _*l*_ *K*_*lj*_, with the precise factor depending on lattice geometry and front orientation. For the symmetric nearest-neighbor coupling used here (*K*(±1) = 1*/*2), the two thresholds differ by an *O*(1) geometric prefactor, and we identify *a*_*j*_’s threshold-crossing condition with *a*_*j*_ = *k*_*a*_ for the one-step estimate that follows. (For identical cells with a delta-function kernel *K*_*j*+1,*j*_ = 1 the two thresholds coincide; the distinction arises whenever the kernel distributes the activator signal across multiple neighbors.) A self-consistent treatment distinguishing input and output thresholds enters the full traveling-wave analysis and is deferred to future theoretical work.

##### Quasi-static approximation

We assume that during the threshold-crossing event of cell *j*, the input *s*_*j*_(*t*) is approximately constant, with the activated neighbors at their quasi-steady-state activator levels on the timescale of *j*’s activation. Concretely, 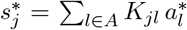, where *A* is the set of already-active neighbors and 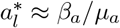 in the well-saturated limit 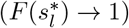. This is a leading-order approximation; a self-consistent treatment of the leading-edge cells, where upstream activator levels and their inputs evolve together, is part of the full traveling-wave analysis deferred to future theoretical work.

Under this approximation, define 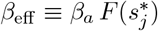, where 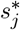 is the (constant) input to cell *j*. Starting from *a*_*j*_(0) = 0, the cell ODE has the closed-form solution

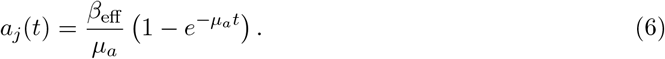

The threshold-crossing time *t*_th_ at which *a*_*j*_(*t*_th_) = *k*_*a*_ is

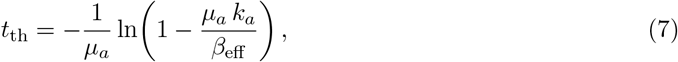

which is finite and positive provided *β*_eff_ *> µ*_*a*_ *k*_*a*_ (else the activated steady state of cell *j, β*_eff_ */µ*_*a*_, lies below *k*_*a*_ and the relay fails altogether). Including the additional intracellular latency *τ*_cell_ from receptor activation to surface ligand presentation, the total time for one relay step is *T*_step_ = *τ*_cell_ +*t*_th_, and the resulting relay velocity scale is

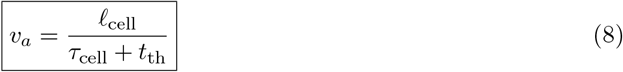

in physical length units (with *ℓ*_cell_ the cell-to-cell distance), or *v*_*a*_ = 1*/*(*τ*_cell_ + *t*_th_) in cell-per-time units.

##### Limit behaviors

Three regimes follow directly from the form of *t*_th_:

- *Well above threshold* (*β*_eff_ ≫ *µ*_*a*_ *k*_*a*_): Taylor expansion of the logarithm gives *t*_th_ ≈ *k*_*a*_*/β*_eff_, so *v*_*a*_ ≈ *ℓ*_cell_*/*(*τ*_cell_ + *k*_*a*_*/β*_eff_ ). The relay rate is dominated by production rate *β*_eff_ and sensitivity *k*_*a*_; degradation enters only as a higher-order correction.
- *Near threshold* (*β*_eff_ → *µ*_*a*_ *k*_*a*_ from above): the argument of the logarithm approaches zero, *t*_th_ → ∞, and *v*_*a*_ → 0. The relay slows and eventually fails.
- *Subthreshold* (*β*_eff_ *< µ*_*a*_ *k*_*a*_): the activated steady state *β*_eff_ */µ*_*a*_ lies below *k*_*a*_, and the relay cannot proceed at all.

This estimate captures the regime-dependent role of each circuit parameter, discussed in the next subsection.

#### Circuit parameters governing *v*_*a*_

For fixed lattice geometry, contact topology, kernel weights, and front orientation, the threshold-crossing time *t*_th_, and hence the relay velocity scale *v*_*a*_, depend on five circuit parameters, all properties of the activation circuit rather than of the medium in which signaling occurs. The sensitivities below are computed at fixed 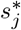 in the full coupled dynamics, the same parameters also enter through the upstream activated-cell levels 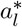 and hence through 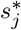 itself.

- **Activator production rate (***β*_*a*_): sets the effective production amplitude 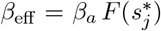. Higher *β*_*a*_ raises *β*_eff_, reduces *t*_th_, and accelerates the relay. Well above threshold, *t*_th_ ≈ *k*_*a*_*/β*_eff_ ∝ 1*/β*_*a*_, so *v*_*a*_ grows monotonically with *β*_*a*_ and saturates at *v*_*a*_ → *ℓ*_cell_*/τ*_cell_ as *β*_*a*_ → ∞.
- **Cooperativity of activation (***n*_*a*_): sets the sharpness of the Hill response *F* (*s*) around *k*_*a*_, and shapes cell behavior on both sides of *k*_*a*_, not only above it. At fixed input 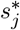 (the quasi-static regime adopted here), only the value 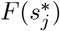 enters 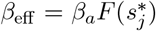, not the slope *∂F/∂s*, and the effect of *n*_*a*_ on the relay velocity depends on whether 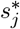 sits above or below *k*_*a*_:

– 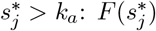 rises toward 1 faster as *n*_*a*_ increases, raising *β*_eff_ and accelerating the relay;

– 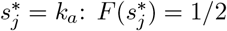 for all *n*_*a*_, so cooperativity has no effect at the half-maximal input;

– 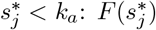 falls toward 0 faster as *n*_*a*_ increases, lowering *β*_eff_ and slowing or preventing the relay.

The local slope *∂F/∂s* enters in a full dynamic front problem where *s*_*j*_(*t*) evolves during the threshold-crossing event, but does not contribute directly in the constant-input estimate.

- **Activation sensitivity (***k*_*a*_): the half-maximal input level of the Hill function (with kernel-geometry prefactors absorbed, as discussed in the Threshold-crossing estimate above). Higher *k*_*a*_ means lower sensitivity to a given juxtacrine input level, so cell *j* requires longer to accumulate enough activator to drive production in its next neighbor. This appears in the relay-time formula as *t*_th_ = −(1*/µ*_*a*_) ln(1 − *µ*_*a*_ *k*_*a*_*/β*_eff_ ), which increases monotonically with *k*_*a*_. *k*_*a*_ also sets the propagation-failure boundary: when *µ*_*a*_ *k*_*a*_ exceeds *β*_eff_, the activated steady state *β*_eff_ */µ*_*a*_ falls below *k*_*a*_ and the relay cannot proceed.
- **Activator degradation rate (***µ*_*a*_): enters in two distinct ways. In the well-above-threshold regime, *t*_th_ ≈ *k*_*a*_*/β*_eff_ and *µ*_*a*_ contributes only as a higher-order correction; the relay rate is essentially independent of degradation. Near threshold, higher *µ*_*a*_ lowers the activated steady state *β*_eff_ */µ*_*a*_ toward *k*_*a*_, increases *t*_th_ (through the logarithm), and slows the relay. Sufficiently high *µ*_*a*_ drives the activated steady state below *k*_*a*_ (equivalently, *β*_eff_ *< µ*_*a*_ *k*_*a*_), and the relay can no longer proceed. The role of *µ*_*a*_ is therefore best characterized as setting the regime in which the relay operates rather than as a leading-order multiplier of velocity.
- **Cell response time (***τ*_cell_): the latency from receiving juxtacrine signal to presenting sufficient surface ligand to a neighbor. This summarizes the intracellular kinetics of the relay step — receptor activation, downstream signaling, transcription, translation, and membrane trafficking of the new ligand. *τ*_cell_ enters the relay step time additively: *T*_step_ = *τ*_cell_ + *t*_th_. The effect of *τ*_cell_ on *v*_*a*_ therefore depends on the regime: when *τ*_cell_ ≫ *t*_th_, the relay rate is set primarily by intracellular latency; when *t*_th_ ≫ *τ*_cell_, by the threshold-crossing dynamics.

These five circuit parameters are genetically tunable, in principle: production rate through promoter strength and translation efficiency, cooperativity through receptor architecture, sensitivity through receptor affinity, degradation rate through degron tags or fusion partners, and cell response time through signaling pathway design. The realized tunability in any particular experimental system depends on cell type, trafficking, and other context-dependent factors. Stochastic integration and multiplicity of infection of lentiviral vectors primarily shape variation in production rate; the other parameters would need to be modified through changes in the protein sequences themselves.

#### Relationship to *D*_*a*_ and to circuit control

In PAPI, the activator’s spatial behavior depends on both the molecular diffusion coefficient *D*_*a*_ (set by the molecular identity of the activator) and on circuit-encoded kinetic parameters (such as *µ*_*a*_, which contributes to the diffusion length 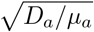. *D*_*a*_ is a biophysical property of the activator molecule and is not accessible to genetic manipulation without changing the molecule itself — the central protein-engineering challenge for synthetic PAPI in mammalian cells.

In JAPI, *D*_*a*_ is absent by construction: the membrane-tethered activator does not diffuse. For fixed tissue geometry and fixed juxtacrine contact kernel *K*_*jl*_, the kinetic parameters governing the activator’s spatial behavior are all circuit-encoded (*β*_*a*_, *n*_*a*_, *µ*_*a*_, *k*_*a*_, *τ*_cell_). This is a reduction in the total number of parameters available to control activator dynamics — JAPI offers fewer handles than PAPI — but the parameter that has been eliminated is the one that has historically been hardest to engineer. The remaining handles, in JAPI, are all genetically tunable rather than tied to molecular biophysics.

This architectural feature is what makes JAPI experimentally accessible in mammalian cells without requiring molecular engineering of differential diffusion coefficients.

#### Dynamic regime and scope

The propagation velocity *v*_*a*_ as derived above characterizes the relay rate in the inhibitor-suppressed limit *f* (·, *i*_*j*_) ≈ *f* (·, 0), corresponding either to the early expansion phase of an activated domain before significant inhibitor accumulation, or to the transceiver configuration where the inhibitor branch is absent. As inhibitor accumulates around an expanding domain, *f* (·, *i*_*j*_) is suppressed at the front by the local inhibitor level, the effective production *β*_eff_ drops, *t*_th_ grows, and the front eventually arrests. The arrest condition therefore depends critically on the inhibitor branch: on *β*_*i*_, *k*_*i*_, *n*_*i*_, *γ*_*i*_, and *D*_*i*_, which together determine the spatial profile of inhibitor around an activated domain and the threshold at which *β*_eff_ falls below *µ*_*a*_ *k*_*a*_ and relay can no longer advance. The full arrest dynamics, including the determination of arrested domain size from the inhibitor profile, are addressed in Supplementary Note 4.

A full traveling-wave analysis yielding *v*_*a*_ in closed form would relax the quasi-static approximation for *s*_*j*_(*t*), treat the coupled dynamics of activator levels across the lattice self-consistently, incorporate lattice-discreteness effects (such as front pinning, anisotropy, and propagation failure under thresholding), and include corrections from inhibitor accumulation and explicit dependence on the inhibitor-side parameters. This is the subject of future theoretical work.

## Supplementary Note 3

### Dimensionless Parameterization and Fixed Point Structure for Numerical Simulations

This note serves as a prerequisite for the numerical simulations. JAPI (juxtacrine-activator paracrine-inhibitor) and PAPI (paracrine-activator paracrine-inhibitor) are the two reaction-diffusion architectures compared throughout this paper; full definitions are given in Supplementary Note 1. Here we reduce the dimensional JAPI system to a minimal set of dimensionless groups, establish the fixed point structure in that parameter space, and define the parameter regimes explored in the numerical results. The PAPI nondimensionalization is given alongside for parallel reference. The linear stability analysis of Supplementary Note 1 is formulated in dimensional parameters and does not require this reduction; however, all numerical simulations are reported in the dimensionless units defined here.

#### 1 Nondimensionalization

##### Dimensional JAPI system

In JAPI the activator is membrane-tethered, while the inhibitor is a diffusible field. Cells respond only to ligand presented on adjacent cells, not their own, so the autoactivation is non-cell-autonomous. The activator on cell *j* is induced only by activator-ligand presented on its *neighboring* cells through synNotch-ligand contact. Both species are produced by the same cell-autonomous response function *f*, evaluated on the signal received from neighbors and on the local inhibitor. With cells indexed by *j* on a lattice, the dimensional dynamics are:

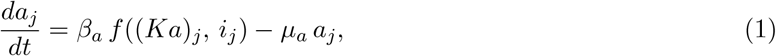

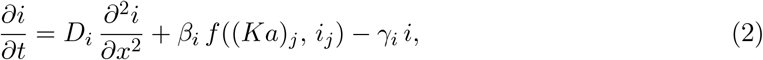

where *i*_*j*_ ≡ *i*(*x*_*j*_, *t*), the Hill response function is (Alon, 2007)

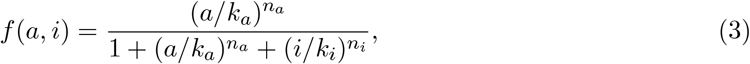

and the juxtacrine kernel sums activator over neighbors with no self-sensing:

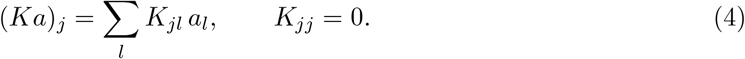

Writing the kernel as a function of the integer cell separation *r* = *l* − *j* (the coupling is translation-invariant on the lattice, so *K*_*jl*_ = *K*(*r*)), symmetric nearest-neighbor coupling on a 1D lattice gives *K*(±1) = 1*/*2 and *K*(*r*) = 0 otherwise; each cell receives the average of its two neighbors. The support of *K* sets the spatial range over which the activator can act: the nearest-neighbor choice fixes this activator interaction range to a single cell spacing by design, so it does not introduce a length scale competing with the inhibitor diffusion length. The total coupling weight is *κ* ≡ ∑_*l*_ *K*_*jl*_ = 1, so that homogeneous activator levels propagate unchanged through the kernel: (*Ka*)_*j*_ = *κ a*_0_ = *a*_0_ at any homogeneous state. Spatial position on the lattice is measured in units of cell-cell spacing, so the dimensionless lattice spacing is one by convention; the Fourier kernel eigenvalue 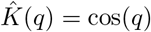 (Supplementary Note 1) follows from this convention, with *q* measured in radians per cell. Equations (1)–(2) write the activator on the discrete lattice and the inhibitor as a continuous diffusive field for analytical convenience and to match the formulation of the linear stability analysis; in numerical simulations both species are implemented on the same lattice with a discrete Laplacian for the inhibitor.

The dimensional system has seven parameters with units (*β*_*a*_, *β*_*i*_, *µ*_*a*_, *γ*_*i*_, *D*_*i*_, *k*_*a*_, *k*_*i*_), plus two already-dimensionless Hill exponents (*n*_*a*_ and *n*_*i*_).

##### Choice of scales

We rescale using the natural scales of the system (Murray, 2003):

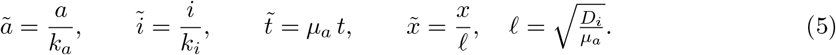

The time scale 1*/µ*_*a*_ is set by the activator’s degradation rate. The two concentration scales *k*_*a*_ and *k*_*i*_ are the Hill thresholds for activation and inhibition. The characteristic length 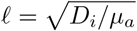 is the distance an inhibitor molecule diffuses on the activator time scale; it is *not* the same as the dimensional inhibitor diffusion length 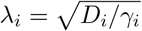, which represents the distance an inhibitor diffuses in its own lifetime. The latter appears in dimensionless units as 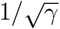 (Section 3).

##### Substitution: activator equation

Substituting (5) into (1) and dividing through by *k*_*a*_*µ*_*a*_:

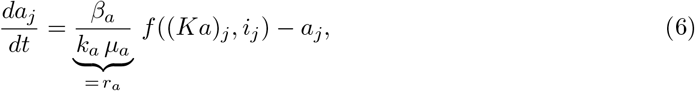

where tildes have been dropped: *a* and *i* now denote the dimensionless variables of (5), with the Hill thresholds *k*_*a*_ and *k*_*i*_ absorbed into their definition (equivalently, *k*_*a*_ = *k*_*i*_ = 1 in these units). The response function (3) accordingly reduces to

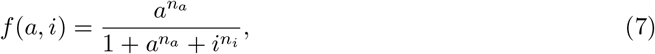

which we use throughout the remainder of the note. The kernel weights *K*_*jl*_ are dimensionless lattice quantities set by geometry and so are unchanged by the rescaling.

##### Substitution: inhibitor equation

Substituting (5) into (2), the diffusion coefficient picks up a factor *D*_*i*_*/*(*µ*_*a*_ *ℓ*^2^):

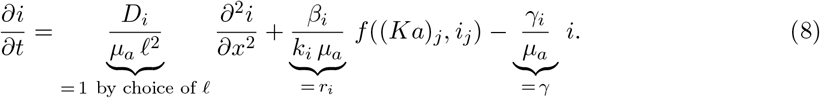

The choice 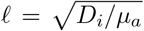 makes *D*_*i*_*/*(*µ*_*a*_*ℓ*^2^) = 1 exactly. This is what is meant by “*D*_*i*_ has been absorbed into the length scale”: *D*_*i*_ does not appear as a free parameter in the dimensionless equations because it has been used up defining *ℓ*; the entire inhibitor diffusion strength now lives in the choice of length unit.

##### Dimensionless JAPI system

Collecting:

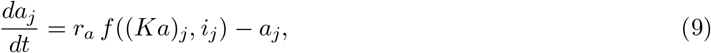

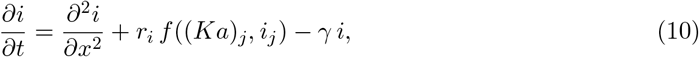

with five dimensionless groups:

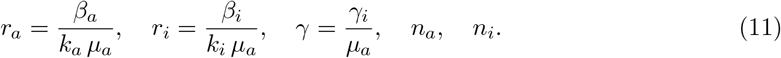

*r*_*a*_ is the dimensionless activator drive (the production-to-degradation ratio); *r*_*i*_ is the analogous quantity for the inhibitor; *γ* is the ratio of inhibitor to activator degradation rates, governing how quickly the inhibitor decays relative to the activator; *n*_*a*_ and *n*_*i*_ govern the steepness of the activation and inhibition responses. The activator diffusion coefficient *D*_*a*_ does not appear: it has been eliminated by the JAPI architecture itself, where the activator is membrane-bound rather than diffusible (Supplementary Note 1).

##### Comparison with PAPI

In PAPI, the activator is also a diffusible field, with diffusion coefficient *D*_*a*_. The dimensional inventory adds *D*_*a*_, and the activator equation gains a Laplacian term. Applying the same rescaling (which uses 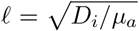 as the unit of length in both architectures) yields the dimensionless PAPI system:

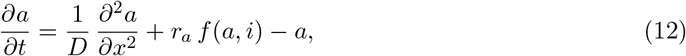

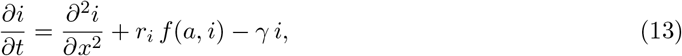

with one additional dimensionless group:

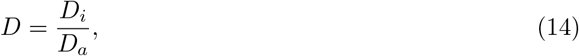

the inhibitor-to-activator diffusion ratio. We adopt the convention *D* = *D*_*i*_*/D*_*a*_ (rather than its reciprocal) so that the Turing-favorable regime corresponds to *D >* 1, consistent with classical reaction-diffusion literature. PAPI therefore has six dimensionless groups: *r*_*a*_, *r*_*i*_, *γ, D, n*_*a*_, *n*_*i*_. The same rescaling absorbs *D*_*i*_ into the length scale in both architectures; the parameter *D*_*a*_ that distinguishes PAPI cannot be absorbed into the same length scale and survives as the residual ratio *D* = *D*_*i*_*/D*_*a*_.

#### 2 Steady State Analysis in Dimensionless Form

Setting spatial gradients to zero, the dimensionless steady states satisfy:

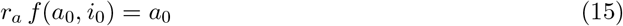

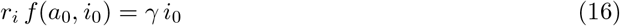

##### Trivial fixed point

(*a*_0_, *i*_0_) = (0, 0) always satisfies (15), (16). For *n*_*a*_ *>* 1, the activator production 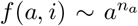 vanishes faster than linearly near the origin, so the linearized dynamics carry no production term and the trivial fixed point is linearly stable; this state therefore does not support Turing instability in the regimes considered here (Supplementary Note 1). The *n*_*a*_ = 1 case is parameter-dependent and is not considered.

##### Reduction to a scalar root-finding problem

Dividing (15) by (16) gives

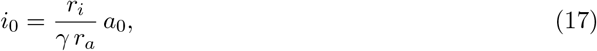

so any nontrivial fixed point lies on this line in the (*a*_0_, *i*_0_) plane. The two steady-state equations therefore reduce to a scalar root-finding problem along (17); the dynamics themselves remain two-dimensional.

##### Nontrivial fixed points

Substituting (17) into (15) and dividing through by the overall factor of *a* (which separates the trivial root *a*_0_ = 0 already accounted for above), the nontrivial fixed points are the positive roots of

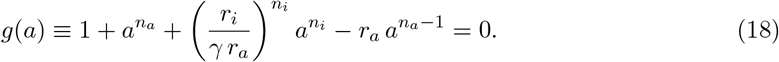

Note that *g*(0) = 1 ≠ 0: the trivial root lives in the full equation *a* · *g*(*a*) = 0, not in *g* itself. For integer Hill exponents *n*_*a*_, *n*_*i*_, Descartes’ rule of signs admits 0, 1 (a double root), or 2 positive real roots, corresponding to the monostable, saddle-node, and bistable cases respectively; for non-integer Hill exponents the same count holds by a generalized argument on the sign changes of *g*(*a*).

##### Saddle-node boundary

In the (*r*_*a*_, *r*_*i*_*/γ*) parameter plane (at fixed *n*_*a*_, *n*_*i*_), the boundary between monostable and bistable regimes is the set on which (18) admits a positive double root, i.e. *g*(*a*) = 0 and *g*^′^(*a*) = 0 hold simultaneously. We solve this pair numerically for each (*n*_*a*_, *n*_*i*_) to map the bistable region used in the simulations.

##### Activator-only limit

In the limit *r*_*i*_ → 0, (18) reduces to 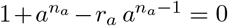, and the saddle-node tangency between the production curve 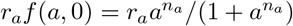 and the degradation line *a* (Strogatz, 1994) gives the closed-form thresholds:

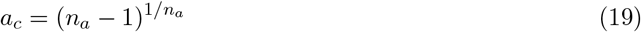

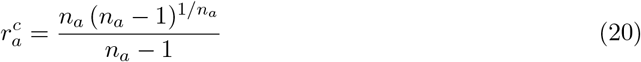

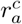 is therefore the saddle-node boundary in the activator-only slice *r*_*i*_ = 0, and serves as a lower-bound calibration for the full inhibited boundary: for *r*_*i*_ *>* 0, the saddle-node boundary lies above 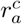 in the (*r*_*a*_, *r*_*i*_*/γ*) plane. The critical value 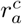 decreases monotonically toward 1 as *n*_*a*_ increases, so higher cooperativity makes bistability easier to achieve in this limit.

##### Fixed-point structure in the bistable region

In the bistable region of the inhibited (*r*_*a*_, *r*_*i*_*/γ*) plane, the system has three fixed points along the line (17): the stable off state at (0, 0), an unstable intermediate fixed point at the smaller positive root of *g*, and a stable activated state at the larger positive root, with *a*_0_ → *r*_*a*_ and *i*_0_ → *r*_*i*_*/γ* for *r*_*a*_ well above the boundary. Stability classifications follow from the standard 2×2 Jacobian analysis of (15)–(16) (Strogatz, 1994) and are not reproduced here.

##### Effect of inhibition

Increasing *r*_*i*_*/γ* shifts the saddle-node boundary toward larger *r*_*a*_, shrinking the bistable region. Mechanically, the inhibitor suppresses the production curve at any nontrivial fixed point relative to the activator-only case (Gierer & Meinhardt, 1972), raising the value of the unstable intermediate fixed point in *a* and shifting the separatrix between the two basins of attraction toward the activated state.

#### 3 Parameter Space for Numerical Simulations

The five dimensionless groups (11) define the parameter space explored numerically; for PAPI, the additional group *D* = *D*_*i*_*/D*_*a*_ is included, fixed at *D* = 10 across the scan (a standard Turing-favorable regime in classical reaction-diffusion literature). Simulations span both the monostable and bistable regimes, separated by the saddle-node boundary in the (*r*_*a*_, *r*_*i*_*/γ*) plane (Section 2), which reduces to 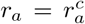 in the activator-only limit *r*_*i*_ → 0. The scan is systematic across the full parameter space (cf. Marcon et al., 2016); visualizations of patterning outcomes are necessarily lower-dimensional projections, with two or three of the groups held fixed to render a 2D plane or 3D volume. The choice of which dimensions to display is dictated by visualization rather than by physics, and the held-fixed values are reported alongside the corresponding figure. Full numerical setup (lattice dimensions, boundary conditions, initial-condition protocol, noise amplitude, time horizon, solver and tolerances, and pattern-classification criteria) is specified in Methods. JAPI and PAPI share the same homogeneous fixed-point structure (Section 2) but differ in their spatial dispersion: in JAPI the activator coupling enters multiplicatively as 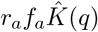, while in PAPI it enters additively as −(1*/D*) Λ(*q*). At small *q* the JAPI cosine kernel can be locally matched to an effective activator diffusion coefficient 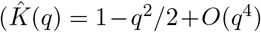, equivalent to a continuum diffusion 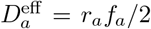 at long wavelengths; Supplementary Note 1), but the full dispersion relations and nonlinear couplings are not equivalent at any single value of *D*, and the two systems are scanned independently.

Two distinct lengths appear in this formulation: the rescaling length 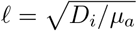 used to define dimensionless space (Section 1), and the dimensional inhibitor diffusion length 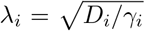, the distance an inhibitor molecule diffuses in its own lifetime. In dimensionless units, 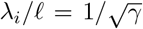, which sets the natural spatial scale of domain size in the irregular patterning regime (Supplementary Note 2). All spatial outputs (domain widths, inter-feature spacing, pattern wavelengths) are reported in dimensionless units, i.e. in multiples of *ℓ*.

## Supplementary Note 4

### Domain Arrest in the Irregular Patterning Regime of Reaction-Diffusion Systems

This note addresses a question raised by the experimental observation that JAPI circuits implemented in mammalian cells produce robust patterns despite operating in a parameter regime where linear stability analysis classifies the homogeneous activated state as stable. The same regime is accessible numerically in both JAPI and PAPI architectures and gives rise to non-periodic, irregular patterns formed of finite-sized activated domains distributed in space (see main text Fig. 2D for example). This note provides a heuristic backbone of an analytical mechanism by which this occurs.

The analysis is general to reaction-diffusion systems with a diffusing inhibitor and applies to both JAPI and PAPI architectures: the inhibitor-field calculation requires only *β*_*i*_, *D*_*i*_, *γ*_*i*_, and the activation threshold *I*_*c*_, not the form of the activator spatial coupling. Architecture-specific content enters the result in two places: (i) through the activator-side parameters that set *I*_*c*_ (the inhibitor concentration that prevents activation at the front, which depends on the receiver cell’s production rate, degradation rate, and Hill response), and (ii) through the activator front velocity *v*, which determines whether the quasi-static approximation underlying the closed-form result is valid. Within the quasi-static regime (defined below), the closed-form arrested domain size depends on (*λ*_*i*_, *β*_*i*_, *I*_*c*_) only; *v* does not appear at leading order. Throughout this note, we treat the activator front velocity as an abstract quantity *v* representing the rate at which the activated domain expands. In the JAPI architecture, *v* reaches its maximum value, *v*_*a*_ (Supplementary Note 2), in the inhibitor-free limit; during pattern formation *v* is bounded above by *v*_*a*_ and decreases as inhibitor accumulates around the expanding domain.

The analysis is restricted to the irregular regime, where individual domains form and stabilize in isolation. The irregular regime arises from both monostable and bistable parameter conditions in JAPI/PAPI architectures; we comment on the implications for stability at the end of the existence-condition section. It does not apply to the Turing regime, where domain boundaries are set by the dispersion relation rather than by self-consistent inhibitor accumulation.

#### Setup: an isolated activated domain in steady state

In the irregular regime, patterns form by nucleation: local fluctuations cross a threshold, an activated domain appears, expands by relay or diffusion of the activator, and produces inhibitor that accumulates around it. We analyze a single such domain treated in isolation, assuming it has reached a steady state in which:

- The domain has half-width *R* and is bounded by an activation front
- Inside the domain, the activator is at its activated steady state, so all cells within the domain produce inhibitor at the maximum rate *β*_*i*_
- Outside the domain, no cells are activated and no inhibitor is produced locally
- The inhibitor field is treated as quasi-static, instantly tracking the slowly evolving domain geometry

We treat the inhibitor field as quasi-static, instantly tracking the slowly evolving domain geometry. This corresponds to the slow-front limit 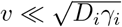, where the inhibitor relaxation time 1*/γ*_*i*_ is much shorter than the timescale on which the activation front advances by one diffusion length. This is a simplifying limit adopted to enable the closed-form solution presented below; the validity of this assumption for the experimental system, the corrections at finite *v*, and the breakdown of the quasi-static framework at high *v* are addressed in the final section of this note.

We additionally treat the activation front as a locally planar one-dimensional boundary. This approximation is exact for stripe-like domains and accurate for compact two- or three-dimensional domains when the boundary curvature radius is large compared to the inhibitor diffusion length *λ*_*i*_. For compact two-dimensional domains with *R* ∼ *λ*_*i*_, curvature corrections modify the boundary inhibitor concentration; the leading-order result of the planar calculation captures the qualitative dependence on *β*_*i*_, *λ*_*i*_, and *I*_*c*_, while quantitative agreement requires the full Green’s-function solution. We adopt the planar approximation for transparency and analytical tractability; numerical simulations in the main text (Fig. 2L-M) do not rely on it.

In one dimension, treating the domain interior as a region of constant inhibitor production and the exterior as a region with no production, the inhibitor concentration *I*(*x*) satisfies the linear equation:

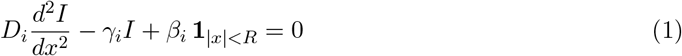

where **1**_|*x*|*<R*_ is the indicator function of the domain. The natural length scale is the inhibitor diffusion length:

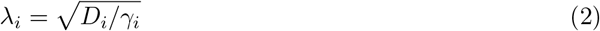

the characteristic distance over which the inhibitory halo around the domain decays. The parameters used throughout this analysis are summarized in Table 1.

**Table 1.**
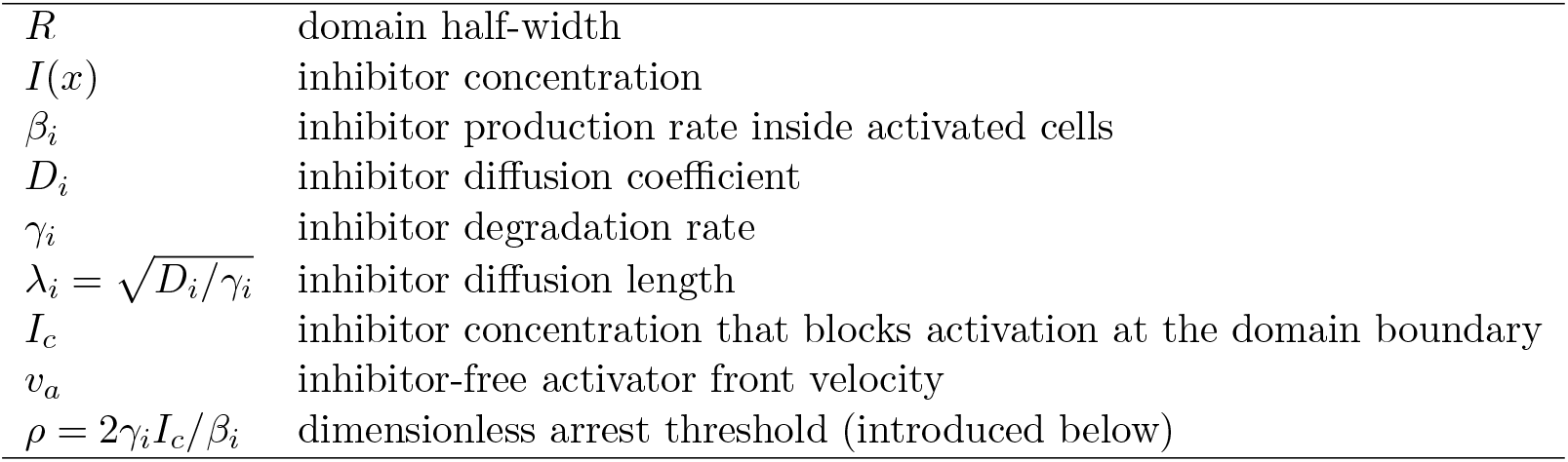
Parameters used in the arrest analysis.

#### Solving for the boundary inhibitor concentration

Equation (1) is solvable exactly. By symmetry the solution is even in *x*. Inside the domain, the general solution combines a particular solution *I*_*p*_ = *β*_*i*_*/γ*_*i*_ with the even homogeneous mode cosh(*x/λ*_*i*_). Outside the domain, the solution decays exponentially as 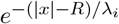 to satisfy *I*(*x*) → 0 at infinity. Matching *I* and *I*^′^ at *x* = *R* and solving the resulting linear system gives:

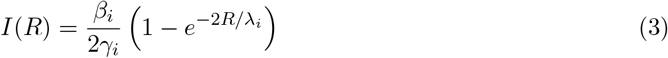

The boundary value has two clean limits. For small domains (*R* ≪ *λ*_*i*_), Taylor expansion gives *I*(*R*) ≈ (*β*_*i*_*/γ*_*i*_) · (*R/λ*_*i*_), so the boundary inhibitor grows linearly with domain size, since the entire domain contributes to the inhibitor field at the edge. For large domains (*R* ≫ *λ*_*i*_), the exponential vanishes and *I*(*R*) → *β*_*i*_*/*(2*γ*_*i*_), so the boundary value saturates, because cells more than a few *λ*_*i*_ away from the boundary do not contribute to it. The factor of one-half reflects the symmetric leakage of inhibitor outward from the boundary.

#### The arrest condition

The expanding front stalls when the inhibitor concentration at the boundary reaches a critical threshold *I*_*c*_ sufficient to prevent further activation: *I*(*R*) = *I*_*c*_. *I*_*c*_ is itself a derived quantity that depends on the activator-side parameters governing the receiver cell’s response (production rate, degradation rate, Hill parameters of activation): the inhibitor level at which the receiver fails to activate is set by the balance between inhibition and the activator signal arriving from the neighbor. Through this dependence, activator-side parameters enter the arrest condition (Eq. 6) by setting *I*_*c*_, separately from the velocity channel discussed in the regime structure section below. This note takes *I*_*c*_ as an empirical input to the arrest analysis; deriving *I*_*c*_ from circuit parameters is part of the future theoretical work. Substituting (3) gives the arrest condition:

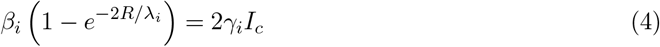

which can be written compactly as *β*_*i*_*h*(*R/λ*_*i*_) = *γ*_*i*_*I*_*c*_ with the dimensionless geometric factor *h*(*x*) = (1 − *e*^−2*x*^)*/*2. We note that nucleation, the formation of the initial activated domain that this analysis assumes as its starting point, requires perturbations above a finite threshold, and is documented numerically in main text Fig. S7C-D rather than analytically here.

#### Existence condition for arrest

Equation (4) has an immediate consequence. Since 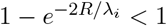 for any finite *R*, arrest is only possible when:

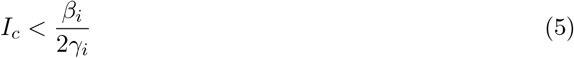

If the activation threshold *I*_*c*_ exceeds half the maximum interior inhibitor concentration *β*_*i*_*/γ*_*i*_, the boundary inhibitor can never reach *I*_*c*_ regardless of how large the domain grows. The front never stalls and the system proceeds to uniform activation. This existence condition therefore defines a quantitative boundary in parameter space between conditions where isolated domains arrest at finite size and conditions where activation propagates indefinitely until the system reaches uniform activation.

We note that equation (6), derived in the next section, identifies the half-width at which the arrest condition is satisfied; stability of the arrested state is addressed there.

#### Domain size when arrest occurs

When (5) is satisfied, solving (4) explicitly for *R*:

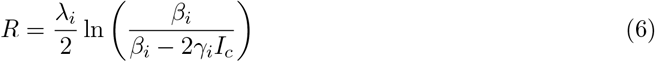

In dimensionless form, defining *ρ* = 2*γ*_*i*_*I*_*c*_*/β*_*i*_ as the dimensionless arrest threshold (satisfying 0 *< ρ <* 1 when the existence condition holds), Eq. (6) becomes

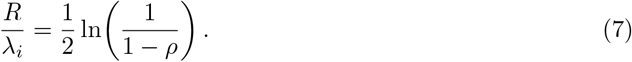

This separates the length scale *λ*_*i*_ from the dimensionless arrest threshold *ρ*, and makes transparent that the existence-condition divergence (*R* → ∞ as *ρ* → 1^−^) corresponds to approaching the boundary *I*_*c*_ → *β*_*i*_*/*(2*γ*_*i*_) from below.

The qualitative dependencies on independent parameters follow directly: increasing *D*_*i*_ at fixed *β*_*i*_, *γ*_*i*_, *I*_*c*_ increases *λ*_*i*_ while leaving *ρ* unchanged, so *R* increases proportionally. Increasing *β*_*i*_ at fixed *D*_*i*_, *γ*_*i*_, *I*_*c*_ decreases *ρ*, so *R* decreases. Increasing *I*_*c*_ at fixed *β*_*i*_, *γ*_*i*_, *D*_*i*_ increases *ρ*, so *R* increases. The effect of changing *γ*_*i*_ in isolation is ambiguous because *γ*_*i*_ enters both *λ*_*i*_ and *ρ*; the dimensionless form clarifies that the net effect depends on which other quantities are held fixed.

##### Stability of the arrested state

Equation (6) identifies the half-width at which the arrest condition *I*(*R*) = *I*_*c*_ is satisfied; it does not by itself establish dynamical stability of the arrested state. Local stability requires a boundary-motion law (e.g., *dR/dt* = *V* (*I*_*c*_ − *I*(*R*)) with *V* (0) = 0 and *V* ^′^(0) *>* 0) and analysis of small perturbations to *R*. Because *dI/dR >* 0 in the planar quasi-static calculation, the geometric condition for dynamical stability is satisfied: a small increase in *R* raises the boundary inhibitor above *I*_*c*_, suppressing further expansion; a small decrease lowers it below *I*_*c*_, releasing the front. A formal stability analysis with an explicit front-motion law is the subject of future theoretical work.

Empirically, arrested domains persist in numerical simulations across the parameter ranges examined here (main text Fig. 2 and Fig. S7), consistent with dynamical stability in both the bistable and monostable regimes accessible to the system. In the bistable case, the arrested configuration sits between two homogeneous steady states; in the monostable case, persistence of arrested domains arises despite the absence of an alternative stable homogeneous state, distinguishing the irregular regime from a simple bistable picture.

#### Activator front velocity and the regime structure of arrest

The closed-form arrest condition derived above (Eq. 6) makes no reference to the activator front velocity *v*: *R* is determined entirely by inhibitor parameters and the activation threshold. In the quasi-static limit, the arrested domain size follows from the geometric self-consistency condition *I*(*R*) = *I*_*c*_, which involves only the inhibitor field; the role of *v* is to set how quickly arrest is reached, not what size it stalls at.

This *v*-independence holds only in the quasi-static regime, 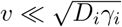. Because *v* varies in time during expansion (reaching its maximum *v*_*a*_ in the inhibitor-free limit at the start of the front’s advance, and approaching zero near arrest), we use *v*_*a*_ as the relevant *v* in regime comparisons: it is the maximum value *v* takes during expansion, and serves as a conservative upper bound. If 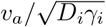 places the system in regime (a), the actual 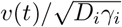 does so at all times during expansion. Three regimes can be distinguished:

- **(a) Quasi-static**, 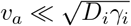: Equation (6) applies at leading order; *R* does not depend on *v*. A heuristic order-of-magnitude estimate for the subleading correction is Δ*R* ∼ *v*_*a*_*/γ*_*i*_, valid in the bulk of regime (a) where *dI/dR* remains of order *β*_*i*_*/*(*γ*_*i*_*λ*_*i*_). This estimate breaks down near the existence boundary (*ρ* → 1^−^), where *dI/dR* vanishes and small finite-velocity reductions in boundary inhibitor can produce disproportionately large corrections to *R*. A rigorous moving-front derivation including the velocity-dependent boundary inhibitor concentration is the subject of future theoretical work.
- **(b) Lagging**, 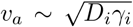: The boundary inhibitor field is substantially below its quasi-static value at any given *R. R* remains finite but is larger than equation (6) predicts and depends on *v* in a nontrivial nonlinear way.
- **(c) Runaway**, 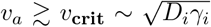 : The boundary inhibitor saturates at a value below *I*_*c*_, the arrest condition is never satisfied, and the domain expands without bound until external constraints (tissue size, depletion of resources) intervene.

The transition from (b) to (c) is set by a critical velocity *v*_crit_ at which the dynamically saturated boundary inhibitor falls below *I*_*c*_. Determining *v*_crit_ explicitly, and the full functional form *R*(*v*) in regime (b), requires a traveling-wave analysis and is left to future theoretical work.

In regime (a) the dependence of *R* on inhibitor parameters takes a simple form: *R* increases monotonically with *λ*_*i*_ and decreases monotonically with *β*_*i*_. The dependence on *v* is regime-specific: *R* is approximately *v*-independent at leading order in regime (a), depends nontrivially on *v* in regime (b), and is undefined in regime (c).

##### Empirical placement of JAPI

For the JAPI experimental system, the propagation velocity in L929 fibroblast transceivers (a configuration that lacks the inhibitor branch and therefore reports *v*_*a*_ directly) is *v*_*a*_ ∼ 0.13 mm/day (Santorelli et al., 2024), equivalent to ∼ 1.5 × 10^−3^ *µ*m/s. Using inhibitor parameters consistent with engineered secreted morphogens (diffusion *D*_*i*_ ∼ 10 *µ*m^2^/s, half-life *t*_1*/*2_ ∼ 2–6 hours, comparable to measured values for canonical Lefty in zebrafish; Müller et al., 2012), we estimate the inhibitor degradation rate as *γ*_*i*_ = ln(2)*/t*_1*/*2_ ≈ 3.2 × 10^−5^ to 9.6 × 10^−5^ s^−1^, giving 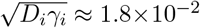 to 3.1×10^−2^ *µ*m/s. The dimensionless ratio 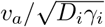 therefore falls in the range ∼ 0.05–0.08, placing the system in regime (a): equation (6) applies as the leading-order description and arrest occurs robustly. Finite-velocity corrections are bounded in the bulk of regime (a): the heuristic lag scale *v*_*a*_*/γ*_*i*_ evaluates to ∼ 16–47 *µ*m, a fraction of the domain half-width *R* ∼ 50–100 *µ*m (the half-width corresponding to arrested domain diameters of ∼ 100–200 *µ*m), suggesting bulk corrections to *R* of order ∼ 30–50% when evaluated with the conservative upper-bound velocity *v*_*a*_. The actual correction is likely smaller still, since the front velocity near arrest is well below *v*_*a*_; corrections near the existence boundary *ρ* → 1 require the moving-front analysis deferred to future theoretical work. These are order-of-magnitude estimates; precise placement would require direct measurement of the diffusion coefficient and degradation rate of the specific inhibitor used in the JAPI implementation. The architectural distinction between JAPI and PAPI enters here through both *I*_*c*_ (set by activator-side parameters of each architecture) and *v*_*a*_: in PAPI, *v*_*a*_ scales as the square root of the activator diffusion coefficient times the activator’s linear growth rate (the Fisher-KPP front scaling) and therefore depends on *D*_*a*_, while in JAPI, *D*_*a*_ is absent and *v*_*a*_ is set by the parameters of Supplementary Note 2.

#### Biological summary

The arrest mechanism analyzed here gives a simple picture of what controls domain size in the irregular regime of reaction-diffusion systems:

- **Stronger inhibitor production** (higher *β*_*i*_) −→ boundary inhibitor reaches *I*_*c*_ at smaller *R* −→ smaller domains.
- **Wider inhibitor reach** (higher *λ*_*i*_ at fixed dimensionless threshold *ρ* = 2*γ*_*i*_*I*_*c*_*/β*_*i*_) −→ inhibitor halo extends further −→ larger domains. Increasing *D*_*i*_ at fixed *β*_*i*_, *γ*_*i*_, *I*_*c*_ increases *R* proportionally to *λ*_*i*_; the effect of changing *γ*_*i*_ in isolation depends on which other quantities are held fixed, since *γ*_*i*_ enters both *λ*_*i*_ and the dimensionless threshold *ρ*.
- **Front velocity** affects domain size in a regime-specific way: *R* is approximately independent of *v* in the quasi-static regime 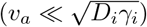, depends nontrivially on *v* when *v*_*a*_ is comparable to 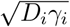, and arrest fails entirely when *v*_*a*_ exceeds a critical value. The relevant comparison uses *v*_*a*_, the inhibitor-free upper bound on the front velocity; the actual front velocity during expansion is bounded above by *v*_*a*_ and approaches zero near arrest.
- **Arrest is only possible when** *I*_*c*_ *< β*_*i*_*/*(2*γ*_*i*_). When this condition fails, the front cannot be stalled and the system proceeds to uniform activation.

This analysis applies to both JAPI and PAPI architectures, with architecture entering through two channels: *I*_*c*_ (set by the activator-side parameters governing the receiver cell’s response) and *v*_*a*_ (set by how the activator’s spatial coupling generates a front velocity). The inhibitor-field calculation itself is architecture-independent. Within regime (a), the closed-form result for *R* (Eq. 6) depends on (*λ*_*i*_, *β*_*i*_, *I*_*c*_); *v*_*a*_ does not appear at leading order, but determines regime placement. A formal derivation of *v*_*a*_ in terms of activator-side circuit parameters, together with the activator-side derivation of *I*_*c*_ and corrections from inhibitor coupling, is the subject of future theoretical work. In JAPI, *v*_*a*_ is set by a smaller number of parameters since *D*_*a*_ is absent, making domain size control more parsimonious through this channel.

